# Beyond forest succession: a gap model to study ecosystem functioning and tree community composition under climate change

**DOI:** 10.1101/2020.06.10.140616

**Authors:** Xavier Morin, François de Coligny, Nicolas Martin-StPaul, Harald Bugmann, Maxime Cailleret, Jean-Marc Limousin, Jean-Marc Ourcival, Bernard Prevosto, Guillaume Simioni, Michel Vennetier, Joannès Guillemot

## Abstract

Climate change impacts forest functioning and dynamics, and large uncertainties remain regarding the interactions between species composition, demographic processes, and environmental drivers. There are few robust tools available to link these processes, which precludes accurate projections and recommendations for long-term forest management. Forest gap-models present a balance between complexity and generality and are widely used in predictive forest ecology. However, their relevance to tackle questions about the links between species composition, climate and forest functioning is unclear. In this regard, demonstrating the ability of gap-models to predict the growth of forest stands at the annual time scale – representing a sensitive and integrated signal of tree functioning and mortality risk - appears as a fundamental step.

In this study, we aimed at assessing the ability of a gap-model to accurately predict forest growth in the short-term and potential community composition in the long-term, across a wide range of species and environmental conditions. To do so, we present the gap-model ForCEEPS, calibrated using an original parameterization procedure for the main tree species in France. ForCEEPS was shown to satisfactorily predict forest annual growth (averaged over a few years) at the plot level from mountain to Mediterranean climates, regardless the species. Such an accuracy was not gained at the cost of losing precision for long-term predictions, as the model showed a strong ability to predict potential community composition along a gradient of sites with contrasted conditions. The mechanistic relevance of ForCEEPS parameterization was explored by showing the congruence between the values of key model parameter and species functional traits. We further showed that accounting for the spatial configuration of crowns within forest stands, the effects of climatic constraints and the variability of shade tolerances in the species community are all crucial to better predict short-term productivity with gap-models.

The dual ability of predicting short-term functioning and long-term community composition, as well as the balance between generality and realism (i.e., predicting accuracy) of the new generation of gap-models may open great perspectives for the exploration of the biodiversity-ecosystem functioning relationships, species coexistence mechanisms, and the impacts of climate change on forest ecosystems.

## INTRODUCTION

Forests cover about 30% of the land at the global scale, harbor most of terrestrial biodiversity, are an important carbon sink (Pan et al. 2011), play a pivotal role for climate regulation (Chapin et al. 2008) and provide key ecosystem services to humans (TEEB 2010). However, climate change puts forests at high risk, including disruption in forest dynamics (McDowell et al. 2020), as harsher environmental conditions strongly impact forest structure and composition (Esquivel-Muelbert et al. 2019) and functioning (Boisvenue and Running 2006, Allen et al. 2010, Lindner et al. 2010). In turn, compositional changes have been shown to affect forest functioning (Nadrowski et al. 2010, Liang et al. 2016), in interaction with climatic drivers (Coomes et al. 2014, Jactel et al. 2018). Yet, we lack robust tools to explore the interactive effects of biodiversity and climate change on forest dynamics and functioning.

Trees are long-lived organisms, which complicates the implementation of experiments designed to assess the influences of future environmental conditions (e.g., increased atmospheric CO_2_ (Korner et al. 2005) or water stress (Limousin et al. 2009)) and community composition (including species richness, Castagneyrol et al. 2013, Verheyen et al. 2016) on forest ecosystem functioning. While such experiments are key to study forest ecosystems, they require years to yield relevant results, which are necessarily conditioned by specific site conditions, thereby limiting their generality (Nadrowski et al. 2010, Norby and Zak 2011). An alternative approach lies in the design of field sampling along climate and/or diversity gradients, which has yielded significant results in the last years (e.g., Jucker et al. 2016, del Río et al. 2017, Jourdan et al. 2019), although it can be affected by confounding factors.

Complementing these approaches, forest models represent a crucial tool to explore the interactions and feedbacks among species composition, forest functioning and climate (Cordonnier et al. 2018b). Yet, the term " forest models " covers a wide range of approaches, as recently reviewed (Pretzsch et al. 2015, Ruiz-Benito et al. 2020). Forest models were indeed used to predicting forest functioning and growth at scales ranging from tree, to stand (Makela et al. 2000) and landscape (Pacala et al. 1993). Moreover forest models differ in their complexity, from empirical yield tables (Skovsgaard and Vanclay 2008) to ecophysiology-based models (Dufrêne et al. 2005, Simioni et al. 2016) that explicitly describe part of the biological mechanisms at stake but require a large amount of data to be properly calibrated and forced. By contrast, forest gap models (hereafter referred to as “gap models”), operating mostly at the stand scale, rely on empirical relationships, physiological knowledge and first principles from ecological theory (Bugmann 2001). Because these models incorporate physically-or ecologically-based hypotheses while relying on a small set of species-specific parameters, we believe that they are good candidates to explore forest responses to future growing conditions across spatial scales.

The design of gap model was originally motivated by the recognition that canopy gaps created by falling trees are a key driver shaping forest structure, dynamics, and succession (Botkin et al. 1972). Although gap models also incorporate representations of abiotic constraints (e.g., water or nutrient stress) on forest functioning, and in some instances competition for belowground resources, their key feature is a representation of the ability of trees of contrasted sizes and different species to compete for light resource. Gap models have been originally developed to understand the processes at play during forest succession (Botkin et al. 1972, Canham et al. 1994, Bugmann 2001). Consequently, they are commonly validated against potential natural vegetation (hereafter “PNV”), or against standing biomass accumulated over long (>50 years) time periods at the tree or stand level (Bugmann 1996, Strigul et al. 2008, Didion et al. 2009, Rasche et al., 2011). Gap models have been used to address a variety of basic and applied research questions, including the effects of climate on forest biomass and composition (e.g., Pfister and Bugmann 2000) or forest management planning (e.g. Rasche et al. 2011, Mina et al. 2017).

Recent developments have shown that gap models can be further used to explore species coexistence mechanisms (Chauvet et al. 2017), diversity effects on the functioning of forest ecosystems (Morin et al. 2011, Bohn and Huth 2017) and their response to climate change (Morin et al. 2018). These new perspectives highlight the importance of forest structure and light-related interactions for forest functioning. In fact, forest structure has been shown to influence forest growth (Hardiman et al. 2011, Gough et al. 2019) and to partly mediate tree diversity effects on productivity (Dănescu et al. 2016, Cordonnier et al. 2019, Schnabel et al. 2019). Enhanced canopy space occupation (‘canopy packing’, Jucker et al. 2015) and light capture, which is mediated by the coexistence of species with contrasting shade tolerance, was shown to be key in the functioning of diverse and structurally complex forests (Williams et al. 2017). The presence of shade-tolerant species in tree species mixtures indeed strongly modulates the way tree diversity affects forest functioning and productivity (Toïgo et al. 2018, Van de Peer et al. 2018, Cordonnier et al. 2018a). Gap models can be parameterized for a wide range of species and environmental conditions, and could thus be a crucial tool to explore how differences in shade-tolerance affect the relationships between species richness and forest functioning (Morin et al. 2011, Toïgo et al. 2018). However, the multi-dimensional configuration of crowns in forest stands is not often represented explicitly in gap models (but see Maréchaux and Chave 2017, Pacala et al. 1993, Purves et al. 2008), which hinders the assessment of the importance of architectural plasticity and canopy packing on forest productivity, species succession and coexistence.

Moreover, exploring Biodiversity-Ecosystem Functioning (BEF) relationships or species coexistence under climate change using gap models will require to assess (i) whether they are able to predict key patterns linking forest composition and functioning and (ii) whether they embed a sound representation of the underlying mechanistic processes. Annual tree growth was shown to be a sensitive and integrated signal of tree functioning and mortality risk (Dobbertin 2005, IFN 2016, Cailleret et al. 2017, DeSoto et al. 2020), in contrast to PNV and standing biomass, which result from the accumulated effects of multiple ecological processes (e.g. tree recruitment, growth and mortality). Demonstrating the ability of gap models to predict the growth of forest stands at the annual time step or across a few years (i.e., to predict biomass *fluxes* in addition to biomass *stocks*; Guillemot et al., 2017), would open important research avenues to investigate how the mechanisms underlying BEF-relationships shape forest dynamics and community assembly (Cordonnier et al. 2018b). In addition, progress in trait-based ecophysiology has allowed identifying key functional traits involved in tree survival and growth in contrasting environments (Falster et al. 2018). Testing the congruence between key model parameters and functional traits is thus another way to evaluate the mechanistic relevance of these models.

Here, we aim to test whether a gap model can predict the annual growth of forests differing widely in species composition and climatic conditions throughout France, using only a small set of parameters that can be calibrated based on forest inventories. French mainland forests are found in a wide range of conditions including mountain, continental, oceanic and Mediterranean climates (Verkerk et al. 2019) and are therefore ideal to evaluate the generality of the hypotheses embedded in models. We present the ForCEEPS model (Forest Community Ecology and Ecosystem ProcesseS), derived from ForClim (Bugmann 1996, Didion et al. 2009). Among other novelties, ForCEEPS embeds an improved representation of tree-tree competition for light by considering individual crown sizes in the vertical canopy space. ForCEEPS was parameterized for the main French tree species, and evaluated against annual growth (averaged across a few years) at the tree and stand scale, and against PNV. In addition, we verified the mechanistic relevance of ForCEEPS by assessing the congruence of key species parameters with functional traits. Finally, we conducted a sensitivity analysis on the ForCEEPS stand growth predictions, to quantify the importance of 1) an explicit representation of crown size, 2) the variability of shade-tolerance among species, and 3) the climatic constraints for accurately simulating stand growth.

## MODEL DESCRIPTION

### Overview

The ForCEEPS model is a forest gap model (also called forest dynamics model). Forest gap models simulate abiotic (climate and soil properties) and biotic constraints (tree-tree competition for light) on tree establishment, growth, and survival in small parcels of land (“patches”). The mechanisms embedded in gap models rely on ecological hypotheses clearly stated, such as the trade-off between growth in full light and survival under shade (Bazzaz 1979). Tree height and crown dimensions are inferred from allometry, based on tree trunk diameter, which is also the main variable measured in forestry surveys. Gap models commonly simulate forest dynamics at an annual time step, and do not explicitly represent biogeochemical cycling. ForCEEPS shares many features with the JABOWA (Botkin et al. 1972) and ForCLIM models (Bugmann 1996), and more precisely with ForCLIM 2.9.6 (Didion et al. 2009). Below, we present the central principles of ForCEEPS and the key developments that differentiate it from other gap models (a full description of the model is provided in Appendix A).

The simulated patches are independent from each other, and properties at the forest level are obtained by aggregating the properties over all patches (Shugart 1984, Bugmann 2001). Within each patch (i.e., usually between 400 and 1000 m^2^), environmental conditions are assumed to be horizontally homogeneous. The spatial location of trees is therefore implicit, and the competitive ability of a tree is assumed equal for all trees of similar size and species. This hypothesis allows for several simplifications in the representation of tree-tree interactions, but imposes that the patch size cannot be larger than 1000 m^2^, which is assumed to be the maximum area influenced by a tree (Shugart 1984). Gap models are often cohort-based, assuming that all trees of the same species and age behave similarly, for the sake of simulation efficiency. By contrast, ForCEEPS is completely individual-based, which notably allows to simulate the intraspecific variability in competitive ability. Another novel aspect in ForCEEPS is the possibility of imposing a feedback between the actual forest composition and the identity of the colonizing seedlings each year. This latter feature may be crucial for examining mechanisms of species coexistence in tree communities (Cordonnier et al. 2018b). However, with regard to the objective of the present paper, the most crucial development of ForCEEPS in comparison with ForClim is the implementation of a new module for tree-tree competition for light, i.e. a key factor controlling growth and forest structure (Schwinning and Weiner 1998), where the individual crown lengths are explicitly represented in the vertical canopy space (see Appendix A).

Tree establishment, growth, and mortality are simulated at a yearly time step, but monthly climatic data (monthly mean temperature and precipitation sum) are used to estimate annual or seasonal degree-days sum (*GDD*), winter temperatures, and a drought index (*DrI*). The latter depends on monthly soil water content (*SWC*) that is calculated from a monthly water budget (Bugmann and Solomon 2000) and is influenced by the site-specific maximum soil water holding capacity. Last, soil nutrients content (*N*_*soil*_) is another abiotic factor simulated in ForCEEPS, considered constant at the site level (Appendix A).

### Seedling establishment

Seedlings are established with a diameter at breast height of 1 cm. Establishment success is simulated as a function of species-specific responses to *DrI, GDD*, winter temperature (see Table 1 for species parameter description, and Appendix A), light availability at the forest floor (see *realized tree growth* section), and browsing pressure (Didion et al. 2011). By default, the model assumes that there is a constant seed rain in the patches and thus no dispersal limitation, but alternatively it is possible to activate a feedback between the actual forest composition at year *n* and species composition of the new seedlings at year *n*+1. Seedling establishment can be constrained by defining a maximum number of seedlings potentially colonizing the patches and/or by imposing a feedback of actual forest composition on the composition of colonizing seedlings (Appendix A).

**Table 1:** Description of the species parameters in ForCEEPS. References refer to the literature used to calibrate all or part of the species for the specific parameter.

| Parameter | Details | Unit | Example<br>(*Fagus sylvatica*) | Calibration | References |
| --- | --- | --- | --- | --- | --- |
| $f_s$ | Foliage type | Unitless<br>E - evergreen - or D - deciduous and a number between 1 and 5 | D3 | Literature | Rameau et al. 1989; Didion et al. 2009 |
| $Hmax_s$ | Maximum height | m | 50 | Literature + NFI | Rameau et al. 1989, 2008 |
| $s_s$ | Allometry | Unitless | 76 | Literature + NFI | Didion et al. 2009 |
| $g_s$ | Optimal growth rate | Unitless | 260 | Literature + NFI | Didion et al. 2009 |
| $Amax_s$ | Maximum age | years | 400 | Literature | Rameau et al. 1989; Bugmann 1994 |
| $Ddmins$ | Minimal required annual or seasonal degree-days sum | °C | 841 | Literature | San-Miguel-Ayanz et al. 2016 |
| $DrTol_s$ | Drought tolerance index, to be compared to the evapotranspiration deficit based on a bucket model of soil moisture | Continuous index with values between 0 (sensitive) to 1 (tolerant) | 0.25 | Literature | Ellenberg and Mueller-Dombois 1966; Niinemets and Valladares 2006; Rameau et al. 1989, 2008 |
| $Nreq_s$ | Soil nitrogen requirement | Integer Index with values between 1 (weak requirements) to 5 (strong req.) | 2 | Literature | Ellenberg and Mueller-Dombois 1966; Bugmann 1994 |
| $ShTol_s$ | Shade tolerance | Integer index with values between 1 (shade tolerant) to 9 (shade intolerant) | 1 | Literature | Ellenberg and Mueller-Dombois 1966; Niinemets and Valladares 2006; Rameau et al. 1989, 2008 |
| $ShTol\_seedling_s$ | Shade tolerance of seedlings, to be compared to the relative amount of light reaching the ground | Continuous index with values between 0 (tolerant) to 1 (sensitive) | 0.05 | Literature | Ellenberg and Mueller-Dombois 1966; Niinemets and Valladares 2006; Rameau et al. 1989, 2008 |
| $Wtmin_s$ | Monthly minimum winter temperature tolerated for regeneration (°C) | °C | -6 | Literature | Didion et al. 2009 ; San-Miguel-Ayanz et al. 2016 |
| $Wtmax_s$ | Monthly maximum winter temperature tolerated for regeneration | °C | 9 | Literature | Didion et al. 2009 ; San-Miguel-Ayanz et al. 2016 |
| $Br_s$ | Browsing susceptibility of seedlings | Integer index with values between 1 (less susceptible) to 5 (more susceptible) | 3 | Literature | Didion et al. 2009 |

### Tree mortality

Tree mortality depends on two components: (i) a “background” mortality that is constant across time, and (ii) a growth-related mortality (Appendix A). The background mortality is purely stochastic. It depends on species’ maximum longevity and simulates mortality events induced by ‘random’ small-scale disturbances (e.g., attack of pathogen in an endemic phase). Large-scale disturbances (e.g., windthrows, wildfires) can be taken into account by increasing the background mortality rate, but are not considered here. The growth-related mortality is a proxy for stress conditions, i.e., tree mortality probability increases with the decrease in absolute or relative tree growth (i.e. tree vigor) induced by abiotic factors or by competition (DeSoto et al. 2020). It is thus noteworthy that competition has an indirect effect on mortality rates via the growth-related mortality.

### Potential tree growth

Annual tree growth is modelled through stem diameter increment at breast height (*ΔD*). Following the classical scheme of gap models, *ΔD* is calculated in two steps. First, the potential (i.e. maximum) diameter increment (*ΔD*_opt_) of each tree is predicted in each year using the following empirical equation (Moore 1989):

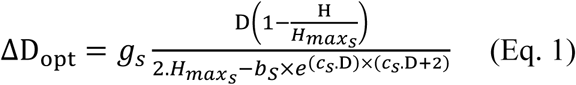

where *D* is tree diameter at breast height, *H* is tree height, *g*_*s*_ is the maximum growth rate of species *s*, 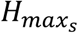 is the maximum height reachable by the species *s*, and *b*_*s*_ and *c*_*s*_ are species-specific parameters (with 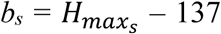; and *c*_*s*_ = *S*_*s*_ / *b*_*s*_); *s*_*s*_ is a species-specific allometric parameter relating tree height and diameter as follows (Bugmann 1996):

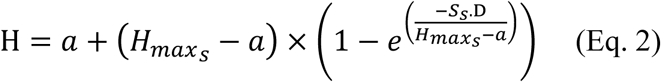

with a = 1.37 m (i.e. breast height). Therefore, simulating the potential diameter increment of a tree in ForCEEPS requires to determine the values of the species-specific parameters *g*_*s*_, *s*_*s*_ and 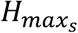 (Table 1).

### Realized tree growth

Realized tree diameter increment *ΔD* is calculated by modifying *ΔD*_opt_ according to abiotic or biotic growth reduction factors (all factors are bounded between 0 and 1) followong a modified geometric mean (Bugmann 1996, Didion et al. 2009):

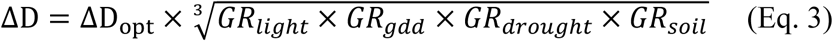

where *GR*_*light*_ is the growth reduction factor related to light availability for the tree, *GR*_*gdd*_ is the growth reduction factor related to growing season temperatures of the site (*GDD*), *GR*_*drought*_ is the growth reduction factor related to the site drought index (*DrI*), and *GR*_*soil*_ is the growth reduction factor related to soil nutrients content (*N*_*soil*_) (see Appendix A). The effects of each of these growth reduction factors on realized tree growth depend on species-specific parameters: *GR*_*light*_ depends on species shade tolerance *ShT*_*s*_; *GR*_*gdd*_ depends on species minimum sum of growing degree-days *GDD*_*S*_; *GR*_*drought*_ depends on species drought tolerance *DrT*_*s*_; and *GR*_*soil*_ depends on species requirements for soil nutrients *NReq*_*s*_ (see Table 1). All growth reduction factors vary among site conditions and species, and *GR*_*light*_ varies also among trees, because it is influenced by the sizes of the neighbouring trees in the patch (see next section).

### Effects of the competition for light on tree growth

In ForClim 2.9.6 (Didion et al. 2009), the amount of light available for a tree (with *H* being its total height) is reduced by the leaf area of the trees found in the same patch whose height is greater than *H* or equal to *H*. Thus, all the foliage of trees taller than the target tree contribute to the shading. ForCEEPS embeds a more realistic description of the competition for light, by representing individual crown lengths in the vertical space of the canopy (Fig. S1 and Appendix A).

In ForCEEPS, the growth reduction factor related to light availability (*GR*_*light*_) has two components:

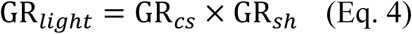

with *GR*_*cs*_ representing the feedback of crown size on tree growth, i.e., tree leaf area is positively linked to tree growth rate (Mitscherlich and von Gadow 1968). *GR*_*sh*_ is the reduction factor related to shading by competing trees. The key feature is that individual tree crowns are characterized by crown length *cl*, calculated as follows for each tree *i*:

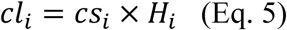

with *H* being tree height and *cs* being the ratio of the height with a green crown, which is related to light exposition of the tree (Didion et al. 2009). For each tree, *cs* varies between two extreme species-specific values that represent the case where the tree is fully shaded 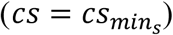 or in full light 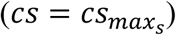, with:

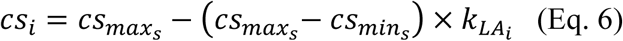

where the extreme values 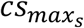 and 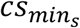 have been derived from the relationship between foliage fresh weight and DBH described in Wehrli et al. (2007) and depends on the foliage type parameter *f*_*S*_ (see Appendix A), and *k*_*LAI*_ is the correction factor - ranging from 0 (no shading) to 1 (full shading) - calculated by Didion et al. (2009) as follows:

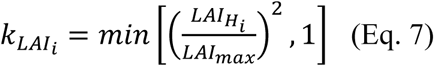

with *LAI*_*H*_ being the cumulative double-sided leaf area index between the top of the canopy and the top of the target tree (i.e. between the top of the canopy and the height *H*) and *LAI*_*max*_ being the maximum value of double-sided leaf area index in a patch, resulting from the light compensation point of the most shade-tolerant European tree species [i.e. *LAI*_*max*_ = 11.98 (Bugmann 1994, Didion et al. 2009)].

The vertical space of the patch *p* at simulation step t=*t*_*1*_ is discretized in *n(p,t*_*1*_) layers of a given width *w*, whose value is bounded between 0 (ground level) to *H*_*max*_(*p, t*_1_) (height of the tallest tree in the patch *p* at t=*t*_*1*_), with *w* = 1 m. We assumed that tree leaf area decreases linearly from the top to the base of the crown, i.e. from the highest to the lowest layer in which the crown of the tree is found (Fig. S1-B) (Eermak 1998, Van Pelt et al. 2016). We are aware that tree crown shape and vertical leaf area distribution vary among tree species and are also affected by the size and identity of neighbouring trees (Poorter et al. 2006, Williams et al. 2017, Niklaus et al. 2017). Our assumption should thus be seen as a first parsimonious step that can be refined using species- and context-specific architectural data. Further details about the calculation of *GR*_*cs*_ and *GR*_*sh*_ are described in Appendix A.

### Effects of the environmental conditions on tree growth

Belowground competition for water and nutrients is not explicit in ForCEEPS. However, while the model focuses on competition for light in its current version, it is noteworthy that soil nutrient content and soil moisture indirectly affect competition for light, in a way that differs among species (Table 1). In fact, *GR*_*soil*_ and *GR*_*drought*_ affect tree dimensions (diameter and height) (Eq. 3) and thus tree leaf area (Eq. 11), which in turn modifies the competitive ability of a tree because shading directly depends on leaf area (Eq. 12). Therefore, site conditions (soil and climate) modulate competition among trees.

## CALIBRATION, VALIDATION, AND SENSITIVITY ANALYSIS

### Species

The calibration and validation of ForCEEPS was done for nine species (Table S1) - four Angiosperm species and five Gymnosperm species, including the seven most widespread tree species in France (*Quercus petraea, Q. robur, Fagus sylvatica, Abies alba, Picea abies, Pinus sylvestris* and *P. pinaster*) (IGN 2018), and two emblematic species of Mediterranean French forests (*Pinus halepensis* and *Quercus ilex*). Furthermore, *P. pinaster* is the planted species covering the largest area in France. These species dominate in contrasted stages of the vegetation succession: pioneer (*Pinus*), intermediate-(*Picea*) or late-succession species (*Quercus, Fagus, Abies*).

Furthermore, for the PNV simulations, we complemented the set of studied species by considering 13 additional species (‘*other species*’ in Table S1) to cover most possible forest types: *Acer campestre, A. platanoides*, and *A. pseudoplatanus* (grouped in “*Acer*” species); *Larix decidua* and *Pinus cembra* (grouped in “*mountain gymnosperms*”); *Sorbus aria, S. aucuparia, and Ulmus glabra* (grouped in “*mountain broadleaves*”); *Betula pendula, Fraxinus excelsior* and *Populus tremula* (grouped in “*other broadleaves*” species); *Carpinus betulus*, and *Quercus pubescens*. However, no forest growth data was available to properly calibrate or validate the model for these other species as done for the nine main ones. This notably occurred because growth data at the stand scale were not available for these species (see Validation section) and growth data at the tree scale were only available for *C. betulus* and *Q. pubescens* (see Table S2).

The workflow of the study is summarized in Fig. 1.

**Figure 1.**
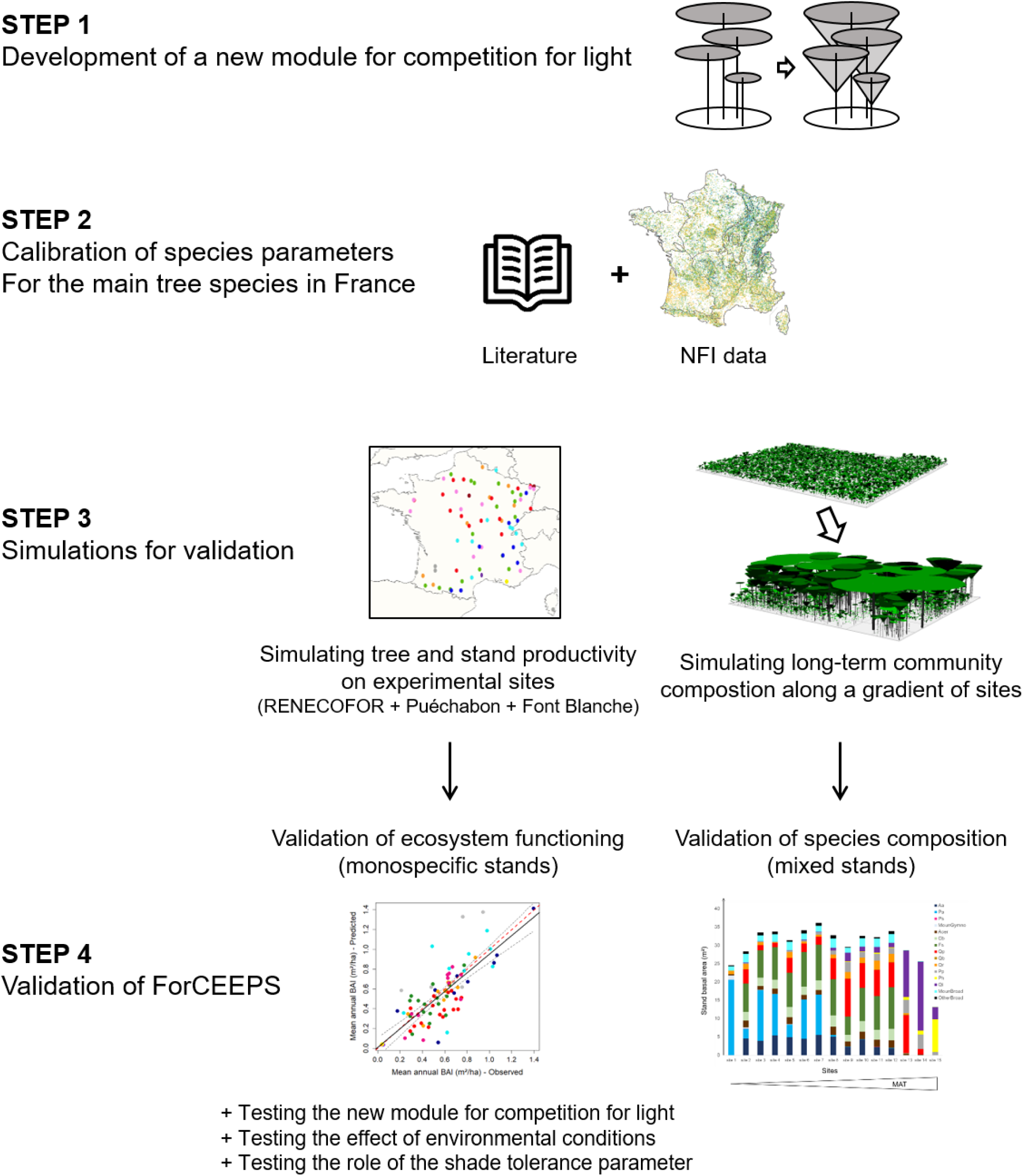
Summary of the workflow of the study. This figure illustrates the sequence of the main steps of the study.

### Calibration

Each species simulated in ForCEEPS is defined by 13 key parameters described in Table 1 (and Table S1) from which other parameters were derived (*b*_*s*_ and *c*_*s*_ in Eq. 1, *f*_*s*_ and *a*_*s*_ in Eq. 11, 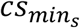, 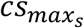 in Eq. 6, *LCP*_*s*_ in Eq. 8). The variability among functional traits reflects fundamental trade-offs of species life-history strategies (Bazzaz 1979, Violle et al. 2007). In ForCEEPS, like in many gap models, the variability among parameters’ values aims at reflecting such trade-offs (Bugmann 2001), and in this sense we further assume that the parameters describing the species in the model are proxies of life-history or functional traits. For instance, late-successional species are generally characterized by slow growth (i.e. low values of *g*_*s*_), long lifespan (i.e. low values of *Amax*_*s*_), and high shade tolerance (i.e. low values of *ShTol*_*s*_), in contrast to early-successional ones (Reich 2014).

In the present study, the calibration of potential tree growth (i.e., species-specific parameters *g*_*s*_ and 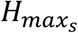) and the allometry relating tree height and diameter (i.e., parameter *s*_*s*_) were based on data from the French National Forest Inventory (NFI) (IGN 2018). The values of other parameters were based on the literature. The NFI sampling design warrants an exhaustive representation of environmental gradients within the realized distribution of the species over the mainland French territory, while individual plots may not be locally representative (Charru et al. 2010). Therefore, we used NFI to calibrate the potential growth model in ForCEEPS, but did not use it for the validation at the plot level. More detailed information about NFI data is available in Appendix D.

#### Parameter g_s_

This parameter is the most difficult to calibrate as it requires data from trees growing in “optimal conditions”, which are scarce in observational datasets as the growth of trees is usually constrained by environmental conditions or biotic factors (e.g. competition). To cope with this challenge, we took advantage of the NFI that covers a very wide range of conditions (in both space and time), providing a large number of “annual diameter increment *vs*. diameter” pairs for each of the 11 species (i.e., the nine main species and *C. betulus* and *Q. pubescens*) for which abundant data were available (n=206,569 for all species confounded, Table S2). For each of these 11 species, we grouped trees according to their diameter (according to 1-cm size classes) and selected the 10% of trees with the greatest annual diameter increment, assuming that these trees grew in “optimal conditions” or at least under almost unconstrained conditions. However, we note that the annual increments are derived from five-year average, which may lead to an underestimation of the actual greatest annual diameter increments. Then we fitted *g*_*s*_ from Eq. 1 with this dataset, using a non-linear least squares approach implemented by the nls function in the R software (R Core Team 2018). For the remaining species (n = 11), the *g*_*s*_ values have been set from previous studies (Didion et al. 2009).

The fitted values for the parameters *g*_*s*_ ranged from 79 to 399 (Table S1). These values are consistent with former estimates for the same or related species (Bugmann 1994, Didion et al. 2009).

#### Parameter s_s_

The calibration of *s*_*s*_ (Eq. 2) relied on NFI data because of their representativeness of the conditions in which each species occurs. The whole NFI dataset was used for the calibration to cover the largest range of conditions in which each species occurs. Although diameter-height relationships were shown to be affected by environmental conditions, e.g. climate, tree social status and stand density (Trouvé et al. 2015, Fortin et al. 2019), these factors were not accounted for in the model. The rationale for this lies in our aim to keep the model structure as simple as possible to allow for an easy parameterization and use at large scale for a large number of species. We fitted the height-diameter relationships (Eq. 2) on the NFI dataset, using the nls R function, and extracted *s*_*s*_ values for each species. As for *g*_*s*_, this calibration was conducted for the 11 main species, while we relied on Didion et al. (2009) for the 11 additional species.

#### Parameter 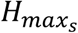

This parameter was calibrated using NFI data and/or literature (Rameau et al. 1989, 2008) for all the species. Maximum height may indeed be underestimated in the NFI data because forest managers tend to harvest the largest trees before they reach their maximum height.

#### Other parameters

The values of the parameters describing species’ response to abiotic conditions (i.e. effect of the growing season temperature on tree growth, *DDmin*_*s*_; drought tolerance, *DrTol*_*s*_; and soil nitrogen requirement, *Nreq*_*s*_), and species intrinsic characteristics (i.e. foliage type, *f*_*s*_; maximum age, *Amax*_*s*_, shade tolerance, *ShTol*_*s*_, and shade tolerance of seedlings *ShTol_seedling*_*s*_, browing susceptibility of seedlings *Br*_*s*_) were based on the literature (Table 1 and references therein). Parameters describing the thermal regeneration niche for seedlings (i.e., monthly minimum and maximum winter temperature tolerated for regeneration *WTmin*_*s*_ and *WTmax*_*s*_, Table 1) were calibrated according to species-specific diagrams of occurrence (San-Miguel-Ayanz et al. 2016).

#### Congruence of key parameter values with functional traits

To gain mechanistic insight into the parameters values derived from the calibration procedure, we evaluated the congruence of key model parameters with functional traits extracted from the literature. To do so, we first selected the most meaningful ForCEEPS parameters in terms of species ecological strategies, including *g*_*S*_, *DrTol*_*S*_, *ShTol*_*S*_, *ShTol_seedling*_*S*_, and *Nreq*_*S*_. Then we collected data on relevant traits from various database, including: xylem cavitation resistance (assess through the water potential causing 50% cavitation, *Ψ50* in MPa), leaf turgor loss point (*Ψtlp*, in MPa), water potential causing stomatal closure (*Ψclose*, in MPa) and safety margins from *Ψtlp* and *Ψclose* [from the SurEAu database, Martin-StPaul, Delzon, & Cochard (2017)], wood density (g/m^3^, Chave et al. 2009), light saturated CO2 assimilation (or maximal photosynthesis *Amax*, in µmol/m^2^/s), nitrogen content per unit leaf area *Na* (g/m^2^) and leaf mass per area *LMA* (g/m^2^) [from the CANTRIP database, Keenan & Niinemets (2016)]. The final trait database and associated references are reported in Appendix E. For each of the selected ForCEEPS parameters, we tested the Pearson’s correlations between the ForCEEPS parameters and some of the traits at the interspecific level. Note that the consistence of the results across both Angiosperms and Gymnosperms was taken into account to assess the robustness of the congruence of species parameters with functional traits.

### Validation against forest growth data

#### Forest growth dataset

The validation of simulated annual productivity at the tree and stand levels was conducted using a dataset independent from the one used in the calibration process. Following Guillemot et al. (2017), we primarily relied on the RENECOFOR permanent forest plot network (Ulrich 1997) that includes 103 half-hectare plots in even-aged managed forests covering most of the main tree species and climate conditions in France. After excluding the plots that had experienced a natural or anthropic disturbance (e.g., thinning) less than 4 years before the last diameter inventory, 77 plots remained. Most of the stands included in the validation dataset are monospecific or strongly dominated by one species

The RENECOFOR network does not include forests growing under Mediterranean conditions. Therefore, we completed the validation by using data from the long term experimental sites of Puéchabon (*Quercus ilex*, Rambal et al. 2014) and Font Blanche (mixed forest dominated by *Pinus halepensis*, Simioni et al. 2016). Diameter inventories were used to estimate the tree and stand basal area increment (BAI) in all validation plots. The time interval between the initial and final inventories in RENECOFOR plots varied between 4 and 14 years, while they were of 14 and 10 years for the Puéchabon and Font Blanche sites, respectively (see further details about the validation datasets in Appendix D). The BAI data recorded over contrasted time intervals were normalized to mean annual BAI. Local measurements of soil water holding capacity (SWHC) were available for all plots, and climate time-series were obtained from the SAFRAN atmospheric reanalysis (Vidal et al. 2010) for the RENECOFOR plots, and from on-site measurements for the Puéchabon and Font Blanche plots. The validation plots covered a large range of environmental conditions, with mean annual temperature (*MAT*) ranging between 5.8°C and 14.3°C, mean annual precipitation sum (*MAP*) between 700 and 2030 mm, while the drought index ranged from 0.003 to 0.35 (values below 0.05 indicate there is no marked drought stress for the trees, while values above 0.3 indicate strong stress for most tree species) (Fig. S2).

#### ForCEEPS simulations

We initialized the model for each stand using the first inventory campaign of the respective plot. For each RENECOFOR plot, 5 patches of 1000 m^2^ were simulated, in order to obtained comparable observed and simulated forest plot areas (the average size of the observed plots is ca. 5000 m^2^). To simulate the patches, trees were randomly sampled in the inventory dataset of a given plot until the stand basal area per square meter of the simulated patch was comparable to the observed stand basal area per square meter. Local measurements of SWHC and local climate time-series were used as inputs. ForCEEPS simulations were run over the time period for which BAI measurements were available in each plot (i.e. from 4 to 14 years), and subsequently normalized to mean annual BAI. As the results were very consistent across the five repetitions carried out per plot (as shown in Fig. S3 for the RENECOFOR plots), we only present the results for one repetition at the tree level for the sake of clarity (the results for each repetition are shown in supplementary material - Table S4 and Fig. S3). For results at stand level, we present averages across the five repetitions (the results for each repetition are shown in supplementary material - Table S4).

Gap models like ForCEEPS are designed to explore processes occurring at the stand level and are thus more relevant at this scale. However, as neighborhood interactions are reported to be key in driving BEF relationships and for the sake of comprehensiveness, we also present the results at the tree level (Schnabel et al. 2019, Jourdan et al. 2019a).

### Quantifying the importance of the hypotheses embodied in ForCEEPS for forest growth

After simulating BAI for each plot using the full model, we carried out three types of simulations to quantify the importance of some hypotheses and ecological processes embedded in ForCEEPS. First, we ran simulations without the new module for competition for light, to test whether an explicit representation of individual crown lengths in the vertical canopy space increased the prediction accuracy of stand growth (*Test 1*). Second, we ran simulations without considering the limiting effect of drought stress and thermal constraints on tree growth, i.e. under optimal climatic conditions (*Test 2*). Third, we aimed at testing the importance of the species-specific tolerance to shade in ForCEEPS (*Test 3*), as it has been shown to be a key parameter driving diversity effects in ForClim 2.9.6 (Morin et al. 2011). To do so, we changed the specific values of the parameter *ShTol*_*S*_ by assigning the maximum value to all species. Note that this kind of tests has been rarely done with gap models [but see (Morin et al. 2011, Huber et al. 2018)].

### Validation against potential natural vegetation

#### Study sites

To validate the model’s predictions in terms of outcomes of climate effects and interspecific competition in the long term, we compared the community composition simulated by ForCEEPS with the tree species composing the potential natural vegetation (PNV) along an environmental gradient. Defining PNV for a given site is subject to personal judgment. Here, similarly as in Bugmann (1996), we simply relied on the assumed dominant tree species (assuming no large disturbances) in a space spanned by annual precipitation (*MAP*) and mean annual temperature (*MAT*), following Ellenberg (1986), Rameau et al. (1989, 2008) and San-Miguel-Ayanz et al. (2016) (Fig. 4-B). More precisely, we selected 14 sites with contrasted conditions among the 79 plots used for the validation of forest growth simulations. This gradient thus includes dry and warm conditions through the two Mediterranean sites, but it did not include the coldest conditions in which forests can grow in France. Therefore we added another site with average MAT of 2.9°C (±0.64) and ASP of 1577 mm (±253), corresponding to the conditions of a subalpine site according to Ellenberg (1986) (grey dot in Fig. S2, and site 1 in Fig. 4-B).

#### ForCEEPS simulations

For each of the 15 sites, we ran 2500-yr simulations, starting from bare ground. Thus, the PNV simulations accounted for seedling establishment, tree growth and mortality. This simulation duration was necessary to avoid the communities to be in a transient phase and to ensure that they reached a pseudo-equilibrium in terms of composition and basal area. The 2500-yr climate time-series were obtained by randomizing the years from which time-series were available for each site. In other words, we considered inter-annual variability in climate, but there was no trend in the long term, as commonly done in studies aiming at depicting forest succession with gap models (e.g., (Bugmann 1996, Morin et al. 2011, Chauvet et al. 2017). We considered 200 patches of 1000 m^2^ for each simulation. At the end of the simulation, we extracted the mean basal area per hectare of the simulated stands and the basal area of each species.

The performance of the model was assessed using Pearson correlation coefficient (r), the root mean square error (RMSE), and the average bias (AB) between observations and model predictions.

## RESULTS

### Prediction of aboveground tree growth

ForCEEPS was able to capture the observed mean annual BAI (Fig. 2) at the tree level, with a good correlation between observations and predictions (r=0.72, n=2662; Table 2), while the difference between observations and predictions was satisfactory (RMSE = 0.0012, AB = 12.4%). There was, however, a slight tendency to underestimate the growth of the most productive trees (Fig. 2), and the uncertainty of the model predictions increased with tree diameter (Fig. S6). When the species were examined separately, the Pearson correlation coefficient ranged from 0.49 (*P. sylvestris*) to 0.77 (*F. sylvatica*) (Table 2, Fig. S4) but the difference between observations and predictions strongly varies between species (RMSE = 0.0013 and AB = 21.8%, on average).

**Table 2:** ForCEEPS accuracy in predicting tree growth across all species and for each species separately, through Pearson correlation, root mean square error (RMSE) and average bias (AB) between observed and predicted tree growth. Significance of the Pearson correlation coefficient (***: p < 0.001; **: p < 0.01; *: p < 0.05; *ns*: p > 0.05).

|  | All trees | *F. sylvatica* | *Q. petraea* | *Q. robur* | *A. alba* | *P. abies* | *P. sylvestris* | *P. pinaster* | *Q. ilex* | *P. halepensis* |
| --- | --- | --- | --- | --- | --- | --- | --- | --- | --- | --- |
| Ntrees | 2662 | 877 | 272 | 183 | 367 | 319 | 323 | 197 | 91 | 34 |
| Nplots | 82 | 16 | 16 | 10 | 9 | 9 | 11 | 6 | 3 | 2 |
| Pearson r | 0.72*** | 0.77*** | 0.66*** | 0.65*** | 0.74*** | 0.603*** | 0.491*** | 0.561*** | 0.616*** | 0.694*** |
| RMSE | 0.0012 | 0.0009 | 0.0011 | 0.0014 | 0.0014 | 0.0027 | 0.001 | 0.0014 | 0.001 | 0.0005 |
| AB | 0.124 | -0.024 | -0.050 | 0.334 | -0.227 | 0.252 | 0.367 | 0.898 | -0.073 | 0.481 |

**Figure 2.**
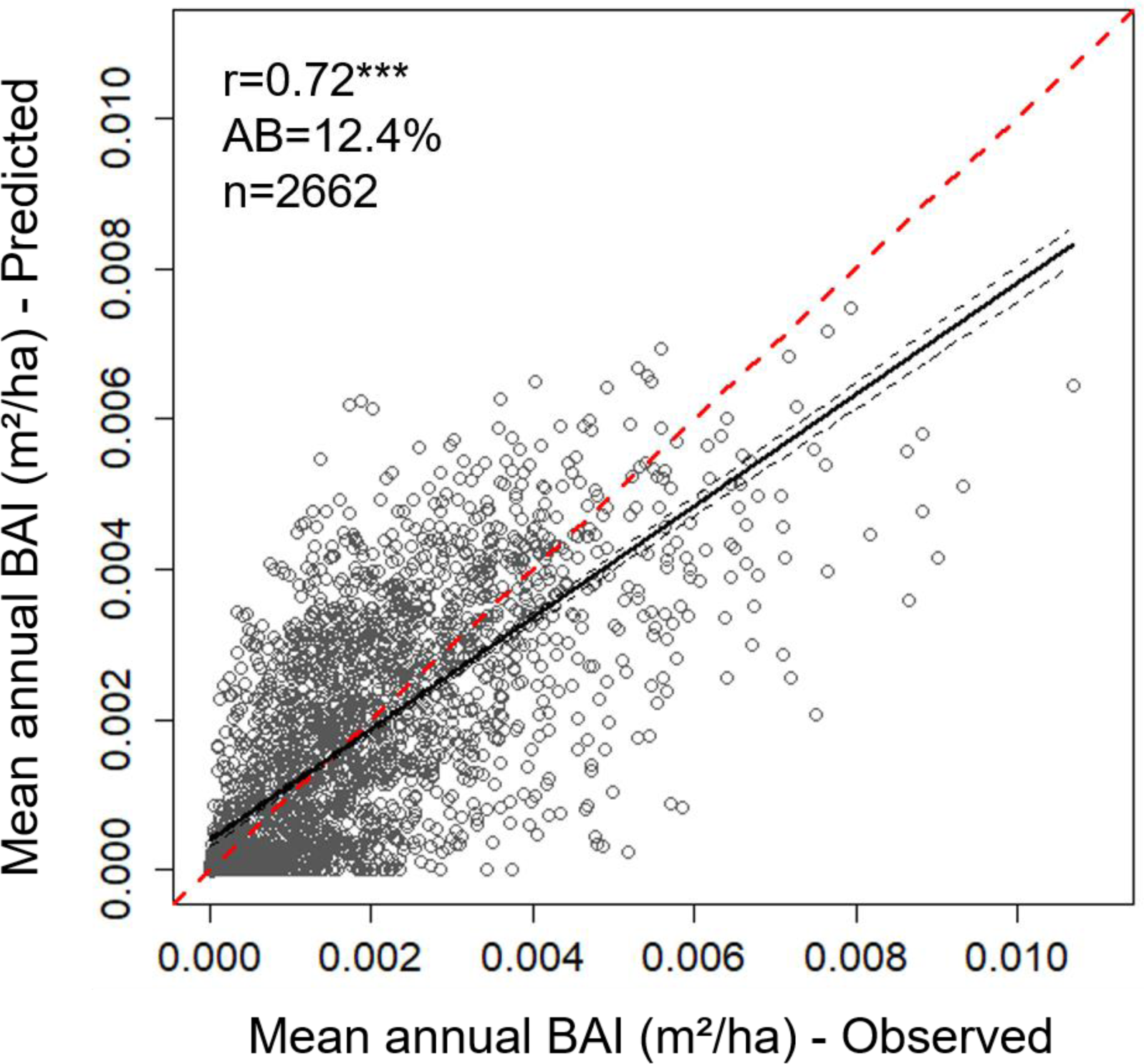
Predicted (by ForCEEPS) against observed mean annual tree basal area increment (BAI) for all considered trees (over 82 sites) and the 5 repetitions. The plain black line is the regression line of the linear model of the relationship between observed and predicted tree growth, with confidence interval represented with the grey dashed lines; the dashed red line is the 1:1 line. Statistics associated: see Table 2.

### Prediction of aboveground stand growth

At the stand level, ForCEEPS showed a good ability to reproduce observed mean annual BAI regardless of the species or the environmental conditions. Across all plots, the correlation was strong between observations and predictions (r=0.79, P<0.001, Table 3) with a very low difference between observations and predictions (RMSE = 0.019 and AB=4.5% – Fig. 3-A, Table 3) without strong bias related to the basal area of the stand (Fig. S7). When species were examined separately, the accuracy varied across species, but the results did not show strong systematic bias (Fig. S5, RMSE = 0.014 and AB = 26.7% on average, Table 3-b) except for *Q. petraea*, for which productivity of the most productive plots was underestimated (RMSE=0.016 and AB=-16.7%, Fig. S5), and *P. pinaster*, which showed the highest variability (RMSE=0.034 and AB = 50.3%, but it is the species with the smallest number of observations – except *Q. ilex* and *P. halepensis* for which one can hardly make any conclusion with only three plots).

**Table 3:**
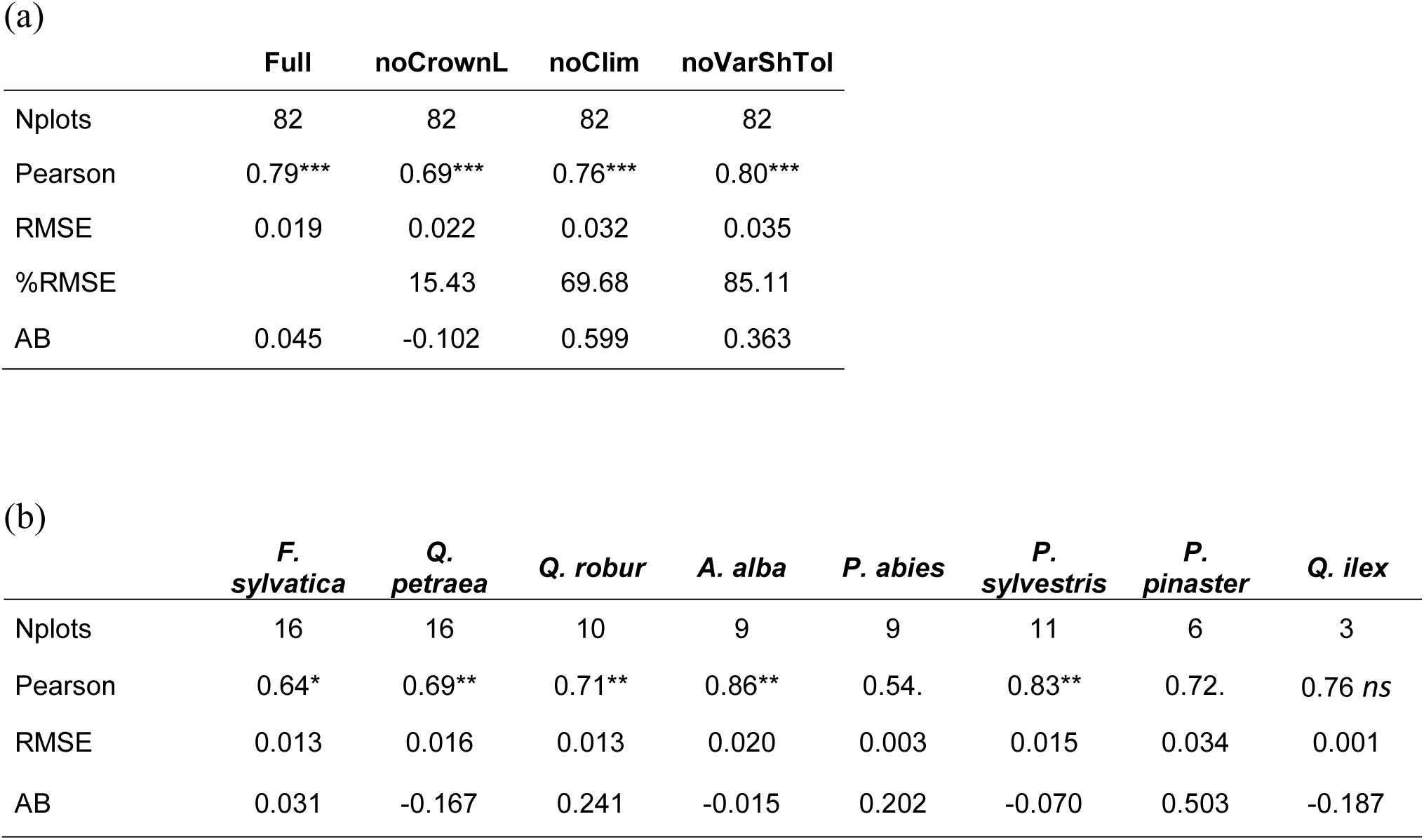
ForCEEPS accuracy in predicting stand productivity and test of the differences between the various versions tested: (a) Across all plots, though through Pearson correlation, root mean square error (RMSE) and average bias (AB) between observed and predicted stand productivity. *Full*: ForCEEPS simulations with the new crown length module, climatic constraints on tree growth and interspecific variability in parameter *ShTol*_*S*_; *noCrownL*: ForCEEPS simulations without the new crown length module; *noClim*: ForCEEPS simulations without climatic constraints on tree growth; *noVarShTol*: ForCEEPS simulations without interspecific variability in parameter *ShTol*_*S*_. %RMSE: percentage difference between the RMSE of the version tested and the “full” version. (b) For each species taken separately for the full version, through Pearson correlation, root mean square error (RMSE) and average bias (AB) between observed and predicted stand productivity tree growth. Significance of the Pearson correlation coefficient (***: p < 0.001; **: p < 0.01; *: p < 0.05; *ns*: p > 0.05).

|  | Full | noCrownL | noClim | noVarShTol |
| --- | --- | --- | --- | --- |
| Nplots | 82 | 82 | 82 | 82 |
| Pearson | 0.79*** | 0.69*** | 0.76*** | 0.80*** |
| RMSE | 0.019 | 0.022 | 0.032 | 0.035 |
| %RMSE |  | 15.43 | 69.68 | 85.11 |
| AB | 0.045 | -0.102 | 0.599 | 0.363 |

(b)
|  | *F.*<br>*sylvatica* | *Q.*<br>*petraea* | *Q. robur* | *A. alba* | *P. abies* | *P.*<br>*sylvestris* | *P.*<br>*pinaster* | *Q. ilex* |
| --- | --- | --- | --- | --- | --- | --- | --- | --- |
| Nplots | 16 | 16 | 10 | 9 | 9 | 11 | 6 | 3 |
| Pearson | 0.64* | 0.69** | 0.71** | 0.86** | 0.54. | 0.83** | 0.72. | 0.76 *ns* |
| RMSE | 0.013 | 0.016 | 0.013 | 0.020 | 0.003 | 0.015 | 0.034 | 0.001 |
| AB | 0.031 | -0.167 | 0.241 | -0.015 | 0.202 | -0.070 | 0.503 | -0.187 |

**Figure 3.**
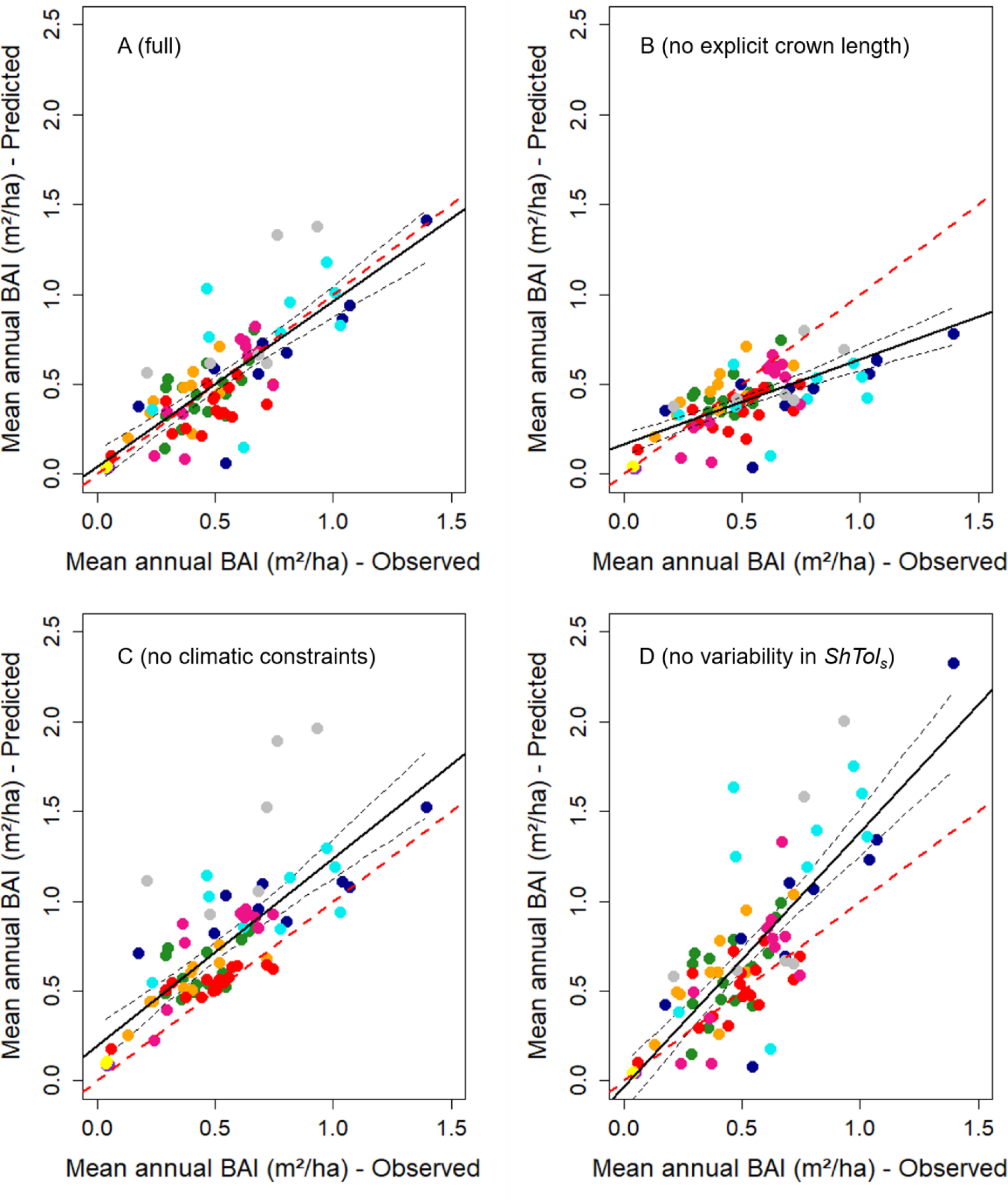
Predicted (by ForCEEPS) against observed mean annual stand basal area increment (BAI) for the 82 sites, using different model configurations: A-ForCEEPS simulations with the new crown length module, climatic constraints on tree growth and interspecific variability in shade tolerance (parameter *ShTol*_*s*_). B-ForCEEPS simulations without the new crown length module. C-ForCEEPS simulations without climatic constraints on tree growth. D-ForCEEPS simulations without interspecific variability in parameter *ShTol*_*s*_. For all panels: the plain black line is the regression line of the linear model of the relationship between observed and predicted stand productivity, with confidence interval represented with the grey dashed lines; the dashed red line is the 1:1 line. Statistics associated: see Table 3-a. Colour code for the species as follows: • Fagus sylvatica •Quercus robur • Quercus petraea • Quercus ilex • Abies alba • Picea abies • Pinus sylvatica • Pinus pinaster • Pinus helepensis

### The importance of light competition, environmental conditions and shade tolerance for simulating forest growth in ForCEEPS

#### Testing the representation of light competition

The new module for competition for light, which include an explicit representation of individual crown lengths in the vertical canopy space, yielded on average better results than the former version (decrease by 15.4% in RMSE; Table 3-a). The former version tended to underestimate the productivity of the most productive plots, while this was not the case with the new version (Fig. 3-A and 3-B).

#### Testing the effect of environmental conditions

The model without climatic constraints on tree growth was less accurate than the full version (increase by 69.7% in RMSE; Fig. 4-A and 4-C; Table 3-a), except for a few plots - especially for *Q. petraea* stands. The simulations without climatic constraints logically tended to overestimate stand productivity (Fig. 3-C). It is thus noticeable that on average, the effect of climatic conditions improved the accuracy of the simulations over such a large range of environmental conditions tested in this study (illustrated in Fig. S2). One may also notice that this improved accuracy is consistent across species, regardless their averaged productivity.

#### Testing the importance of the variability in the shade tolerance parameter

When the variability in the ability of species to tolerate shade was not taken into account in ForCEEPS, the model’s performance strongly decreased, with an increase in RMSE by 85.11% across plots (Fig. 3-A and 3-D; Table 3-a). The bias notably increased for the most productive stands, especially dominated by *A. alba* and *P. abies* (Fig. 3-D).

### Prediction of species composition in the long term

When comparing the distribution of the dominant tree species at the end of the 2500-yr simulations carried out along the environmental gradient covered by the 15 sites (Fig. 4), it appeared that the ability of ForCEEPS to predict reliable PNV varied across sites: the overall likelihood of the simulated communities is strong, but with be a greater uncertainty about Mediterranean forest types. In 10 out of the 15 sites, the dominating species were accurately predicted according to the PNV diagram (green dots in Fig. 4-B). In the five other sites, at least one of the dominating species was accurately predicted (blue dots in Fig. 4-B), while there was no site in which the simulated community was dominated by species other than those expected. Long-term simulation of stand basal area cannot be directly evaluated against field observations as there is no forest stands unaffected by management for several centuries at these sites. Yet, one may notice that the values appear consistent (albeit a bit low) with mature stands, and that the simulated basal area was lower in the harshest conditions (i.e., at both extremes of the gradient). However, the basal area for the Font Blanche site seemed to be underestimated (ca. 15 m^2^) (Simioni et al. 2016).

It is noticeable that the cumulated basal area of the species that were not validated against forest growth data in the present study (ie. the “other species” in Table S1) represent on average only 17% (across the 15 sites) of stand basal area at the end of the simulations, and it remains below 25% at all sites.

### Congruence of key parameter values with functional traits

We found correlations between traits and ForCEEPS parameters, but their sign and significance strongly varied. The species nitrogen requirement *Nreq*_*S*_ was found to correlate with *Na* (Table S5). The growth parameter *g*_*S*_ was significantly negatively correlated with wood density (Figure 5), while the correlation with *LMA* was not consistent for Angiosperms and Gymnosperms (Table S5). Seedling and adult shade tolerance were correlated with light saturated photosynthesis (*A*_*max*_, Fig. 5 and Table S5). Other traits, including *LMA* and wood density were poorly correlated with shade tolerance. Finally, correlations were found between *DrTol*_*S*_ and different drought-related functional traits. In particular, a strong correlation was found between *DrTol*_*S*_ and the stem xylem embolism resistance (assessed by *P50*, i.e., the water potential causing 50% embolism, Fig. 5). The correlation between *DrTol*_*S*_ and *P50* was very strong for angiosperms (r^2^=0.7, *p*<0.001) but not significant for gymnosperms (*p* = 0.1), which could be explain by the fact that the studied conifers all belong to the *Pinaceae* family that rely on a tight stomatal control of transpiration during drought. Positive but less pronounced relationships were found between *DrTol*_*S*_ and the turgor loss point (Table S5). *DrTol*_*S*_ was also correlated with wood density and *LMA* but to a lower extent (Table S5).

**Figure 4.**
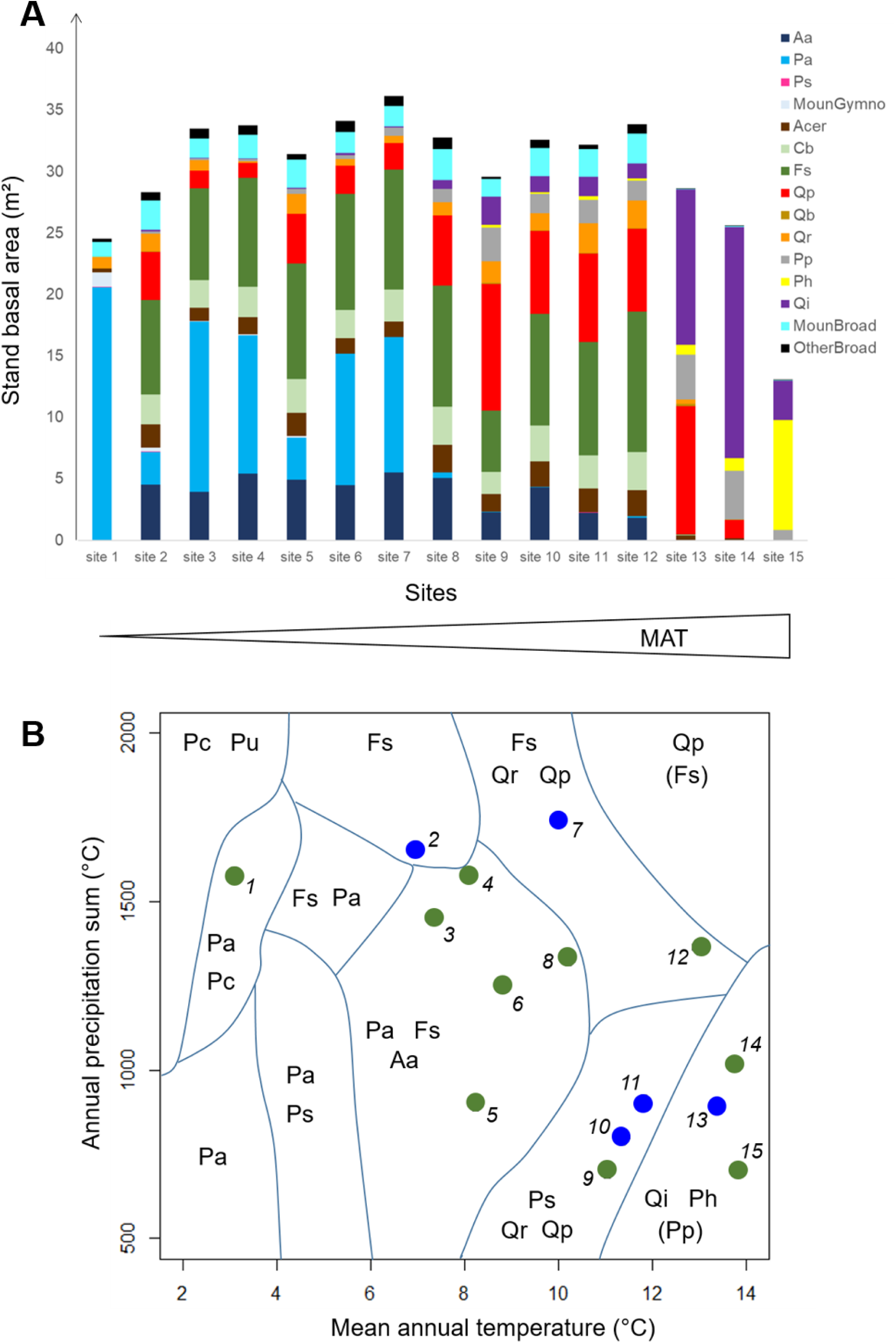
(A) Simulated basal area (m^2^ per ha) at the end of long-term ForCEEPS simulations along sites representing a gradient of environmental conditions from cold and moist alpine conditions (left) to warm-dry Mediterranean conditions (right). The site names and conditions are stated in Table S3, with Aa (*A. alba*); Pa (*P. abies*); Ps (*P. sylvestris*); Cb (*C. betulus*); Fs (*F. sylvatica*); Qp (*Q. petraea*); Qb (*Q. pubescens*); Qr (*Q. robur*); Pp (*P. pinaster*); Ph (*P. halepensis*); Qi (*Q. ilex*); MounGymno (mountainous gymnosperm species including *L. decidua* and *P. cembra*); MounBroad (mountainous broadleaf species including *S. aria, S. aucuparia* and *U. glabra*); OtherBroad (broadleaf species including *B. pendula, F. excelsior* and *P. tremula*). (B) Distribution of the 15 tested sites in the PNV diagram of the supposed dominating species (built according to mean annual temperature and annual precipitation sum). Green dots: sites for which the dominating species in the simulated communities were accurately predicted according to the PNV diagram; Blue dots: sites for which at least one of the dominating species was accurately predicted but with another dominating species not supposed to dominate according to PNV diagram. Red dots: sites in which the simulated community was dominated by other species than supposed by the PNV diagram. Numbers refer to the site number (see Table S3). PNV dominating species are Pc (*P. cembra*), Pu (*P. uncinata*); Aa (*A. alba*); Pa (*P. abies*); Fs (*F. sylvatica*); Qp (*Q. petraea*); Qr (*Q. robur*); Pp (*P. pinaster*); Ph (*P. halepensis*); Qi (*Q. ilex*).

**Figure 5.**
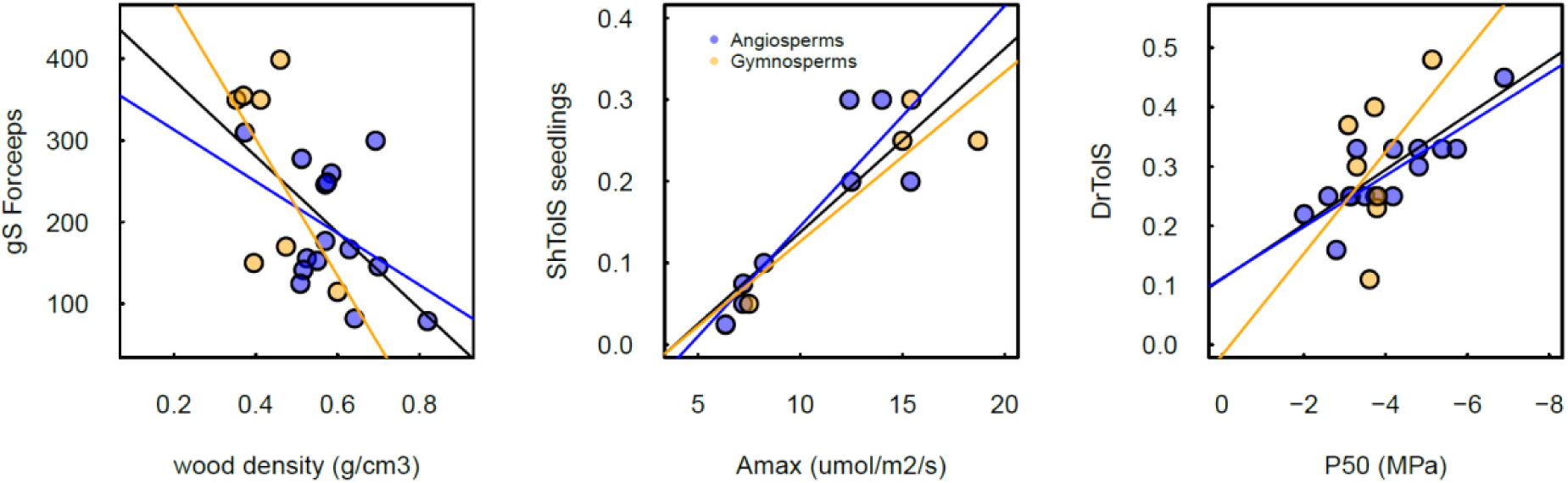
Correlations between key ForCEEPS parameters and ecophysiological traits extracted from the literature (see Appendix E). Blue dots: Angiosperms; orange dots: Gymnosperms. Associated statistics are presented in Table S5.

## DISCUSSION

### A gap model predicting annual productivity and community composition

ForCEEPS relies on ecological hypotheses, notably the trade-off between maximum growth and tolerance to competition (Rees et al. 2001) and the fact that cyclical succession is occurring in each individual patch (Botkin et al. 1972), allowing to simulate long-term species ecological succession. Although most biogeochemical processes are implicit in the model, as in most gap models, our results show that ForCEEPS accurately predicts both the dominant species occurring at a site in the long term and the wood productivity of monospecific stands across a few years.

Gap models have long demonstrated their ability in predicting the long-term dominant species of forests (Bugmann 2001), but it is noticeable that ForCEEPS appeared robust across a large range of environmental conditions, i.e. from alpine to Mediterranean forests. Indeed, if gap models were already shown to accurately predict dominant species composition in temperate and subalpine forests (e.g. Bugmann 1996; Didion et al. 2009), the good performances of ForCEEPS at Mediterranean sites appears as a major achievement. Although this validation remains mostly qualitative, the accuracy of predicted community composition from the long-term simulations is remarkable, and suggests that the interspecific competition and abiotic constraints are well represented in ForCEEPS. The good performances of ForCEEPS across large environmental gradients and for the most important tree species found in mainland France, suggest that the model could be applied to a large part of the European forest ecosystems.

The validation of the ability of ForCEEPS to predict forest functioning in the short term (i.e. across a few years) was conducted using forest growth data from monospecific stands. The rationale for this choice was to evaluate its behavior and predictive ability in a context with low influence of complex interspecific interactions. Because gap models are often validated using species composition of PNV at selected sites, their validation is actually conducted in mixed forests in most cases (Bugmann 2001). Thus, this test of the ability of gap models to accurately simulate the functioning of monospecific stands in various environmental conditions and for a wide range of species has been very rarely assessed. Yet, monocultures are often compared to mixed stands to quantify biodiversity effects in forests (e.g., as in Morin et al. 2011). Ensuring that the functioning of monospecific stands is well reproduced by a gap model is thus a *sine qua non* condition to simulate non-biased biodiversity effects in tree communities.

Validation against forest growth data was rarely done for gap models (Bohn et al. 2014), especially for such a wide range of species and conditions. Gap models have not originally been designed to work at short temporal scales, and are thus not expected to accurately simulate annual tree or stand growth (Mette et al. 2009, Fyllas et al. 2014). Although ForCEEPS may never offer detailed mechanistic insights into ecosystem biogeochemistry and tree growth as ecophysiological models do (Makela et al. 2000, Dufrêne et al. 2005, Guillemot et al. 2017), it can nevertheless be considered as a parsimonious alternative – notably in terms of calibration to explore how productivity will respond to changes in species composition and climate.

Recent advances in forest ecology have resulted in physiological process-based models that can be fully parameterized (e.g. Maréchaux and Chave 2017, Martin-StPaul et al. 2017) using functional traits available from global databases (Kattge et al. 2011). Although these models provide a unique insight on the physiological mechanism driving forest growth and survival, they are not aimed to describe the long-term ecological processes shaping forest composition on the long-term. In this study, we evidence that the processes embodied in gap models to simulate long-term forest succession can also predict annual forest growth in species with contrasted ecology and under various climate conditions, making them an important tool to study forest responses to climate change. ForCEEPS requires a rather small number of parameters to describe a species, allowing both a straightforward calibration of some parameters using forest inventory data and an *a priori*-calibration of the other parameters relying on literature and ecological knowledge. Consequently, the hypotheses embodied in ForCEEPS regarding the complex feedback loops and threshold mechanisms that drive forest functioning and forest community dynamics can be conceptualized, parameterized and evaluated against measured field data. This limits the uncertainty that can affects model predictions in case of equifinality. Of course, ForCEEPS - like all gap models - could also greatly benefit from the current increasing availability of forest inventory data to improve its calibration using inverse modeling approaches (Hartig et al. 2012).

### Hypotheses, limitations and future directions to improve the model

The high accuracy of ForCEEPS in predicting mean annual stand productivity of forests over a few years thus opens great perspectives for ecological studies. However, this potential should not conceal the simplifications and limits of our approach. Our results showed that explicitly representing 2D-competition for light by considering crown size in the vertical canopy space improved the accuracy of the predictions of short-term productivity compared to the ‘classic’ scheme of gap models (Bugmann 2001). Meanwhile, this novel development did not affect the reliability of the model’s predictions of community composition and standing biomass in the long term. Yet, introducing this change in the model implied to make some assumptions on crown traits and foliage distribution in vertical space. There is an increasing number of studies showing that these properties vary depending on species identity (Bayer et al. 2013, Forrester and Albrecht 2014, Forrester et al. 2018), and the size and identity of neighboring trees (Poorter et al. 2006, Williams et al. 2017, Niklaus et al. 2017). While future work may further improve the representation of canopy space exploration by taking into account the plasticity of tree architecture, we believe that the current version of the model relies on a sufficiently parsimonious approach to explore new questions regarding above-ground tree-tree interactions in mixed stands. Keeping track of tree coordinates in horizontal space - as already done in other models (Bohn et al. 2014, Maréchaux and Chave 2017) - would allow to more finely tackle the mechanism driving tree interactions, but this may come at the cost of losing the generality of the model, as well as strongly increasing the simulation time.

We demonstrate in this study that both the climatic constraints and the variability in species’ shade tolerances are crucial to predict short-term productivity with gap models. In particular, we showed that differences in shade tolerance among species are key community features driving diverse forest productivity, which has not been shown across such a wide environmental gradient to our knowledge (Toïgo et al. 2018, Van de Peer et al. 2018). In turn, this reinforces the need for further exploration of light-mediated tree interactions to understand the mechanisms driving species assemblage and productivity in mixed forests. Although these quantifications are necessarily related to the way the climatic growth-reducing factors and competition for light are modelled, they nevertheless provide an *a posteriori* justification of the processes embedded in these models. This also confirms the large potential of such models for exploring how diversity affects forest functioning (Toïgo et al. 2018, Van de Peer et al. 2018, Cordonnier et al. 2018a) and how climate change is mediating this effect (Morin et al. 2018).

Yet, this study considered short-term growth, i.e. tree or stand growth averaged across a few years. Testing the performance of ForCEEPS on actual annual data of tree and stand increments would have constituted an even stronger test. However, this kind of data is rarely available for all trees on ∼1000 m^2^ plots (see Nehrbass-Ahles et al. 2014), especially for large number of species and range of environmental conditions.

For the sake of generality, ForCEEPS relies on generic DBH-height relationships, although DBH-height relationships are known to change with tree age and tree density (Trouvé et al. 2015, Fortin et al. 2019). Improvements in this direction may be possible, even though calibrating this allometric parameter would require more detailed inventory data (Rasche et al. 2012), and may have a very limited effect on the model’s results when compared to the effect of other parameters (see sensitivity analysis of the ForCLIM model by Morin et al. 2011 and Huber et al. 2018).

More generally about long term predictions, reaching stronger robustness in predicting long-term species coexistence and community composition would necessitate to better model the occurrence of mortality events and regeneration. In fact, improving the representation of these two processes is a main challenge in forest modelling, especially to better assess climate change impacts on forest functioning (e.g. for mortality Bugmann et al. 2019, Cailleret et al. 2017, Hülsmann et al. 2018, Vanoni et al. 2019). Besides, although nutrients and water content in the soil indirectly affects competition between trees (see Methods section), future developments may lead to a multi-dimensional competition along several niche axes. One may also notice that the results for the two Mediterranean sites presented here are already satisfying. Furthermore, the impacts of abiotic (e.g., fire, extreme drought events) and biotic (e.g., pathogens, herbivory) disturbances are also key factors, that should be better considered by these models in the future (Seidl et al. 2017).

### Mechanistic relevance of ForCEEPS parameters

The analysis exploring the congruence between key ForCEEPS’ parameters and functional traits retrieved from the literature aimed at highlighting to what extent the parameters describing species in ForCEEPS can be linked to their ecophysiology. First the negative correlation between the growth parameter (*g*_*S*_) and wood density appears meaningful as wood density describes the carbon investment per unit volume of stem (Chaves et al 2009), thus indicating that fast-growing species favored wood volume (i.e., space exploration) at the expense of wood resistance to mechanical or biotic damages.

Shade tolerance is one of the features that segregates ecological groups of tree species and that explain BEF patterns in forests. Some studies indicate that shade tolerance is related to a combination of structural properties maximizing leaf area per unit of respiring biomass, and to a combination of leaf properties optimizing photosynthesis per unit of nitrogen investment. In particular shade-intolerant or pioneer species are frequently thought to display higher light-saturated net photosynthesis (*Amax*) than shade-tolerant or late successional species (Coste et al., 2005; Reich & Walters, 1994). Consistent with this later assertion, we found a significant and consistent correlation between *ShTol_seedling*_*S*_ and *A*_*max*_ (Fig. 5), and to a lower extent between *ShTol*_*S*_ and *A*_*max*_ (Table S5). However no correlation was found with *LMA*, which echoes the debate regarding the multiple factors influencing this trait - including ontogeny, leaf life span, and light environment - that can blur any expected pattern (Lusk & Warton, 2007).

Drought tolerance (*DrTol*_*S*_) is another key parameter that was positively correlated with a number of functional traits (Table S5, Fig. 5). The best correlation, however, was found with species embolism resistance (assessed through the water potential causing 50% loss of conductivity, *P50*). This pattern is consistent with current ecophysiological knowledge that xylem embolism is a key driver of species mortality during drought (Martin-StPaul et al 2017; Adams et al 2018). Additionally, a significant but weaker correlation was found between *DrTol*_*S*_ and the turgor loss point – a trait linked to the maintenance of leaf hydration and functions at low water potential (Bartlett, Scoffoni, & Sack, 2012) and to stomatal control (Brodribb & Holbrook, 2003; Martin-StPaul et al., 2017). This lower correlation is consistent with the fact that the variability of turgor loss point is much more constrained among plants than the *P50* (Martin-StPaul et al 2017). Interestingly, as for *Sh_tol*_*S*_, *DrTol*_*S*_ was only weakly correlated with wood density and *LMA*, which is probably related to their poor mechanistic relevance in the species resistance to drought (Chave et al 2009 ; Bartlett et al 2012). Although performed on a relatively small number of species, these results nevertheless pave the way for potential improvement of the representation of drought tolerance in ForCEEPS, for instance by implementing an hydraulic failure module that mechanistically integrate multiple traits (e.g., Martin-StPaul et al 2017). More generally, exploring the mechanistic relevance of gap model parameters allows using functional trait databases to constrain them within realistic values and avoid equifinalities issues.

### Research avenues for a new generation of forest gap models

The large potential of forest dynamic models to tackle key questions in forest ecology has been reviewed elsewhere (Ruiz-Benito et al. 2020), but we highlight that their role in providing more robust predictions in response to global change components is increasingly emphasized (McDowell 2020). Furthermore, we would like to focus on two related perspectives that are arising from the validation at both short and long term shown here.

#### Biodiversity and ecosystem functioning in forests

The validation presented here opens perspectives for further tests of the effects of species richness or functional diversity on forest productivity. Several attempts were conducted to use gap models for studying diversity-productivity relationships (Morin et al. 2011, Bohn et al. 2017). Nevertheless, the models used had not been validated rigorously for monospecific forests across such a wide range of species and environmental conditions, although the analyses about the effect of diversity on ecosystem functioning strongly rely on the comparison with monospecific stands (Loreau and Hector 2001, 2019). More precisely, the increased confidence in the ability of gap models to simulate monospecific stands will improve their ability to test non-additive effects in species mixtures (Gamfeldt and Roger 2017), i.e., effects directly related to interspecific interactions. Furthermore, as ForCEEPS accurately predicts stand productivity and long-term composition for the main species in Western Europe under a wide range of conditions, we may expect a high robustness of the simulated BEF relationship.

Forest gap models simulate local interactions among trees, which have been reported as fundamental drivers of mixture effects on forest functioning (Fichtner et al. 2018). Thus, the simulated biodiversity patterns necessarily emerge from selection and complementarity effects (Loreau and Hector 2001), the latter referring to niche differentiation processes among co-existing species (as detailed in Morin et al. 2011) but also facilitative processes, depending on the model structure. Niche differentiation processes notably include complementarity in occupying canopy space (Jucker et al. 2015, Williams et al. 2017), and the 2-D crown representation of ForCEEPS enables to better explore the way canopy packing occurs in simulated mixtures and affects forest productivity. More generally, a growing body of evidence suggests that structural diversity is a key driver of productivity in forests, independently of the potential effects of other facets of diversity such as species richness and functional diversity (Dănescu et al. 2016, Schnabel et al. 2019, Gough et al. 2019, Aponte et al. 2020). ForCEEPS is a suitable tool to quantify the importance of these - often tangled - diversity facets across large environmental gradients, with important consequence for our understating of BEF relationships and for the management of diverse forests.

Furthermore, plasticity in crown size (or more precisely tree foliage area) may emerge in the simulations due to species complementarity in light capture and/or response to environmental fluctuations (e.g., climate). Intraspecific changes in crown architecture are ultimately determined by changes in within-tree biomass allocation and branching patterns, which have been shown to occur in mixed stands (Pretzsch 2014, Kunz et al. 2019, Guillemot et al. 2020) but are not considered here. The modelling of such mixture effects is currently hindered by data scarcity, and would probably necessitate implementing the spatial distribution of the simulated trees in the horizontal space (Forrester et al. 2018).

#### Testing coexistence mechanisms in the short and long term

Species coexistence in forest gap models is based on two main mechanisms: first, trade-offs arising from the life-history strategies such as high rates of colonization often being tied to low shade tolerance, or a typically short lifespan of early successional, fast-growing trees; and second, the fact that cyclical succession is occurring on each individual patch, so that species with different properties are able to dominate during different parts of the cycle (Bazzaz 1979, Rees et al. 2001). Exploring the relative importance of these mechanisms for allowing species coexistence of simulated communities, but also for creating and maintaining diversity effects on ecosystem functioning is a promising avenue for gap model applications (Falster et al. 2017, Cordonnier et al. 2018b), especially if such an exploration is to be carried out across a large range of conditions. This may ultimately lead to the formulation of new hypotheses, for instance about the impact of climate change on species coexistence and forest functioning.

Finally, we also see further potential applications of models like ForCEEPS in the design of forest policy. Large-scale forest restoration and reforestation programs are key to prevent the most deleterious effects of climate change in the coming decade (Lewis et al. 2019). Global initiatives such as the Bonn challenge are planning restoration at an unprecedented scale (Verdone and Seidl 2017). Yet, we currently lack science-based guidelines for the design of productive and resilient forest plantations in most environmental contexts. As mixed-species plantations are thought to be a crucial nature-based solution for climate mitigation and adaptation (Paquette et al. 2018), a generic and validated tool such as ForCEEPS can be used to explore “management versus climate scenario” interactions and promote climate-smart forestry at large scale. Thus, a new generation of forest gap models could foster the transfer of BEF knowledge into forestry practice.

Generating new hypotheses from model outcomes is one of the main reasons of using models in ecology in the first place, together with the support they may provide for better understanding the systems and processes at play, and their ability to yield predictions across spatial and temporal scales (Levins 1966). As they did for more than 50 years, we believe that gap models in general, and the ForCEEPS model presented here in particular, maintain a key role for these purposes in forest ecology and management. More generally, because they seek for generality while aiming at relying on functional processes, such models are likely to be highly relevant to provide robust predictions of ecosystem composition, structure and functioning in a context of very uncertain future for forests (McDowell et al. 2020).

## Appendices of Morin et al

Appendix A. Description of the model ForCEEPS.

Appendix B. Supplementary Tables.

S1. Parameter values for all species

S2. Number of plots and trees per species considered in the analyses

S3. Sites used in the PNV analysis

S4. All repetitions – tree level

S5. Correlations between ForCEEPS parameters and ecological traits at the interspecific level

Appendix C. Supplementary Figures.

S1. Representation of tree crowns in ForCEEPS

S2. Distribution of the sites according to climate variables or indices

S3. Repetitions at the tree level

S4. Results at the tree level per species

S5. Results at the stand level per species

S6. Residuals of simulated tree growth against tree diameter

S7. Residuals of simulated stand growth against stand basal area

Appendix D. Additional information about field data for calibration.

Appendix E. Information about ecophysiological traits

## Acknowledgements

The authors wish to thank the Institut Géographique National for providing the national forest inventory data, and the Office National des Forêts and the RENECOFOR team, particularly Manuel Nicolas and Marc Lanier, for providing their database. In both cases, these long-term monitoring networks are invaluable tools. The SAFRAN database was provided by Météo-France. The authors are also grateful to the CAPSIS modelling platform (http://www7.inra.fr/capsis/). They thank Maude Toigo, Georges Kunstler and Patrick Vallet for helpful discussions about the study, and Gregor Cresnar from the Noun Project. This study was funded by the projects BIOPROFOR (ANR-11-PDOC-030-01), DISTIMACC (ECOFOR-2014-23), French Ministry of Ecology and Sustainable Development, French Ministry of Agriculture and Forest) and DIPTICC (ANR-16-CE32-0003).

## Data availability statement

The observed and simulated data that support the findings of the study will be deposited in Figshare.

## References

Allen, C. D., A. K. Macalady, H. Chenchouni, D. Bachelet, N. McDowell, M. Vennetier, T. Kitzberger, A. Rigling, D. D. Breshears, E. H. Hogg, P. Gonzalez, R. Fensham, Z. Zhang, J. Castro, N. Demidova, J. H. Lim, G. Allard, S. W. Running, A. Semerci, and N. Cobb. 2010. A global overview of drought and heat-induced tree mortality reveals emerging climate change risks for forests. Forest Ecology and Management 259:660–684.

Aponte, C., S. Kasel, C. R. Nitschke, M. A. Tanase, H. Vickers, L. Parker, M. Fedrigo, M. Kohout, P. Ruiz-Benito, M. A. Zavala, and L. T. Bennett. 2020. Structural diversity underpins carbon storage in Australian temperate forests. Global Ecology and Biogeography:geb.13038.

Bartlett, M. K., Scoffoni, C., and Sack, L. 2012. The determinants of leaf turgor loss point and prediction of drought tolerance of species and biomes: a global meta-analysis. Ecology Letters, 15:393–405. doi: 10.1111/j.1461-0248.2012.01751.x

Bayer, D., S. Seifert, and H. Pretzsch. 2013. Structural crown properties of Norway spruce (Picea abies [L.] Karst.) and European beech (Fagus sylvatica [L.]) in mixed versus pure stands revealed by terrestrial laser scanning. Trees-Structure and Function 27:1035–1047.

Bazzaz, F. A. 1979. Physiological Ecology of Plant Succession. Annual Review of Ecology and Systematics 10:351–371.

Bohn, F. J., K. Frank, and A. Huth. 2014. Of climate and its resulting tree growth: Simulating the productivity of temperate forests. Ecological Modelling 278:9–17.

Bohn, F. J., and A. Huth. 2017. The importance of forest structure to biodiversity– productivity relationships. Royal Society Open Science 4:160521

Boisvenue, C., and S. W. Running. 2006. Impacts of climate change on natural forest productivity – evidence since the middle of the 20th century. Global Change Biology 12:862–882.

Brodribb, T. J., and Holbrook, N. M. 2003. Stomatal closure during leaf dehydration, correlation with other leaf physiological traits. Plant Physiology, 132:2166–2173. doi: 10.1104/pp.103.023879

Botkin, D. B., J. F. Janak, and J. R. Wallis. 1972. Some ecological consequences of a computer model of forest growth. Journal of Ecology 60:849–872.

Bugmann, H. 1994. On the Ecology of mountainous forests in a changing climate: A simulation study. Eidgenössische Technische Hochschule, Zürich.

Bugmann, H. 1996. A simplified forest model to study species composition along climate gradients. Ecology 77:2055–2074.

Bugmann, H. 2001. A review of forest gap models. Climatic Change 51:259–305.

Bugmann, H., R. Seidl, F. Hartig, F. Bohn, J. Bruna, M. Cailleret, L. François, J. Heinke, A.-J. Henrot, T. Hickler, L. Hülsmann, A. Huth, I. Jacquemin, C. Kollas, P. Lasch-Born, M. J. Lexer, J. Merganič, K. Merganičová, T. Mette, B. R. Miranda, D. Nadal-Sala, W. Rammer, A. Rammig, B. Reineking, E. Roedig, S. Sabaté, J. Steinkamp, F. Suckow, G. Vacchiano, J. Wild, C. Xu, and C. P. O. Reyer. 2019. Tree mortality submodels drive simulated long-term forest dynamics: assessing 15 models from the stand to global scale. Ecosphere 10:e02616.

Bugmann, H., and A. M. Solomon. 2000. Explaining forest composition and biomass across multiple biogeographical regions. Ecological Applications 10:95–114.

Cailleret, M., S. Jansen, E. M. R. Robert, L. Desoto, T. Aakala, J. A. Antos, B. Beikircher, C. Bigler, H. Bugmann, M. Caccianiga, V. Čada, J. J. Camarero, P. Cherubini, H. Cochard, M. R. Coyea, K. Čufar, A. J. Das, H. Davi, S. Delzon, M. Dorman, G. Gea-Izquierdo, S. Gillner, L. J. Haavik, H. Hartmann, A. Hereş, K. R. Hultine, P. Janda, J. M. Kane, V. I. Kharuk, T. Kitzberger, T. Klein, K. Kramer, F. Lens, T. Levanic, J. C. Linares Calderon, F. Lloret, R. Lobo-Do-Vale, F. Lombardi, R. López Rodríguez, H. Mäkinen, S. Mayr, I. Mészáros, J. M. Metsaranta, F. Minunno, W. Oberhuber, A. Papadopoulos, M. Peltoniemi, A. M. Petritan, B. Rohner, G. Sangüesa-Barreda, D. Sarris, J. M. Smith, A. B. Stan, F. Sterck, D. B. Stojanović, M. L. Suarez, M. Svoboda, R. Tognetti, J. M. Torres-Ruiz, V. Trotsiuk, R. Villalba, F. Vodde, A. R. Westwood, P. H. Wyckoff, N. Zafirov, and J. Martínez-Vilalta. 2017. A synthesis of radial growth patterns preceding tree mortality. Global Change Biology 23:1675–1690.

Canham, C. D., A. C. Finzi, S. W. Pacala, and D. H. Burbank. 1994. Causes and consequences of resource heterogeneity in forests: Interspecific variation in light transmission by canopy trees. Canadian Journal of Forest Research 24:337–349.

Castagneyrol, B., B. Giffard, C. Péré, and H. Jactel. 2013. Plant apparency, an overlooked driver of associational resistance to insect herbivory. Journal of Ecology 101:418–429.

Chapin, F. S., J. T. Randerson, A. D. McGuire, J. A. Foley, and C. B. Field. 2008. Changing feedbacks in the climate-biosphere system. Frontiers in Ecology and the Environment 6:313–320.

Charru, M., I. Seynave, F. Morneau, and J.-D. Bontemps. 2010. Recent changes in forest productivity: An analysis of national forest inventory data for common beech (Fagus sylvatica L.) in north-eastern France. Forest Ecology and Management 260:864–874.

Chauvet, M., G. Kunstler, J. Roy, and X. Morin. 2017. Using a forest dynamics model to link community assembly processes and traits structure. Functional Ecology in press.

Chave, J., Coomes, D., Jansen, S., Lewis, S. L., Swenson, N. G., and Zanne, A. E. 2009. Towards a worldwide wood economics spectrum. Ecology Letters, 12:351–366. doi: 10.1111/j.1461-0248.2009.01285.x

Coomes, D. A., O. Flores, R. Holdaway, T. Jucker, E. R. Lines, and M. C. Vanderwel. 2014. Wood production response to climate change will depend critically on forest composition and structure. Global Change Biology 20:3632–3645.

Cordonnier, T., T. Bourdier, G. Kunstler, C. Piedallu, and B. Courbaud. 2018a. Covariation between tree size and shade tolerance modulates mixed-forest productivity. Annals of Forest Science 75:101.

Cordonnier, T., G. Kunstler, B. Courbaud, and X. Morin. 2018b. Managing tree species diversity and ecosystem functioning through coexistence mechanisms. Annals of Forest Sciences.

Cordonnier, T., C. Smadi, G. Kunstler, and B. Courbaud. 2019. Asymmetric competition, ontogenetic growth and size inequality drive the difference in productivity between two-strata and one-stratum forest stands. Theoretical Population Biology.

Coste, S., Roggy, J. C., Imbert, P., Born, C., Bonal, D., and Dreyer, E. 2005. Leaf photosynthetic traits of 14 tropical rain forest species in relation to leaf nitrogen concentration and shade tolerance. Tree Physiology, 25:1127–1137. doi: 10.1093/treephys/25.9.1127

Danescu, A., A. T. Albrecht, and J. Bauhus. 2016. Structural diversity promotes productivity of mixed, uneven-aged forests in southwestern Germany. Oecologia 182:319–333.

DeSoto, L., M. Cailleret, F. Sterck, S. Jansen, K. Kramer, E. M. R. Robert, T. Aakala, M. M. Amoroso, C. Bigler, J. J. Camarero, K. Čufar, G. Gea-Izquierdo, S. Gillner, L. J. Haavik, A. M. Hereş, J. M. Kane, V. I. Kharuk, T. Kitzberger, T. Klein, T. Levanič, J. C. Linares, H. Mäkinen, W. Oberhuber, A. Papadopoulos, B. Rohner, G. Sangüesa-Barreda, D. B. Stojanovic, M. L. Suárez, R. Villalba, and J. Martínez-Vilalta. 2020. Low growth resilience to drought is related to future mortality risk in trees. Nature Communications 11:1–9.

Didion, M., A. D. Kupferschmid, A. Wolf, and H. Bugmann. 2011. Ungulate herbivory modifies the effects of climate change on mountain forests. Climatic Change 109:647–669.

Didion, M., A. D. Kupferschmid, A. Zingg, L. Fahse, and H. Bugmann. 2009. Gaining local accuracy while not losing generality — extending the range of gap model applications. Canadian Journal of Forest Research 39:1092–1107.

Dobbertin, M. 2005. Tree growth as indicator of tree vitality and of tree reaction to environmental stress: a review. European Jounral of Forest Research 124:319–333.

Dufrêne, E., H. Davi, C. François, G. le Maire, V. Le Dantec, and A. Granier. 2005. Modelling carbon and water cycles in a beech forest: Part I: Model description and uncertainty analysis on modelled NEE. Ecological Modelling 185:407–436.

Eermak, J. 1998. Leaf distribution in large trees and stands of the floodplain forest in southern Moravia. Tree Physiology 18:727–737.

Ellenberg, H., and D. Mueller-Dombois. 1966. Tentative physiognomic-ecological classification of plant formations of the Earth. Ber. Geobot. Inst. ETH 37:21–55

Esquivel-Muelbert, A., T. R. Baker, K. G. Dexter, S. L. Lewis, R. J. W. Brienen, T. R. Feldpausch, J. Lloyd, A. Monteagudo-Mendoza, L. Arroyo, E. Álvarez-Dávila, N. Higuchi, B. S. Marimon, B. H. Marimon-Junior, M. Silveira, E. Vilanova, E. Gloor, Y. Malhi, J. Chave, J. Barlow, D. Bonal, N. Davila Cardozo, T. Erwin, S. Fauset, B. Hérault, S. Laurance, L. Poorter, L. Qie, C. Stahl, M. J. P. Sullivan, H. ter Steege, V. A. Vos, P. A. Zuidema, E. Almeida, E. Almeida de Oliveira, A. Andrade, S. A. Vieira, L. Aragão, A. Araujo-Murakami, E. Arets, G. A. Aymard C, C. Baraloto, P. B. Camargo, J. G. Barroso, F. Bongers, R. Boot, J. L. Camargo, W. Castro, V. Chama Moscoso, J. Comiskey, F. Cornejo Valverde, A. C. Lola da Costa, J. del Aguila Pasquel, A. Di Fiore, L. Fernanda Duque, F. Elias, J. Engel, G. Flores Llampazo, D. Galbraith, R. Herrera Fernández, E. Honorio Coronado, W. Hubau, E. Jimenez-Rojas, A. J. N. Lima, R. K. Umetsu, W. Laurance, G. Lopez-Gonzalez, T. Lovejoy, O. Aurelio Melo Cruz, P. S. Morandi, D. Neill, P. Núñez Vargas, N. C. Pallqui Camacho, A. Parada Gutierrez, G. Pardo, J. Peacock, M. Peña-Claros, M. C. Peñuela-Mora, P. Petronelli, G. C. Pickavance, N. Pitman, A. Prieto, C. Quesada, H. Ramírez-Angulo, M. Réjou-Méchain, Z. Restrepo Correa, A. Roopsind, A. Rudas, R. Salomão, N. Silva, J. Silva Espejo, J. Singh, J. Stropp, J. Terborgh, R. Thomas, M. Toledo, A. Torres-Lezama, L. Valenzuela Gamarra, P. J. van de Meer, G. van der Heijden, P. van der Hout, R. Vasquez Martinez, C. Vela, I. C. G. Vieira, and O. L. Phillips. 2019. Compositional response of Amazon forests to climate change. Global Change Biology 25:39–56.

Falster, D. S., Å. Brännström, M. Westoby, and U. Dieckmann. 2017. Multitrait successional forest dynamics enable diverse competitive coexistence. Proceedings of the National Academy of Sciences of the United States of America 114:E2719–E2728.

Falster, D. S., Duursma, R. A., and FitzJohn, R. G. 2018. How functional traits influence plant growth and shade tolerance across the life cycle. Proceedings of the National Academy of Sciences 115: E6789–E6798.

Fichtner, A., W. Härdtle, H. Bruelheide, M. Kunz, Y. Li, and Goddert Von Oheimb. 2018. Neighbourhood interactions drive overyielding in mixed-species tree communities. Nature Communications 9:1144.

Forrester, D. I., and A. T. Albrecht. 2014. Light absorption and light-use efficiency in mixtures of Abies alba and Picea abies along a productivity gradient. Forest Ecology and Management 328:94–102.

Forrester, D. I., C. Ammer, P. J. Annighöfer, I. Barbeito, K. Bielak, A. Bravo-Oviedo, L. Coll, M. del Río, L. Drössler, M. Heym, V. Hurt, M. Löf, J. den Ouden, M. Pach, M. G. Pereira, B. N. E. Plaga, Q. Ponette, J. Skrzyszewski, H. Sterba, M. Svoboda, T. M. Zlatanov, and H. Pretzsch. 2018. Effects of crown architecture and stand structure on light absorption in mixed and monospecific *Fagus sylvatica* and *Pinus sylvestris* forests along a productivity and climate gradient through Europe. Journal of Ecology 106:746–760.

Fortin, M., R. Van Couwenberghe, V. Perez, and C. Piedallu. 2019. Evidence of climate effects on the height-diameter relationships of tree species. Annals of Forest Science 76:1.

Fyllas, N. M., E. Gloor, L. M. Mercado, S. Sitch, C. A. Quesada, T. F. Domingues, D. R. Galbraith, A. Torre-Lezama, E. Vilanova, H. Ramírez-Angulo, N. Higuchi, D. A. Neill, M. Silveira, L. Ferreira, G. A. Aymard C., Y. Malhi, O. L. Phillips, and J. Lloyd. 2014. Analysing Amazonian forest productivity using a new individual and trait-based model (TFS v.1). Geoscientific Model Development 7:1251–1269.

Gamfeldt, L., and F. Roger. 2017, June 22. Revisiting the biodiversity-ecosystem multifunctionality relationship. Nature Publishing Group.

Gough, C. M., J. W. Atkins, R. T. Fahey, and B. S. Hardiman. 2019. High rates of primary production in structurally complex forests. Ecology 100.

Guillemot, J., M. Kunz, F. Schnabel, A. Fichtner, C. P. Madsen, T. Gebauer, W. Härdtle, G. von Oheimb, and C. Potvin. 2020. Neighbourhood-mediated shifts in tree biomass allocation drive overyielding in tropical species mixtures. New Phytologist:nph.16722.

Guillemot, J., C. Francois, G. Hmimina, E. Dufrêne, N. K. Martin-StPaul, K. Soudani, G. Marie, J.-M. Ourcival, and N. Delpierre. 2017. Environmental control of carbon allocation matters for modelling forest growth. New Phytologist 214:180–193.

Hardiman, B. S., G. Bohrer, C. M. Gough, C. S. Vogel, and P. S. Curtis. 2011. The role of canopy structural complexity in wood net primary production of a maturing northern deciduous forest. Ecology 92:1818–1827.

Hartig, F., J. Dyke, T. Hickler, S. I. Higgins, R. B. O’Hara, S. Scheiter, and A. Huth. 2012. Connecting dynamic vegetation models to data - an inverse perspective. Journal of Biogeography 39:2240–2252.

Huber, N., H. Bugmann, and V. Lafond. 2018. Global sensitivity analysis of a dynamic vegetation model: Model sensitivity depends on successional time, climate and competitive interactions. Ecological Modelling 368:377–390.

Hülsmann, L., H. Bugmann, M. Cailleret, and P. Brang. 2018. How to kill a tree: empirical mortality models for 18 species and their performance in a dynamic forest model. Ecological Applications 28:522–540.

IFN. 2016. Les Résultats Issus des Campagnes d’Inventaire entre 2011 et 2016. Nogent-sur-Vernisson.

IGN. 2018. Données brutes de l’Inventaire forestier national. https://inventaire-forestier.ign.fr/spip.php?rubrique159.

Jactel, H., E. S. Gritti, L. Drössler, D. I. Forrester, W. L. Mason, X. Morin, H. Pretzsch, and B. Castagneyrol. 2018. Positive biodiversity-productivity relationships in forests: climate matters. Biology Letters 14:1–4.

Jourdan, M., G. Kunstler, and X. Morin. 2019a. How neighbourhood interactions control the temporal stability and resilience to drought of trees in mountain forests. Journal of Ecology:1365-2745.13294.

Jourdan, M., F. Lebourgeois, and X. Morin. 2019b. The effect of tree diversity on the resistance and recovery of forest stands in the French Alps may depend on species differences in hydraulic features. Forest Ecology and Management 450:117486.

Jucker, T., D. Avacari?ei, I. Bărnoaiea, G. Duduman, O. Bouriaud, and D. A. Coomes. 2016. Climate modulates the effects of tree diversity on forest productivity. Journal of Ecology 104:388–398.

Jucker, T., O. Bouriaud, and D. A. Coomes. 2015. Crown plasticity enables trees to optimize canopy packing in mixed-species forests. Functional Ecology 29.

Kattge, J., S. Diaz, S. Lavorel, C. Prentice, P. Leadley, G. Bonisch, E. Garnier, M. Westoby, P. B. Reich, I. J. Wright, J. H. C. Cornelissen, C. Violle, S. P. Harrison, P. M. van Bodegom, M. Reichstein, B. J. Enquist, N. A. Soudzilovskaia, D. D. Ackerly, M. Anand, O. Atkin, M. Bahn, T. R. Baker, D. Baldocchi, R. Bekker, C. C. Blanco, B. Blonder, W. J. Bond, R. Bradstock, D. E. Bunker, F. Casanoves, J. Cavender-Bares, J. Q. Chambers, F. S. Chapin, J. Chave, D. Coomes, W. K. Cornwell, J. M. Craine, B. H. Dobrin, L. Duarte, W. Durka, J. Elser, G. Esser, M. Estiarte, W. F. Fagan, J. Fang, F. Fernandez- Mendez, A. Fidelis, B. Finegan, O. Flores, H. Ford, D. Frank, G. T. Freschet, N. M. Fyllas, R. V Gallagher, W. A. Green, A. G. Gutierrez, T. Hickler, S. I. Higgins, J. G. Hodgson, A. Jalili, S. Jansen, C. A. Joly, A. J. Kerkhoff, D. Kirkup, K. Kitajima, M. Kleyer, S. Klotz, J. M. H. Knops, K. Kramer, I. Kuhn, H. Kurokawa, D. Laughlin, T. D. Lee, M. Leishman, F. Lens, T. Lenz, S. L. Lewis, J. Lloyd, J. Llusia, F. Louault, S. Ma, M. D. Mahecha, P. Manning, T. Massad, B. E. Medlyn, J. Messier, A. T. Moles, S. C. Muller, K. Nadrowski, S. Naeem, U. Niinemets, S. Nollert, A. Nuske, R. Ogaya, J. Oleksyn, V. G. Onipchenko, Y. Onoda, J. Ordonez, G. Overbeck, W. A. Ozinga, S. Patino, S. Paula, J. G. Pausas, J. Penuelas, O. L. Phillips, V. Pillar, H. Poorter, L. Poorter, P. Poschlod, A. Prinzing, R. Proulx, A. Rammig, S. Reinsch, B. Reu, L. Sack, B. Salgado-Negre, J. Sardans, S. Shiodera, B. Shipley, A. Siefert, E. Sosinski, J. F. Soussana, E. Swaine, N. Swenson, K. Thompson, P. Thornton, M. Waldram, E. Weiher, M. White, S. White, S. J. Wright, B. Yguel, S. Zaehle, A. E. Zanne, and C. Wirth. 2011. TRY - a global database of plant traits. Global Change Biology 17:2905–2935.

Keenan, T. F., and Niinemets, Ü. 2016. Global leaf trait estimates biased due to plasticity in the shade. Nature Plants, 3:1–6. doi: 10.1038/nplants.2016.201

Korner, C., R. Asshoff, O. Bignucolo, S. Hättenschwiler, S. G. Keel, S. Peláez-Riedl, S. Pepin, R. T. W. Siegwolf, and G. Zotz. 2005. Carbon flux and growth in mature deciduous forest trees exposed to elevated CO2. Science 309:1360–1362.

Kunz, M., A. Fichtner, W. Härdtle, P. Raumonen, H. Bruelheide, and G. von Oheimb. 2019, December 1. Neighbour species richness and local structural variability modulate aboveground allocation patterns and crown morphology of individual trees. Blackwell Publishing Ltd.

Levins, R. 1966. The strategy of model building in population ecology. American Scientist 54:421–451.

Lewis, S. L., C. E. Wheeler, E. T. A. Mitchard, and A. Koch. 2019. Restoring natural forests is the best way to remove atmospheric carbon. Nature 568:25–28.

Liang, J., T. W. Crowther, N. Picard, S. Wiser, M. Zhou, G. Alberti, E.-D. Schulze, A. D. McGuire, F. Bozzato, H. Pretzsch, S. de-Miguel, A. Paquette, B. Hérault, M. Scherer- Lorenzen, C. B. Barrett, H. B. Glick, G. M. Hengeveld, G.-J. Nabuurs, S. Pfautsch, H. Viana, A. C. Vibrans, C. Ammer, P. Schall, D. Verbyla, N. Tchebakova, M. Fischer, J. V Watson, H. Y. H. Chen, X. Lei, M.-J. Schelhaas, H. Lu, D. Gianelle, E. I. Parfenova, C. Salas, E. Lee, B. Lee, H. S. Kim, H. Bruelheide, D. A. Coomes, D. Piotto, T. Sunderland, B. Schmid, S. Gourlet-Fleury, B. Sonké, R. Tavani, J. Zhu, S. Brandl, J. Vayreda, F. Kitahara, E. B. Searle, V. J. Neldner, M. R. Ngugi, C. Baraloto, L. Frizzera, R. Bałazy, J. Oleksyn, T. Zawiła-Niedzwiecki, O. Bouriaud, F. Bussotti, L. Finér, B. Jaroszewicz, T. Jucker, F. Valladares, A. M. Jagodzinski, P. L. Peri, C. Gonmadje, W. Marthy, T. O’Brien, E. H. Martin, A. R. Marshall, F. Rovero, R. Bitariho, P. A. Niklaus, P. Alvarez-Loayza, N. Chamuya, R. Valencia, F. Mortier, V. Wortel, N. L. Engone-Obiang, L. V Ferreira, D. E. Odeke, R. M. Vasquez, S. L. Lewis, and P. B. Reich. 2016. Positive biodiversity-productivity relationship predominant in global forests. Science 354.

Limousin, J. M., S. Rambal, J. M. Ourcival, A. Rocheteau, R. Joffre, and R. Rodriguez- Cortina. 2009. Long-term transpiration change with rainfall decline in a Mediterranean Quercus ilex forest. Global Change Biology 15:2163–2175.

Lindner, M., M. Maroschek, S. Netherer, A. Kremer, A. Barbati, J. Garcia-Gonzalo, R. Seidl, S. Delzon, P. Corona, M. Kolström, M. J. Lexer, and M. Marchetti. 2010. Climate change impacts, adaptive capacity, and vulnerability of European forest ecosystems. Forest Ecology and Management 259:698–709.

Loreau, M., and A. Hector. 2001. Partitioning selection and complementarity in biodiversity experiments. Nature 412:72–76.

Loreau, M., and A. Hector. 2019. Not even wrong: Comment by Loreau and Hector. Ecology.

Lusk, C. H., and Warton, D. I. 2007. Global meta-analysis shows that relationships of leaf mass per area with species shade tolerance depend on leaf habit and ontogeny. New Phytologist, 176:764–774. doi: 10.1111/j.1469-8137.2007.02264.x

Makela, A., J. Landsberg, A. R. Ek, T. E. Burk, M. Ter-Mikaelian, G. I. Agren, C. D. Oliver, and P. Puttonen. 2000. Process-based models for forest ecosystem management: current state of the art and challenges for practical implementation. Tree Physiology 20:289–298.

Maréchaux, I., and J. Chave. 2017. An individual-based forest model to jointly simulate carbon and tree diversity in Amazonia: description and applications. Ecological Monographs 87:632–664.

Martin-StPaul, N., S. Delzon, and H. Cochard. 2017. Plant resistance to drought depends on timely stomatal closure. Ecology Letters 20:1437–1447. doi: 10.1111/ele.12851

McDowell, N. G., C. D. Allen, K. Anderson-Teixeira, B. H. Aukema, B. Bond-Lamberty, L. Chini, J. S. Clark, M. Dietze, C. Grossiord, A. Hanbury-Brown, G. C. Hurtt, R. B. Jackson, D. J. Johnson, L. Kueppers, J. W. Lichstein, K. Ogle, B. Poulter, T. A. M. Pugh, R. Seidl, M. G. Turner, M. Uriarte, A. P. Walker, and C. Xu. 2020. Pervasive shifts in forest dynamics in a changing world. Science 368:eaaz9463.

Mette, T., A. Albrecht, C. Ammer, P. Biber, U. Kohnle, and H. Pretzsch. 2009. Evaluation of the forest growth simulator SILVA on dominant trees in mature mixed Silver fir– Norway spruce stands in South-West Germany. Ecological Modelling 220:1670–1680.

Mina, M., H. Bugmann, T. Cordonnier, F. Irauschek, M. Klopcic, M. Pardos, and M. Cailleret. 2017. Future ecosystem services from European mountain forests under climate change. Journal of Applied Ecology 54:389–401.

Mitscherlich, G., and K. von Gadow. 1968. Über den zuwachsverlust bei der ästung von nabelbäumen. Allgemaine Forst- und Jagdzeitung.

Moore, A. D. 1989. On the maximum growth equation used in forest gap simulation models. Ecological Modelling 45:63–67.

Morin, X., L. Fahse, H. Jactel, M. Scherer-Lorenzen, R. García-Valdés, and H. Bugmann. 2018. Long-term response of forest productivity to climate change is mostly driven by change in tree species composition. Scientific Reports 8:5627.

Morin, X., L. Fahse, M. Scherer-Lorenzen, and H. Bugmann. 2011. Tree species richness promotes productivity in temperate forests through strong complementarity between species. Ecology Letters 14:1211–1219.

Nadrowski, K., C. Wirth, and M. Scherer-Lorenzen. 2010. Is forest diversity driving ecosystem function and service? Current Opinion in Environmental Sustainability 2:75–79.

Nehrbass-Ahles, C., Babst, F., Klesse, S., Nötzli, M., Bouriaud, O., Neukom, R., …, and Frank, D. 2014. The influence of sampling design on tree-ring-based quantification of forest growth. Global Change Biology 20:2867–2885.

Niklaus, P. A., M. Baruffol, J.-S. He, K. Ma, and B. Schmid. 2017. Can niche plasticity promote biodiversity-productivity relationships through increased complementarity? Ecology 98:1104–1116.

Niinemets, U., and F. Valladares. 2006. Tolerance to shade, drought and waterlogging of temperate, Northern hemisphere trees and shrubs. Ecological Monographs 76:521–547.

Norby, R. J., and D. R. Zak. 2011. Ecological Lessons from Free-Air CO _2_ Enrichment (FACE) Experiments. Annual Review of Ecology, Evolution, and Systematics 42:181–203.

Pacala, S. W., C. D. Canham, and J. A. Silander. 1993. Forest Models Defined by Field-Measurements .1. the Design of a Northeastern Forest Simulator. Canadian Journal of Forest Research-Revue Canadienne De Recherche Forestiere 23:1980–1988.

Pan, Y., R. A. Birdsey, J. Fang, R. Houghton, P. E. Kauppi, W. A. Kurz, O. L. Phillips, A. Shvidenko, S. L. Lewis, J. G. Canadell, P. Ciais, R. B. Jackson, S. W. Pacala, A. D. McGuire, S. Piao, A. Rautiainen, S. Sitch, and D. Hayes. 2011. A large and persistent carbon sink in the world’s forests. Science (New York, N.Y.) 333:988–93.

Paquette, A., A. Hector, B. Castagneyrol, M. Vanhellemont, J. Koricheva, M. Scherer- Lorenzen, and K. Verheyen. 2018. A million and more trees for science. Nature Ecology & Evolution 2:763–766.

Van de Peer, T., K. Verheyen, Q. Ponette, N. N. Setiawan, and B. Muys. 2018. Overyielding in young tree plantations is driven by local complementarity and selection effects related to shade tolerance. Journal of Ecology 106:1096–1105.

Van Pelt, R., S. C. Sillett, W. A. Kruse, J. A. Freund, and R. D. Kramer. 2016. Emergent crowns and light-use complementarity lead to global maximum biomass and leaf area in Sequoia sempervirens forests. Forest Ecology and Management 375:279–308.

Pfister, C., and H. Bugmann. 2000. Impacts of interannual climate variability on past and future forest composition. Regional Environmental Change 1:112–125.

Poorter, L., L. Bongers, and F. Bongers. 2006. Architecture of 54 moist-forest tree species: traits, trade-offs, and functional groups. Ecology 87:1289–1301.

Pretzsch, H. 2014. Canopy space filling and tree crown morphology in mixed-species stands compared with monocultures. Forest Ecology and Management 327:251–264.

Pretzsch, H., D. I. Forrester, and T. Rötzer. 2015. Representation of species mixing in forest growth models. A review and perspective. Ecological Modelling 313:276–292.

Purves, D. W., J. W. Lichstein, N. Strigul, and S. W. Pacala. 2008. Predicting and understanding forest dynamics using a simple tractable model. Proceedings of the National Academy of Science of the U.S.A 105:17018–17022.

R-Core-Team. 2018. R: A language and environment for statistical computing. R Foundation for Statistical Computing, Vienna, Austria.

Rambal, S., M. Lempereur, J. M. Limousin, N. K. Martin-StPaul, J. M. Ourcival, and J. Rodríguez-Calcerrada. 2014. How drought severity constrains gross primary production(GPP) and its partitioning among carbon pools in a *Quercus ilex* coppice? Biogeosciences 11:6855–6869.

Rameau, J.-C., D. Mansion, and G. Dumé. 1989. Flore forestière française. Guide écologique illustré. 1. Plaines et collines. Institut pour la Développement Forestier, Ministère de l’Agriculture et de la forêt, Paris.

Rameau, J.-C., D. Mansion, G. Dumé, and C. Gauberville. 2008. Flore forestière française. Guide écologique illustré. 3. Région Méditerranéenne. Institut pour la Développement Forestier, Paris.

Rasche, L., L. Fahse, A. Zingg, and H. Bugmann. 2011. Getting a virtual forester fit for the challenge of climatic change. Journal of Applied Ecology 48:1174–1186.

Rasche, L., L. Fahse, A. Zingg, and H. Bugmann. 2012. Enhancing gap model accuracy by modeling dynamic height growth and dynamic maximum tree height. Ecological Modelling 232:133–143.

Rees, M., R. Condit, M. Crawley, S. Pacala, and D. Tilman. 2001. Long-term studies of vegetation dynamics. Science 293:650–655.

Reich, P. B. 2014. The world-wide ‘fast–slow’ plant economics spectrum: a traits manifesto. Journal of Ecology 102: 275–301. doi: 10.1111/1365-2745.12211

Reich, P. B., and Walters, M. B. 1994. Photosynthesis-nitrogen relations in Amazonian tree species. Oecologia 97(1), 73–81. doi: 10.1007/BF00317910

del Río, M., H. Pretzsch, R. Ruíz-Peinado, E. Ampoorter, P. Annighöfer, I. Barbeito, K. Bielak, G. Brazaitis, L. Coll, L. Drössler, M. Fabrika, D. I. Forrester, M. Heym, V. Hurt, V. Kurylyak, M. Löf, F. Lombardi, E. Madrickiene, B. Matović, F. Mohren, R. Motta, J. den Ouden, M. Pach, Q. Ponette, G. Schütze, J. Skrzyszewski, V. Sramek, H. Sterba, D. Stojanović, M. Svoboda, T. M. Zlatanov, and A. Bravo-Oviedo. 2017. Species interactions increase the temporal stability of community productivity in *Pinus sylvestris-Fagus sylvatica* mixtures across Europe. Journal of Ecology 105:1032–1043.

Ruiz-Benito, P., G. Vacchiano, E. R. Lines, C. P. O. Reyer, S. Ratcliffe, X. Morin, F. Hartig, A. Mäkelä, R. Yousefpour, J. E. Chaves, A. Palacios-Orueta, M. Benito-Garzón, C. Morales-Molino, J. J. Camarero, A. S. Jump, J. Kattge, A. Lehtonen, A. Ibrom, H. J. F. Owen, and M. A. Zavala. 2020. Available and missing data to model impact of climate change on European forests. Ecological Modelling 416:108870.

San-Miguel-Ayanz, J., de Rigo, D., Caudullo G., Houston Durrant, T., Mauri, A., Tinner, W., Ballian, D., Beck, P., Birks, H. J. B., Eaton, E., Enescu, C. M., Pasta, S., Popescu, I., Ravazzi, C., Welk, E., Abad Viñas, R., Azevedo, J. C., Barbati, A., Barre, B. 2016. European Atlas of Forest Tree Species. Page (A. San-Miguel-Ayanz, J., de Rigo, D., Caudullo, G., Houston Durrant, T. Mauri, Ed.). European Commission, Luxembourg.

Schnabel, F., J. A. Schwarz, A. Dănescu, A. Fichtner, C. A. Nock, J. Bauhus, and C. Potvin. 2019. Drivers of productivity and its temporal stability in a tropical tree diversity experiment. Global Change Biology 25: 4257–4272

Schwinning, S., and J. Weiner. 1998. Mechanisms determining the degree of size asymmetry in competition among plants. Oecologia 113:447–455.

Seidl, R., D. Thom, M. Kautz, D. Martin-Benito, M. Peltoniemi, G. Vacchiano, J. Wild, D. Ascoli, M. Petr, J. Honkaniemi, M. J. Lexer, V. Trotsiuk, P. Mairota, M. Svoboda, M. Fabrika, T. A. Nagel, and C. P. O. Reyer. 2017. Forest disturbances under climate change. Nature Climate Change 7:395–402.

Shugart, H. H. 1984. A theory of forest dynamics: The ecological implications of forest succession models. Springer-Verlag, New York.

Simioni, G., G. Marie, and R. Huc. 2016. Influence of vegetation spatial structure on growth and water fluxes of a mixed forest: Results from the NOTG 3D model. Ecological Modelling 328:119–135.

Skovsgaard, J. P., and J. K. Vanclay. 2008. Forest site productivity: a review of the evolution of dendrometric concepts for even-aged stands. Forestry 81:13–31.

Strigul, N., D. Pristinski, D. Purves, J. Dushoff, and S. Pacala. 2008. Scaling from trees to forests: tractable macroscopic equations for forest dynamics. Ecological Monographs 78:523–545.

TEEB. 2010. The Economics of Ecosystems and Biodiversity Ecological and Economic Foundations. Page (P. Kumar, Ed.). Earthscan, London and Washington.

Toïgo, M., T. Perot, B. Courbaud, B. Castagneyrol, J.-C. Gégout, F. Longuetaud, H. Jactel, and P. Vallet. 2018. Difference in shade tolerance drives the mixture effect on oak productivity. Journal of Ecology 106:1073–1082.

Trouvé, R., J.-D. Bontemps, I. Seynave, C. Collet, and F. Lebourgeois. 2015. Stand density, tree social status and water stress influence allocation in height and diameter growth of *Quercus petraea* (Liebl.). Tree Physiology 35:1035–1046.

Ulrich, E. 1997. Organization of forest system monitoring in France-the RENECOFOR network. Page World Forestry Congress. Antalya, Turkey.

Vanoni, M., M. Cailleret, L. Hülsmann, H. Bugmann, and C. Bigler. 2019. How do tree mortality models from combined tree-ring and inventory data affect projections of forest succession? Forest Ecology and Management 433:606–617.

Verdone, M., and A. Seidl. 2017. Time, space, place, and the Bonn Challenge global forest restoration target. Restoration Ecology 25:903–911.

Verheyen, K., M. Vanhellemont, H. Auge, L. Baeten, C. Baraloto, N. Barsoum, S. Bilodeau- Gauthier, H. Bruelheide, B. Castagneyrol, D. Godbold, J. Haase, A. Hector, H. Jactel, J. Koricheva, M. Loreau, S. Mereu, C. Messier, B. Muys, P. Nolet, A. Paquette, J. Parker, M. Perring, Q. Ponette, C. Potvin, P. Reich, A. Smith, M. Weih, and M. Scherer-Lorenzen. 2016. Contributions of a global network of tree diversity experiments to sustainable forest plantations. Ambio 45:29–41.

Verkerk, P. J., J. B. Fitzgerald, P. Datta, M. Dees, G. M. Hengeveld, M. Lindner, and S. Zudin. 2019. Spatial distribution of the potential forest biomass availability in Europe. Forest Ecosystems 6:5.

Vidal, J.-P., E. Martin, L. Franchistéguy, M. Baillon, and J.-M. Soubeyroux. 2010. A 50-year high-resolution atmospheric reanalysis over France with the Safran system. International Journal of Climatology 30:1627–1644.

Violle, C., M. L. Navas, D. Vile, E. Kazakou, C. Fortunel, I. Hummel, and E. Garnier. 2007. Let the concept of trait be functional! Oikos 116:882–892.

Wehrli, A., P. J. Weisberg, W. Schönenberger, P. Brang, and H. Bugmann. 2007. Improving the establishment submodel of a forest patch model to assess the long-term protective effect of mountain forests. European Journal of Forest Research 126:131–145.

Williams, L. J., A. Paquette, J. Cavender-Bares, C. Messier, and P. B. Reich. 2017. Spatial complementarity in tree crowns explains overyielding in species mixtures. Nature Ecology & Evolution 1:0063.

