## Appendix A for "Beyond forest succession: a gap model to study ecosystem functioning and tree community composition under climate change"

### Appendix A: Description of the model ForCEEPS

#### Model overview

ForCEEPS is a forest gap model (or forest dynamics model) following the scheme of JABOWA (Botkin et al. 1972) and ForCLIM (Bugmann 1996, Didion et al. 2009), with notable exceptions. It simulates forest dynamics and succession by considering the effects of abiotic (climate and soil properties) and biotic constraints (tree-tree competition for light) on tree establishment, growth, and survival in small parcels of land ("patches") (Fig. A).

Simulated patches are independent from each other, and properties at the forest level are obtained by aggregating the properties over all patches. Within each patch, environmental conditions are assumed to be horizontally homogeneous, and patches are usually do not exceed 1000 $m^2$ to ensure this assumption. The spatial location of trees is therefore implicit. This allows for several simplifications in the representation of tree-tree interactions, but imposes that the patch size cannot be larger than 1000 $m^2$, which is considered as the maximum area influenced by a tree (Shugart 1984). Gap models are often cohort-based, assuming that all trees of the same species and age behave similarly, for the sake of simulation efficiency, but ForCEEPS is completely individual-based, which notably allows to take inter-individual variability into account. ForCEEPS also allows imposing a feedback between the actual forest composition and the identity of the colonizing seedlings each year. The main development of ForCEEPS in comparison with the classic scheme of gap model is the implementation of a new module for light competition where the individual crown lengths are explicitly represented in the vertical canopy space.

The main processes of the model are described in the next sections, according to Figure B illustrating how the input variables and species parameters operate in ForCEEPS.

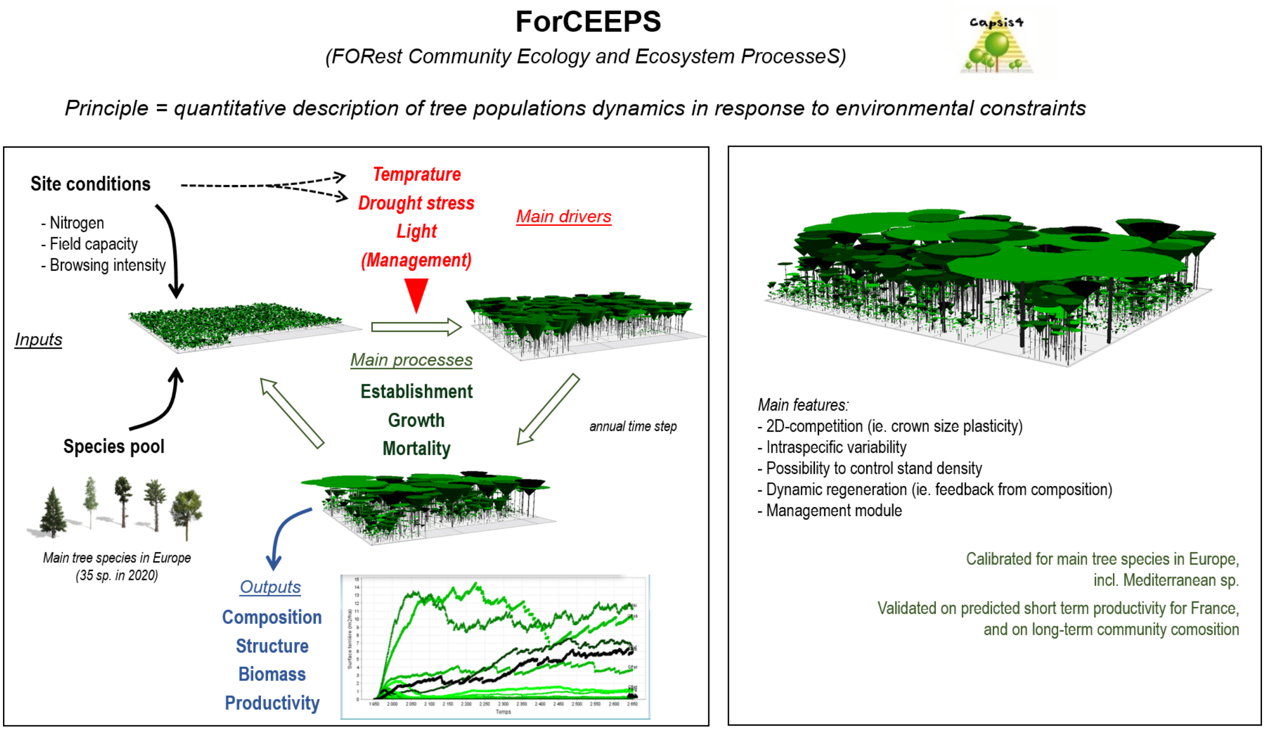


Figure A: Structure and features of ForCEEPS.

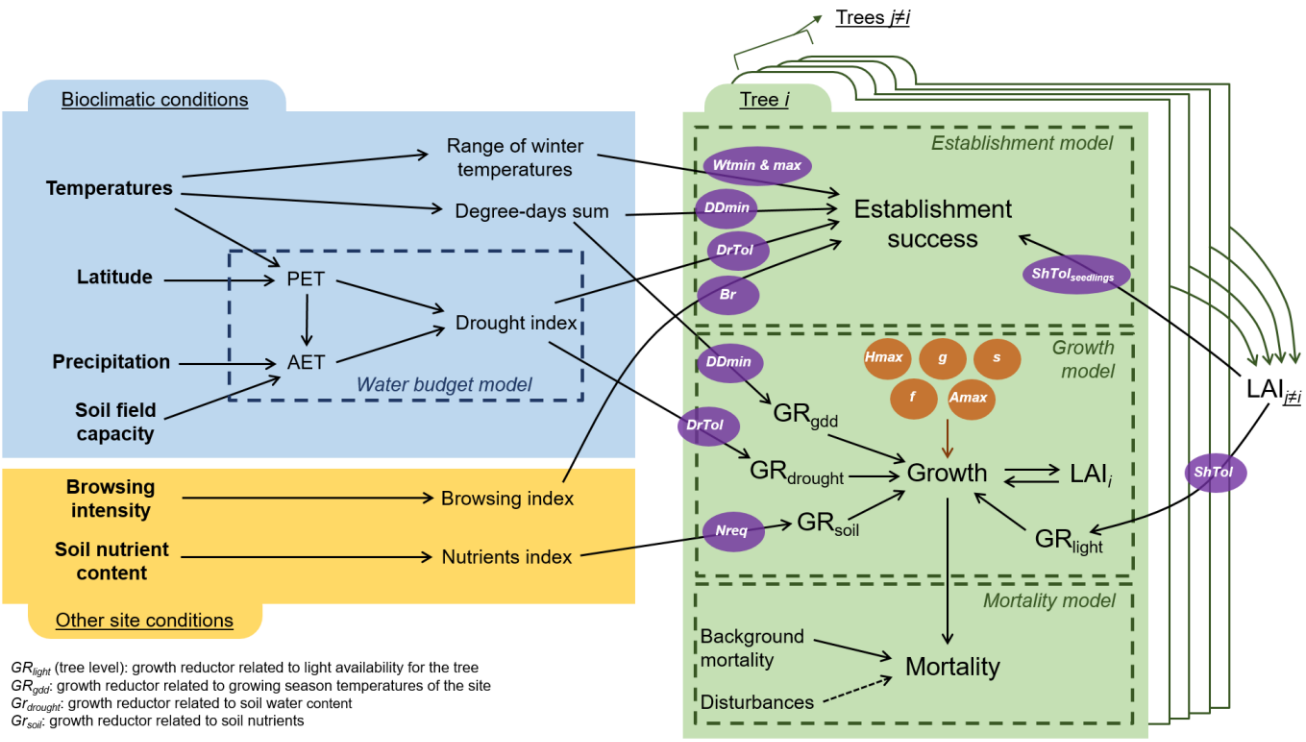


Figure B: Detailed processes embedded in the model. Species parameters (orange: intrisic parameters; purple: response-to-drivers parameters) are described in Table A.

**Detailed description of the model**

The core-processes of the model are tree establishment, growth, and mortality are simulated at a yearly time step, and are all possibly constrained by environmental drivers (Fig. A and B). We first present how the environmental drivers are calculated or assessed, and second we detail how the core-processes are modelled in ForCEEPS.

*Environmental drivers*

Bioclimatic part

ForCEEPS uses time-series of monthly climate data (monthly mean temperature and precipitation sum). Climatic indices are calculated from these climate data: annual (for evergreen species) or seasonal (for deciduous species) growing degree-days sum (*GDD*), minimum and maximum winter temperatures, and a drought index (*DrI*) (Fig. B).

- Growing degree-days sums (*GDD*) are calculated as follows:

$$GDD = \sum_{m=i}^{j} \max(T_m - T_0) \times N_{days} \times k_{corr} \qquad \text{(Eq. 1)}$$

with $T_m$ is the mean monthly temperature, $T_0$ is the base tempearure ($T_0$ = 5.5°C), $N_{days}$ the average number of days per month ($N_{days}$ = 30.5) and $k_{corr}$ being a correction coefficient to avoid bias related to the calculations of degree-days from monthly means (Bugmann 1994):

$$k_{corr} = \begin{cases} 8.52 \cdot 10^{0.165 \cdot T_m} \; if \; T_m \leq 5.5 \\ 187.2 \cdot 10^{0.0908 \cdot T_m} \quad if \; 5.5 < T_m \leq 15.5 \\ -31.8 + 2.377 \cdot T_m \; if \; T_m > 15.5 \end{cases} \qquad \text{(Eq. 2)}$$

For evergreen and deciduous species, *i*=1 and *i*=4, and *j*=12 and *j*=10, respectively.

- Mean winter temperature $T_W$ is the average of the mean temperatures of the three month: December (year *n*-1), January (year *n*) and February (year *n*).

- Drought stress index (DrI) is assessed through the calculation of soil water content (*SWC*) from a water budget based on monthly calculations of potential (*PET*) and actual evapotranspiration (*AET*), and depending on the site-specific soil field capacity (Fig. B), as in the ForClim model (Bugmann & Solomon 2000).

To calculate evapotranspiration and water balance, the more accurate methods require many weather variables with a high temporal resolution. As in ForClim (Bugmann & Solomon 2000), ForCEEPS relies on the model by Thornthwaite & Mather (1957). This is an entirely empirical approach (ie. a «bucket model »), but it is especially useful because it only requires monthly mean temperatures and monthly precipitation sums (and the latitude to derive day length), and has been shown to provide reasonable estimates of PET and AET (Bugmann 1994). We should notice that, at least in this version of the model, this approach assumes that *AET* is independent of the vegetation cover and composition, and is thus based on an average leaf area index. However, such an assumption is justified by the fact that canopy openings caused by the death of single trees are relatively small and their effect on evapotranspiration rates is moderate.

The drought index *DrI* is thus calculated from monthly drought indices $DrI_m$ with

$$DrI_m = \begin{cases} 1 - \frac{E_m}{D_m}, & D_m \neq 0 \\ 0, & D_m = 0 \end{cases} \qquad \text{(Eq. 3)}$$

where $E_m$ is the amount of water transpired by the trees and $D_m$ is their evaporative demand for soil water during month *m* (Bugmann & Cramer 1998; Bugmann and Solomon 2000). Then *DrI* is simply the average of $DrI_m$ over the relevant time period, by only considering months when transpiration and demand are significant (ie. with $T_m \geq T_0$), with thus

$$DrI = mean(DrI_m) \mid i \leq m \leq j, T_m \geq T_0 \qquad \text{(Eq. 4)}$$

where $T_0$ = 5.5°C, and *i*=1 and *i*=4, and *j*=12 and *j*=10, for evergreen and deciduous species, respectively.

<u>Other drivers</u>

- Another abiotic factor taken into account in ForCEEPS is soil nutrients content ($N_{soil}$), which is constant at the site level, while climate conditions do vary among years. $N_{soil}$ is in kg.ha$^{-1}$, although it should rather be seen as a quantitative proxy for the edaphic quality of the site's soil.

- A browsing intensity *Br* can be defined for each site as a continuous value between 0 (no pressure) and 1 (high pressure), constant at the site level across years.

*Seedling establishment*

By default, the model assumes that there is a constant seed rain in the patches and thus no dispersal limitation. Therefore, each species present in the species list at the beginning of the simulation sends seeds each year. Alternatively it is possible in ForCEEPS to activate a feedback between the actual forest composition at year *n* and species composition of the new seedlings at year *n*+1. In this case, the composition of the seed rain is determined by the relative biomass of all species actually present in the patches (ie. with adult trees).

Furthermore, in ForCEEPS, seedling establishment can be constrained by defining a maximum number of seedlings potentially colonizing the patches and/or by imposing a feedback of actual forest composition on the composition of colonizing seedlings.

Seedlings are established with a diameter at breast height of 1 cm. Establishment success is simulated as a function of species-specific responses to *DrI, GDD,* $T_W$ *, Br* (Didion et al. 2011), and light availability at the forest floor $L_{av}$ (see Eq. 28). More precisely, the probability $P_{est_S}$ of establishment success for each seedling of the species *S* is calculated as follows:

$$P_{est_S} = P_{T_W} \times P_{GDD} \times P_{Dr} \times P_{Br} \times P_{LA} \times c_{est} \qquad \text{(Eq. 5)}$$

$P_{T_W}$ is the probability that the seedling can establish according to winter temperature $T_W$, and $Wtmin_S$ and $Wtmax_S$, respectively the monthly minimum and maximum winter temperature (°C) tolerated for regeneration of species *S* (species parameters are described in Table A and values for main species in Table B), with

$$P_{T_W} = \begin{cases} 1, & Wtmin_s \leq T_W \leq Wtmax_s \\ 0, & otherwise \end{cases} \quad \text{(Eq. 6)}$$

$P_{GDD}$ is the probability that the seedling can establish according to GDD of the year (Eq. 1) and the minimal required annual or seasonal degree-days sum $DDmin_S$, the establishment assumed to be impossible when the annual sum of growing degree-days does not conform to the degree-day requirements of the tree species.

$$P_{GDD} = \begin{cases} 1, & GDD \geq DDmin_s \\ 0, & GDD < DDmin_s \end{cases} \quad \text{(Eq. 7)}$$

$P_{DrI}$ is the probability that the seedling can establish according to the drought stress of the year (Eq. 3 and 4) and the species-specific drought tolerance $DrTol_S$, with:

$$P_{Dr} = \begin{cases} 1, & DrI \leq DrTol_s \\ 0, & DrI > DrTol_s \end{cases} \quad \text{(Eq. 8)}$$

$P_{Br}$ is the probability that the seedling can establish according to the browsing intensity of the site *Br* and to $Br_S$, the browsing susceptibility of seedlings of species *S*, that both determine the actual browsing susceptibility of seedlings of species S in the site $B_{actual}$, with:

$$if\ Br_S = 1, \quad B_{actual} = Br^4$$

$$if\ Br_S = 2, \quad B_{actual} = Br^2$$

$$if\ Br_S = 3, \quad B_{actual} = Br$$

$$if\ Br_S = 4, \quad B_{actual} = Br^{0.5}$$

$$if\ Br_S = 5, \quad B_{actual} = Br^{0.25}$$

Then $P_{Br}$ is calculated as:

$$P_{Br} = \begin{cases} 1, & B_{actual} \leq k_{Br} \\ 0, & B_{actual} > k_{Br} \end{cases} \quad \text{(Eq. 9)}$$

$k_{Br}$ being a random number between 0 and 1.

$P_{LA}$ is the probability that the seedling can establish according to the light availability at the forest floor $L_{av}$ and the species-specific shade tolerance of seedlings $ShTol\_seedling_S$, with:

$$P_{LA} = \begin{cases} 1, & L_{av} \geq ShTol\_seedling_s \\ 0, & L_{av} < ShTol\_seedling_s \end{cases} \qquad \text{(Eq. 10)}$$

with $L_{av} = e^{-k \times LA}$, LA being the summed leaf area over all trees in the patch.

The coefficient $c_{est}$ is a random factor comprised between 0 and 1 to introduced stochasticity in the seedling regeneration.

Finally, The establishment of the seedlings of a given tree species $S$ is possible if

$$P_{est_S} \geq k_{est} \qquad \text{(Eq. 11)}$$

where $k_{est}$ is a random number comprised between 0 and 1.

If seedling establishment is possible for species $S$, the number of seedlings that actually established is calculated using a random number with uniform distribution in the range [1 … $k_{EstNr}.k_{PatchSize}.ShTol_S$], where $k_{EstNr}$ is the maximum seedling establishment rate ($k_{EstNr}$ = 0.006 seedlings per $m^{-2}.yr^{-1}$, Bugmann 1996), $k_{PatchSize}$ is the size of the forest patch in $m^2$ (see Table D), and $ShTol_S$ is the shade tolerance of the species (integer between 1 – high tolerance - and 9 – low tolerance) to take into account that pioneer species produce more seedlings than late-successional species (Didion et al. 2009).

*Tree growth*

<u>Potential growth</u>

Annual tree growth is modelled through stem diameter increment at breast height (*ΔD*). Following the classical scheme of gap models, *ΔD* is calculated in two steps.

First, the potential (i.e. maximum) diameter increment ($\Delta D_{\text{opt}}$) of each tree is predicted each year using the following empirical equation (Moore 1989):

$$\Delta \mathrm{D}_{\mathrm{opt}} = g_S \frac{\mathrm{D}\left(1 - \frac{\mathrm{H}}{H_{max_S}}\right)}{2.H_{max_S} - b_S \times e^{(c_S.\mathrm{D}) \times (c_S.\mathrm{D}+2)}} \qquad \text{(Eq. 12)}$$

where $D$ is tree diameter at breast height, $H$ is tree height, $g_s$ is the maximum growth rate of species $s$, $H_{max_s}$ is the maximum height reachable by the species $s$, and $b_s$ and $c_s$ are species-specific parameters (with $b_s = H_{max_s} - 137$; and $c_s = s_s / b_s$); $s_s$ is a species-specific allometric parameter relating tree height $H$ and tree diameter $D$ as follows (Bugmann 1996):

$$\mathrm{H} = a + \left(H_{max_s} - a\right) \times \left(1 - e^{\left(\frac{-s_S.\mathrm{D}}{H_{max_S} - a}\right)}\right) \quad \text{(Eq. 13)}$$

with a = 1.37 m (i.e. breast height). Therefore, simulating the potential diameter increment of a tree in FORCEEPS requires to determine the values of the species-specific parameters $g_s$, $s_s$ and $H_{max_s}$ (Tables A and B).

Realized growth

Second, realized tree diameter increment $\Delta D$ is calculated by modifying $\Delta D_{\mathrm{opt}}$ according to abiotic or biotic growth reduction factors (all factors are bounded between 0 and 1) as follows (Bugmann 1996, Didion et al. 2009):

$$\Delta\mathrm{D} = \Delta\mathrm{D}_{\mathrm{opt}} \times \sqrt[3]{GR_{light} \times GR_{gdd} \times GR_{drought} \times GR_{soil}} \quad \text{(Eq. 14)}$$

where $GR_{light}$ is the growth reduction factor related to light availability for the tree, $GR_{gdd}$ is the growth reduction factor related to growing season temperatures of the site (*GDD*), $GR_{drought}$ is the growth reduction factor related to the drought index (*DrI*), and $GR_{soil}$ is the growth reduction factor related to soil nutrients content ($N_{soil}$). The effects of each of these growth reduction factors on realized tree growth depend on species-specific parameters: $GR_{light}$ depends on species shade tolerance $ShT_s$; $GR_{gdd}$ depends on species minimum sum of growing degree-days $GDD_s$; $GR_{drought}$ depends on species drought tolerance $DrT_s$; and $GR_{soil}$ depends on species requirements for soil nutrients $NReq_s$ (see Table C). All growth reduction factors vary among site conditions and species, and $GR_{light}$ varies also among trees, because it is influenced by the sizes of the neighbouring trees in the patch (see next section).

More precisely, the growth redactors are calculated as follows:

$$\mathrm{GR}_{gdd} = max\left(0, 1 - e^{(DDmin_S - GDD)\times q_{corr}}\right) \qquad \text{(Eq. 15)}$$

With $q_{corr}$ being the curve slope parameter in degree-day growth function ($q_{corr}$ = 1/750 °C$^{-1}$.day$^{-1}$) (Bugmann 1996).

$$\mathrm{GR}_{drought} = \sqrt{max\left(0, 1 - \frac{DrI}{DrTol_S}\right)} \qquad \text{(Eq. 16)}$$

$$\mathrm{GR}_{soil} = max\left(0, 1 - e^{[N1_S \times (N_{soil} - N2_S)]}\right) \qquad \text{(Eq. 17)}$$

With $N1_S$ and $N2_S$ being species-specific parameters derived from $Nreq_S$ as described in Table C.

Growth reduction due to competition for light.

As the trees are not spatialized inside the patches, the interaction for light among trees in ForCEEPS depends on the vertical stratification of tree crowns alone. Following the framework of ForClim 2.9.6 (Didion et al. 2009), the amount of light available for a tree (with *H* being its total height) is reduced by the leaf area of all trees found in the same patch whose height is greater than *H* or equal to *H*. Thus, all the foliage of trees taller than the target tree contribute to the shading. Aiming to reach a more integrative description of the competition for light, ForCEEPS embeds a more realistic competition module, by representing individual crown lengths (Fig. C).

In ForCEEPS, $GR_{light}$ has two components:

$$GR_{light} = GR_{cs} \times GR_{sh} \qquad \text{(Eq. 18)}$$

with $GR_{cs}$ representing the feedback of crown size on tree growth, i.e., tree leaf area is positively linked to tree growth rate (Mitscherlich and von Gadow 1968). $GR_{sh}$ is the reduction factor related to shading by competing trees. Crowns of individual trees are thus explicitly represented in the vertical space of the canopy (Fig. C). To do so, we introduce crown length *cl*, calculated as follows for each tree *i*:

$$cl_i = cs_i \times H_i \qquad \text{(Eq. 19)}$$

with $H$ being tree height and $cs$ being the ratio of the height with a green crown, which is related to light exposition of the tree (Didion et al. 2009). For each tree $i$, $cs$ varies between two extreme species-specific values that represent the case where the tree is fully shaded ($cs = cs_{min_s}$) or in full light ($cs = cs_{max_s}$), with:

$$cs_i = cs_{max_s} - \left(cs_{max_s} - cs_{min_s}\right) \times k_{LAI_i} \qquad \text{(Eq. 20)}$$

where the extreme values $cs_{max_s}$ and $cs_{min_s}$ have been derived from the relationship between foliage fresh weight and DBH described in Wehrli et al. (2007) and depends on the foliage type parameter $f_S$ (see Table C), and $k_{LAIi}$ is the correction factor - ranging from 0 (no shading) to 1 (full shading) - calculated by Didion et al. (2009) as follows:

$$k_{LAI_i} = min\left[\left(\frac{LAI_{H_i}}{LAI_{max}}\right)^2, 1\right] \qquad \text{(Eq. 21)}$$

with $LAI_H$ being the cumulative leaf area index between the top of the canopy and the top of the target tree (i.e. between the top of the canopy and the height $H$), ie. the cumulative leaf area (in $m^2$) divided by the area of the patch (in $m^2$) and $LAI_{max}$ being the maximum value of double-sided lead area in a patch resulting from light compensation point in European forests [i.e. $LAI_{max}$ = 11.98 (Bugmann 1994, Didion et al. 2009)].

Consequently, to allow the competition among trees whose crowns overlap, the vertical space of the patch $p$ at simulation step $t=t_1$ is discretized in $n(p,t_1)$ layers of a given width $w$, whose value is bounded between 0 (ground level) to $H_{max}(p, t_1)$ (height of the tallest tree in the patch $p$ at $t=t_1$), with $w$ = 1 m. Tree leaf area is assumed to decrease linearly from the top to the base of the crown, i.e. from the highest to the lowest layer in which the crown of the tree is found (Fig. C) (Eermak 1998, Van Pelt et al. 2016). We are aware that tree crown shape and vertical leaf area distribution vary among tree species and are also affected by the size and identity of neighbouring trees (Poorter et al. 2006, Williams et al. 2017, Niklaus et al. 2017). Our assumption should thus be seen as a first parsimonious step that can be refined using species- and context-specific architectural data.

$GR_{cs}$ (Eq. 4) is calculated as follows for each tree $i$ of species $s$:

$$GR_{cs_i} = min\left(z \times \frac{cs_i}{cs_{max_s}} \times \frac{LCP_s}{LCP_{mean}}, 1\right) \quad \text{(Eq. 22)}$$

with $z$=4/3, $LCP_s$ is the light compensation point of the species $s$ that directly depends on the shade tolerance parameter $ShTol_s$ as follows

$$LCP_S = LCP_{max} - (LCP_{max} - LCP_{min}) \times \frac{(ShTol_S - ShTol_{max})}{(ShTol_{min} - ShTol_{max})} \quad \text{(Eq. 23)}$$

with $LCP_{min}$ and $LCP_{max}$ being the light compensation point of the species with the minimum and maximum value for the shade tolerance parameter $ShTol_s$ respectively, ie. $ShTol_{min}$ ($ShTol_s$=9) and $ShTol_{max}$ ($ShTol_s$=1) (see Table D). $LCP_{mean}$ is the mean light compensation point among the most shade tolerant and the most shade intolerant species across European tree species (Bugmann 1994), ie. $LCP_{mean} = \frac{LCP_{max} - LCP_{min}}{2}$.

For a given patch, foliage area index is calculated for each layer, by summing the foliage area index of all crown parts found in this layer. Thus, for the layer $x$ of width w, the total foliage area index $SumLAI_x$ is:

$$SumLAI_x = \frac{1}{PA} \times \sum_{i|H_i \geq x-w} LA_{i,x} \quad \text{(Eq. 24)}$$

with $PA$ being the patch area, $H_i$ the height of tree $i$, and $LA_{i,x}$ the double-sided leaf area of the crown part of the tree $i$ belonging to layer $x$, calculated as follows:

$$LA_{i,x} = r_{i,x}.LA \quad \text{(Eq. 25)}$$

where $r_{i,x}$ corresponds to the fraction of total foliage found in layer $x$ (assuming a linear decrease in the foliage area, as shown in Fig. C), and the consistency between the layer and whole tree levels being ensured by the fact that:

$$\sum_{x_{H_0} \leq x \leq x_H} r_{i,x} = 1 \quad \text{(Eq. 26)}$$

with $x_{H_0}$ and $x_H$ being the layer including the base and the top of the crown of tree $i$, respectively, and $LA$ being the total double-sided leaf area of a tree, is calculated from tree diameter $D$ as follows (Didion et al. 2009):

$$LA = f'_S . cs . D^{a_s} \qquad \text{(Eq. 27)}$$

with $f'_s$ and $a_s$ being allometric parameters (see Table C), which depend on the foliage type of the species (parameter $f_S$, Tables A and B) (Wehrli et al. 2007).

Then, an intermediate variable ($\text{GR}_{sh_{i,x}}$) is calculated using light availability $L_{av}$ in the layer according to Beer's extinction law (Botkin et al. 1972, Shugart 1984) to quantify shading intensity in each crown layer of each tree. For a tree *i* found in layer *x,* the light availability $L_{av_{i,x}}$ is calculated as follows:

$$L_{av_{i,x}} = r_{i,x} . e^{-k.SumLAI_x} \qquad \text{(Eq. 28)}$$

with $r_{i,x}$ being the coefficient representing the decrease in foliage area from the top to base of the crown (see Eq. 26) and $k$ is the light extinction coefficient (with k=0.25).

Then $\text{GR}_{sh_{i,x}}$ is obtained by considering the light response function of a species (Bugmann 1996), as follows:

$$\text{GR}_{sh_{i,x}} = max\left(0,\ L_{max} + (L_{min} - L_{max}) \times \frac{ShTol_S - ShTol_{max}}{ShTol_{min} - ShTol_{max}}\right) \qquad \text{(Eq. 29)}$$

with $L_{max} = 1 - c_{max}{}^{-(L_{av_{i,x}} - 0.05)}$ and $L_{min} = 2.24 \times \left(1 - c_{min}{}^{-(L_{av_{i,x}} - 0.08)}\right)$, $c_{max}$ and $c_{min}$ being constant ($c_{max}$ = -4.64 and $c_{min}$ =-1.136), and $L_{av_{i,x}}$being the light availability for tree *i* found in layer *x* (see Eq. 26).

Finally, the reduction factor related to shading by competing trees ($\text{GR}_{sh_i}$) is calculated for a tree *i* by summing the $\text{GR}_{LA_{i,x}}$ of all crown parts of the tree:

$$\text{GR}_{sh_i} = \textstyle\sum_x \text{GR}_{sh_{i,x}} \qquad \text{(Eq. 30)}$$

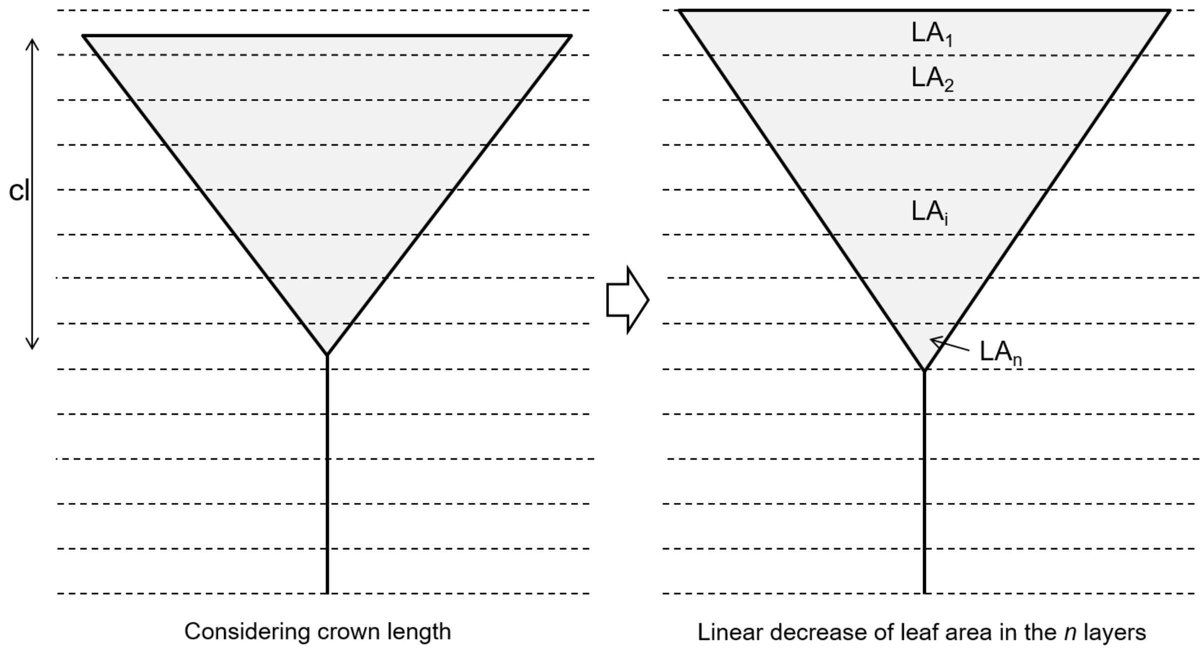


Figure C. Representation of the foliage distribution at the tree level in ForCEEPS. In ForCEEPS, the foliage is distributed in the vertical space along the crown length of the tree ("*cl*" on lower panel), and tree leaf area was assumed to decrease linearly from the top to the base of the crown, through the discretization of the vertical space in layers as represented in the lower panel. *H* is the total height of the tree.

*Tree mortality*

The probability $P_{mort,i}$ that tree *i* dies on year *y* depends on two components: (i) a "background" mortality $P_0$, and (ii) a growth-related mortality $P_g$:

$$P_{mort,i} = \max\left(P_0, P_g\right) \qquad \text{(Eq. 30)}$$

$P_0$ is calculated as follows:

$$P_0 = \begin{cases} 0, & m_0 < p_{dist} \\ 1, & m_0 \geq p_{dist} \end{cases} \qquad \text{(Eq.31)}$$

With $m_0$ being a random number sorted between 0 and 1, and $p_{dist}$ being a constant representing the background or disturbance probability (see Table D). The background mortality $P_0$ is thus purely stochastic and depends on species' maximum longevity, and is implemented to consider mortality events induced by 'random' small-scale disturbances (e.g., attack of pathogen in an endemic phase), but large-scale disturbances (e.g., windthrows, wildfires) can be taken into account by increasing the background mortality rate (ie. $p_{dist}$, Table D for an example of parameter value)

The growth-related mortality is a proxy for stress conditions and considers that tree mortality probability increases with the decrease in absolute or relative tree growth rates (i.e. tree vigor) induced by abiotic factors or by competition. $P_g$ is calculated as follows:

$$P_g = P_{age} + \left(1 - P_{age}\right) \times P_{stress,i} \qquad \text{(Eq. 32)}$$

$P_{age}$ is the age-related mortality probability, depending on the maximum age of the species $Amax_S$:

$$P_{age} = \frac{k_{death}}{Amax_S} \qquad \text{(Eq. 33)}$$

with $k_{death}$ being a constant, corresponding to a death probability coefficient (Table D).

$P_{stress}$ is the stress-induced mortality probability, which depends on tree growth, calculated as follows:

$$P_{stress,i} = \begin{cases} p_{slowgr}, & SlowGr_i \geq k_{SlowGrTime} \\ 0, & elsewhere \end{cases} \qquad \text{(Eq. 34)}$$

With $p_{slowgr}$ being a constant representing the mortality probability for stressed trees (Table D), $SlowGr_i$ is an index corresponding to the number of ongoing consecutive years for which the growth of tree *i* is lower than the threshold $t_{slowgr}$ (constant, Table D), and $k_{SlowGrTime}$ is a constant (in years, Table D) representing the number of years of slow growth after which the tree is considered as stressed (Bugmann 1994, Bugmann 1996).

Finally $P_{mort,i}$ is compared to a random number $m_{tot}$ comprised between 0 and 1. If $m_{tot} < P_{mort,i}$ then tree *i* dies.

*Tree biomass*

In ForCEEPS, the total aboveground biomass of a tree is devided in its stem wood biomass and foliage weight, that are both estimated by using allometric equations (Bugmann 1994; Werhli et al. 2007; Didion et al. 2009) and are summed to calculate the total aboveground biomass.

Foliage weight $F_w$ depends on stem diameter at breast height $D$ and crown size $cs$, as follows:

$$F_w = f''_s \times cs \times D^{a_s} \qquad \text{(Eq. 35)}$$

with $f''_s$ and $a_s$ being allometric parameters (see Table C), which depend on the foliage type of the species (parameter $f_S$, see Table 1) (Wehrli et al. 2007). Note that $F_w = \frac{f''_S}{f'_S} \times LA$ (see Eq. 26).

Stem wood biomass $B$ only depends on stem diameter at breast height $D$ as follows (Bugmann 1994):

$$B = 0.12 \times D^{2.4} \qquad \text{(Eq. 36)}$$

*References*

Botkin, D. B., J. F. Janak, and J. R. Wallis. 1972. Some ecological consequences of a computer model of forest growth. Journal of Ecology 60:849–872.

Bugmann, H. 2001. A review of forest gap models. Climatic Change 51:259–305.

Bugmann, H. 1996. A Simplified Forest Model to Study Species Composition Along Climate Gradients. Ecology 77:2055–2074.

Bugmann, H. 1994. On the Ecology of Mountainous Forests in a Changing Climate: A Simulation Study. PhD Thesis 10638, ETHZ, Zürich.

Bugmann, H., and W. Cramer. 1998. Improving the behaviour of forest gam models along drought gradients. Forest Ecology and Management **103**: 247-263.

Bugmann, H., and A. M. Solomon. 2000. Explaining forest composition and biomass across multiple biogeographical regions. Ecological Applications 10:95–114.

Didion, M., A. D. Kupferschmid, A. Wolf, and H. Bugmann. 2011. Ungulate herbivory modifies the effects of climate change on mountain forests. Climatic Change 109:647–669.

Didion, M., A. D. Kupferschmid, A. Zingg, L. Fahse, and H. Bugmann. 2009. Gaining local accuracy while not losing generality — extending the range of gap model applications. Canadian Journal of Forest Research 39:1092–1107.

Eermak, J. 1998. Leaf distribution in large trees and stands of the floodplain forest in southern Moravia. Tree Physiology 18:727–737.

Mitscherlich, G., and K. von Gadow. 1968. Über den zuwachsverlust bei der ästung von nabelbäumen. Allgemaine Forst- und Jagdzeitung.

Moore, A. D. 1989. On the maximum growth equation used in forest gap simulation models. Ecological Modelling 45:63–67.

Niklaus, P. A., M. Baruffol, J.-S. He, K. Ma, and B. Schmid. 2017. Can niche plasticity promote biodiversity-productivity relationships through increased complementarity? Ecology 98:1104–1116.

Van Pelt, R., S. C. Sillett, W. A. Kruse, J. A. Freund, and R. D. Kramer. 2016. Emergent crowns and light-use complementarity lead to global maximum biomass and leaf area in Sequoia sempervirens forests. Forest Ecology and Management 375:279–308.

Poorter, L., L. Bongers, and F. Bongers. 2006. Architecture of 54 moist-forest tree species : traits, trade-offs, and functional groups. Ecology 87:1289–1301.

Shugart, H. H. 1984. A theory of forest dynamics: The ecological implications of forest succession models. Springer-Verlag, New York.

Thornthwaite, C.W.; Mather, J.R. 1957. Instructions and tables for computing potential evapotranspiration and the water balance. Publication in Climatology 10: 185-311.

Wehrli, A., P. J. Weisberg, W. Schönenberger, P. Brang, and H. Bugmann. 2007. Improving the establishment submodel of a forest patch model to assess the long-term protective effect of mountain forests. European Journal of Forest Research 126:131–145.

Williams, L. J., A. Paquette, J. Cavender-Bares, C. Messier, and P. B. Reich. 2017. Spatial complementarity in tree crowns explains overyielding in species mixtures. Nature Ecology & Evolution 1:0063.

*Tables*

Table A

Description of species parameters in ForCEEPS.

| Parameter | Details | Unit |
|---|---|---|
| $f_s$ | Foliage type | Unitless<br>E - evergreen - or D - deciduous and a number between 1 and 5 |
| $Hmax_s$ | Maximum height | m |
| $s_s$ | Allometry | Unitless |
| $g_s$ | Optimal growth | Unitless |
| $Amax_s$ | Maximum age | years |
| $DDmin_s$ | Minimal required annual or seasonal degree-days sum | °C |
| $DrTol_s$ | Drought tolerance index, to be compared to the evapotranspiration deficit based on a bucket model of soil moisture | Continuous index with values between<br>0 (sensitive) to 1 (tolerant) |
| $Nreq_s$ | Soil nitrogen requirement | Integer Index with values between<br>1 (weak requirements) to 5 (strong req.) |
| $ShTol_s$ | Shade tolerance | Integer index with values between<br>1 (shade tolerant) to 9 (shade intolerant) |
| $ShTol\_seedling_s$ | Shade tolerance of seedlings, to be compared to the relative amount of light reaching the ground | Continuous index with values between<br>0 (tolerant) to 1 (sensitive) |
| $Wtmin_s$ | Monthly minimum winter temperature tolerated for regeneration (°C) | °C |
| $Wtmax_s$ | Monthly maximum winter temperature tolerated for regeneration | °C |
| $Br_s$ | Browsing susceptibility of seedlings | Integer index with values between<br>1 (less susceptible) to 5 (more susceptible) |

Table B
ForCEEPS parameters for the species calibrated in this study. Parameter description, and literature used to calibrate all or part of the species for specific parameters can be found in Table 1.

| | Species | fS | HmaxS | sS | gS | AmaxS | DDminS | DrTolS | NreqS | ShTolS | ShTol_seedlingS | WtminS | WtmaxS | BrS |
|---|---|---|---|---|---|---|---|---|---|---|---|---|---|---|
| main species | *Abies alba* | E5 | 50 | 75 | 350 | 366 | 841 | 0.23 | 3 | 1 | 0.05 | -6 | 5 | 5 |
| | *Picea abies* | E5 | 48 | 83 | 355 | 300 | 421 | 0.11 | 2 | 5 | 0.1 | -20 | 3 | 2 |
| | *Pinus halepensis* | E4 | 22 | 69 | 399 | 200 | 1261 | 0.48 | 1 | 7 | 0.25 | 2 | 13 | 3 |
| | *Pinus pinaster* | E4 | 35 | 69 | 350 | 300 | 1121 | 0.4 | 1 | 7 | 0.25 | -1 | 12 | 3 |
| | *Pinus sylvestris* | E4 | 35 | 58 | 150 | 200 | 631 | 0.37 | 1 | 9 | 0.3 | -20 | 5 | 3 |
| | *Fagus sylvatica* | D3 | 50 | 76 | 260 | 400 | 841 | 0.25 | 2 | 1 | 0.05 | -6 | 9 | 3 |
| | *Quercus ilex* | E4 | 23 | 48 | 79 | 500 | 1121 | 0.45 | 1 | 5 | 0.1 | 0 | 14 | 1 |
| | *Quercus petraea* | D3 | 45 | 76 | 246 | 1000 | 981 | 0.33 | 2 | 7 | 0.2 | -7 | 10 | 4 |
| | *Quercus robur* | D3 | 45 | 66 | 249 | 1000 | 631 | 0.3 | 2 | 9 | 0.3 | -10 | 10 | 4 |
| other species | *Larix decidua* | D2 | 35 | 72 | 170 | 500 | 600 | 0.25 | 1 | 9 | 0.4 | -8 | -5 | 3 |
| | *Pinus cembra* | E5 | 25 | 40 | 115 | 1000 | 600 | 0.3 | 1 | 5 | 0.2 | -9 | -2 | 4 |
| | *Acer campestre* | D2 | 23 | 100 | 156 | 170 | 1051 | 0.33 | 3 | 5 | 0.1 | -8 | 8 | 4 |
| | *Acer platanoides* | D3 | 30 | 108 | 142 | 200 | 631 | 0.25 | 5 | 4 | 0.025 | -17 | 10 | 4 |
| | *Acer pseudoplatanus* | D3 | 30 | 100 | 125 | 500 | 771 | 0.25 | 4 | 4 | 0.025 | -7 | 8 | 4 |
| | *Betula pendula* | D1 | 20 | 103 | 278 | 150 | 491 | 0.22 | 1 | 9 | 0.3 | -15 | 9 | 1 |
| | *Carpinus betulus* | D3 | 30 | 104 | 300 | 220 | 898 | 0.25 | 4 | 3 | 0.075 | -9 | 9 | 2 |
| | *Fraxinus excelsior* | D2 | 32 | 86 | 177 | 250 | 911 | 0.16 | 5 | 6 | 0.075 | -8 | 9 | 3 |
| | *Populus tremula* | D2 | 25 | 126 | 310 | 140 | 421 | 0.25 | 2 | 7 | 0.2 | -20 | 9 | 2 |
| | *Quercus pubescens* | D3 | 35 | 60 | 146 | 600 | 981 | 0.33 | 2 | 7 | 0.3 | -4 | 9 | 4 |
| | *Sorbus aria* | D2 | 22 | 66 | 82 | 180 | 650 | 0.33 | 4 | 7 | 0.2 | -20 | 12 | 4 |
| | *Sorbus aucuparia* | D1 | 20 | 107 | 167 | 110 | 491 | 0.33 | 3 | 7 | 0.2 | -15 | 7 | 4 |
| | *Ulmus glabra* | D3 | 43 | 127 | 153 | 480 | 631 | 0.25 | 5 | 3 | 0.075 | -8 | 9 | 3 |

Table C
ForCEEPS species-specific parameters derived from other species parameters.

| | Species | fS | NreqS | aS | f'S | f"S | cminS | cmaxS | n1S | n2S |
|---|---|---|---|---|---|---|---|---|---|---|
| | | | | *Derived from fS* | | | | | *Derived from NreqS* | |
| main species | *Abies alba* | E5 | 3 | 1.5 | 0.45 | 6 | 0.09 | 0.53 | -0.020 | 20 |
| | *Picea abies* | E5 | 2 | 1.5 | 0.45 | 6 | 0.09 | 0.53 | -0.019 | 10 |
| | *Pinus halepensis* | E4 | 1 | 1.4 | 0.45 | 6 | 0.071 | 0.346 | -0.019 | 2.5 |
| | *Pinus pinaster* | E4 | 1 | 1.4 | 0.45 | 6 | 0.071 | 0.346 | -0.019 | 2.5 |
| | *Pinus sylvestris* | E4 | 1 | 1.4 | 0.45 | 6 | 0.071 | 0.346 | -0.019 | 2.5 |
| | *Fagus sylvatica* | D3 | 2 | 1.7 | 0.35 | 12 | 0.025 | 0.1084 | -0.019 | 10 |
| | *Quercus ilex* | E4 | 1 | 1.4 | 0.45 | 6 | 0.071 | 0.346 | -0.019 | 2.5 |
| | *Quercus petraea* | D3 | 2 | 1.7 | 0.35 | 12 | 0.025 | 0.1084 | -0.019 | 10 |
| | *Quercus robur* | D3 | 2 | 1.7 | 0.35 | 12 | 0.025 | 0.1084 | -0.019 | 10 |
| other species | *Larix decidua* | D2 | 1 | 1.4 | 0.35 | 12 | 0.048 | 0.221 | -0.019 | 2.5 |
| | *Pinus cembra* | E5 | 1 | 1.5 | 0.45 | 6 | 0.09 | 0.53 | -0.019 | 2.5 |
| | *Acer campestre* | D2 | 3 | 1.4 | 0.35 | 12 | 0.048 | 0.221 | -0.020 | 20 |
| | *Acer platanoides* | D3 | 5 | 1.7 | 0.35 | 12 | 0.025 | 0.1084 | -0.024 | 50 |
| | *Acer pseudoplatanus* | D3 | 4 | 1.7 | 0.35 | 12 | 0.025 | 0.1084 | -0.022 | 35 |
| | *Betula pendula* | D1 | 1 | 1.4 | 0.35 | 12 | 0.038 | 0.1768 | -0.019 | 2.5 |
| | *Carpinus betulus* | D3 | 4 | 1.7 | 0.35 | 12 | 0.025 | 0.1084 | -0.022 | 35 |
| | *Fraxinus excelsior* | D2 | 5 | 1.4 | 0.35 | 12 | 0.048 | 0.221 | -0.024 | 50 |
| | *Populus tremula* | D2 | 2 | 1.4 | 0.35 | 12 | 0.048 | 0.221 | -0.019 | 10 |
| | *Quercus pubescens* | D3 | 2 | 1.7 | 0.35 | 12 | 0.025 | 0.1084 | -0.019 | 10 |
| | *Sorbus aria* | D2 | 4 | 1.4 | 0.35 | 12 | 0.048 | 0.221 | -0.022 | 35 |
| | *Sorbus aucuparia* | D1 | 3 | 1.4 | 0.35 | 12 | 0.038 | 0.1768 | -0.020 | 20 |
| | *Ulmus glabra* | D3 | 5 | 1.7 | 0.35 | 12 | 0.025 | 0.1084 | -0.024 | 50 |

Table D

Description of intrisic parameters in ForCEEPS. Parameters written in blue means that the values can be changed according to the simulation.

| Parameter | Value | Unit |
|---|---|---|
| *LCPmax* | 11.98 | - |
| *LCPmin* | 10.10 | - |
| $k_{death}$ | 4.605 | - |
| $p_{slowgr}$ | 0.368 | - |
| $t_{slowgr}$ | 0.03 | $cm.yr^{-1}$ |
| $k_{SlowGrTime}$ | 3 | yr |
| $k_{EstNr}$ | 0.006 | - |
| $k_{PatchSize}$ | 1000 | $m^2$ |
| $p_{dist}$ | 0.005 | - |
