## Appendix B for "Beyond forest succession: a gap model to study ecosystem functioning and tree community composition under climate change"

### Appendix B - Supplementary tables

Table S1. ForCEEPS parameters for the species used in this study. Parameter description, and literature used to calibrate all or part of the species for specific parameters can be found in Table 1.

| | **Species** | **$f_s$** | **$Hmax_s$** | **$S_s$** | **$G_s$** | **$Amax_s$** | **$DDmin_s$** | **$DrTol_s$** | **$Nreq_s$** | **$ShTol_s$** | **$ShTol\_seedling_s$** | **$Wtmin_s$** | **$Wtmax_s$** | **$Br_s$** |
|---|---|---|---|---|---|---|---|---|---|---|---|---|---|---|
| main species in the study | *Abies alba* | E5 | 50 | 75 | 350 | 366 | 841 | 0.23 | 3 | 1 | 0.05 | -6 | 5 | 5 |
| | *Picea abies* | E5 | 48 | 83 | 355 | 300 | 421 | 0.11 | 2 | 5 | 0.1 | -20 | 3 | 2 |
| | *Pinus halepensis* | E4 | 22 | 69 | 399 | 200 | 1261 | 0.48 | 1 | 7 | 0.25 | 2 | 13 | 3 |
| | *Pinus pinaster* | E4 | 35 | 69 | 350 | 300 | 1121 | 0.4 | 1 | 7 | 0.25 | -1 | 12 | 3 |
| | *Pinus sylvestris* | E4 | 35 | 58 | 150 | 200 | 631 | 0.37 | 1 | 9 | 0.3 | -20 | 5 | 3 |
| | *Fagus sylvatica* | D3 | 50 | 76 | 260 | 400 | 841 | 0.25 | 2 | 1 | 0.05 | -6 | 9 | 3 |
| | *Quercus ilex* | E4 | 23 | 48 | 79 | 500 | 1121 | 0.45 | 1 | 5 | 0.1 | 0 | 14 | 1 |
| | *Quercus petraea* | D3 | 45 | 76 | 246 | 1000 | 981 | 0.33 | 2 | 7 | 0.2 | -7 | 10 | 4 |
| | *Quercus robur* | D3 | 45 | 66 | 249 | 1000 | 631 | 0.3 | 2 | 9 | 0.3 | -10 | 10 | 4 |
| other species (for PNV simulaitons) | *Larix decidua* | D2 | 35 | 72 | 170 | 500 | 600 | 0.25 | 1 | 9 | 0.4 | -8 | -5 | 3 |
| | *Pinus cembra* | E5 | 25 | 40 | 115 | 1000 | 600 | 0.3 | 1 | 5 | 0.2 | -9 | -2 | 4 |
| | *Acer campestre* | D2 | 23 | 100 | 156 | 170 | 1051 | 0.33 | 3 | 5 | 0.1 | -8 | 8 | 4 |
| | *Acer platanoides* | D3 | 30 | 108 | 142 | 200 | 631 | 0.25 | 5 | 4 | 0.025 | -17 | 10 | 4 |
| | *Acer pseudoplatanus* | D3 | 30 | 100 | 125 | 500 | 771 | 0.25 | 4 | 4 | 0.025 | -7 | 8 | 4 |
| | *Betula pendula* | D1 | 20 | 103 | 278 | 150 | 491 | 0.22 | 1 | 9 | 0.3 | -15 | 9 | 1 |
| | *Carpinus betulus* | D3 | 30 | 104 | 300 | 220 | 898 | 0.25 | 4 | 3 | 0.075 | -9 | 9 | 2 |
| | *Fraxinus excelsior* | D2 | 32 | 86 | 177 | 250 | 911 | 0.16 | 5 | 6 | 0.075 | -8 | 9 | 3 |
| | *Populus tremula* | D2 | 25 | 126 | 310 | 140 | 421 | 0.25 | 2 | 7 | 0.2 | -20 | 9 | 2 |
| | *Quercus pubescens* | D3 | 35 | 60 | 146 | 600 | 981 | 0.33 | 2 | 7 | 0.3 | -4 | 9 | 4 |
| | *Sorbus aria* | D2 | 22 | 66 | 82 | 180 | 650 | 0.33 | 4 | 7 | 0.2 | -20 | 12 | 4 |
| | *Sorbus aucuparia* | D1 | 20 | 107 | 167 | 110 | 491 | 0.33 | 3 | 7 | 0.2 | -15 | 7 | 4 |
| | *Ulmus glabra* | D3 | 43 | 127 | 153 | 480 | 631 | 0.25 | 5 | 3 | 0.075 | -8 | 9 | 3 |

Table S2. Number of trees ($N_{trees}$) used for the calibration of $s_s$ and $g_s$ in the NFI data.

| **Main species** | **$N_{trees}$** |
|---|---|
| *Picea abies* | 24361 |
| *Fagus sylvatica* | 46766 |
| *Quercus robur* | 49381 |
| *Quercus petraea* | 44863 |
| *Abies alba* | 26009 |
| *Pinus pinaster* | 18285 |
| *Pinus sylvestris* | 29126 |
| *Quercus ilex* | 4981 |
| *Pinus halepensis* | 1519 |
| Total | 245291 |

| **Additional species** | **$N_{trees}$** |
|---|---|
| *Carpinus betulus* | 28465 |
| *Quercus pubescens* | 22813 |

Table S3. Description of 15 sites used for the PNV simulations, representing a gradient of environmental conditions. *MAT*: mean annual temperature; *ASP*: annual sum of precipitation; *DrI*: drought index calculated in ForCEEPS; *GDD*: growing degree-days.

| Site | Original site | MAT (°C) | mean ASP (mm) | DrI | mean annual GDD | Final basal area (m²) |
|---|---|---|---|---|---|---|
| site 1 | Alpine | 2.97 | 1577 | 0.11 | 855 | 24.6 |
| site 2 | HET26 | 6.97 | 1655 | 0.07 | 1550 | 28.3 |
| site 3 | EPC74 | 7.35 | 1450 | 0.00 | 1639 | 33.5 |
| site 4 | SAP26 | 8.09 | 1577 | 0.09 | 1799 | 33.8 |
| site 5 | SAP05 | 8.24 | 902 | 0.13 | 1850 | 31.4 |
| site 6 | SAP09 | 8.81 | 1252 | 0.15 | 1897 | 34.1 |
| site 7 | SAP39 | 9.99 | 1741 | 0.02 | 2283 | 36.1 |
| site 8 | HET88 | 10.21 | 1331 | 0.10 | 2314 | 32.8 |
| site 9 | PS78 | 11.04 | 701 | 0.21 | 2466 | 29.6 |
| site 10 | CHP10 | 11.36 | 800 | 0.15 | 2590 | 32.6 |
| site 11 | PS35 | 11.82 | 901 | 0.08 | 2590 | 32.2 |
| site 12 | HET64 | 13.02 | 1362 | 0.07 | 3065 | 33.9 |
| site 13 | PM40c | 13.38 | 890 | 0.20 | 3208 | 28.6 |
| site 14 | Puéchabon | 13.73 | 1018 | 0.30 | 3350 | 8.9 |
| site 15 | Font Blanche | 13.83 | 700 | 0.29 | 3369 | 8.4 |

Table S4. ForCEEPS accuracy in predicting tree growth across all species for repetition separately, through Pearson correlation, root mean square error (RMSE) and average bias (AB) between observed and predicted tree growth. Significance of the Pearson correlation coefficient (***: p < 0.001; **: p < 0.01; *: p < 0.05; *ns*: p > 0.05).

| | **Rep. 1** | **Rep. 2** | **Rep. 3** | **Rep. 4** | **Rep. 5** |
|---|---|---|---|---|---|
| Ntrees | 2662 | 2477 | 2416 | 2352 | 2295 |
| Pearson r | 0.0012 | 0.0012 | 0.0012 | 0.0012 | 0.0011 |
| RMSE | 0.72*** | 0.70*** | 0.70*** | 0.71*** | 0.71*** |
| AB | 0.124 | 0.1546 | 0.186 | 0.23 | 0.25 |

Table S5

Correlations between ForCEEPS parameters and ecological traits at the interspecific level. Upper table: Pearson's *r* ; lower table: associated p-value.

| | Wood density (g/m3) | LMA (g/m2) | Na (g/m2) | Amax (umol/m2/s) | P50 (Mpa) | Ψ_tlp (Mpa) | Ψclose (Mpa) | SM_Ψclose (Mpa) | SM_Ψtlp (Mpa) |
|---|---|---|---|---|---|---|---|---|---|
| **Pearson's R (All species)** | | | | | | | | | |
| $g_S$ | -0.574 | 0.557 | 0.249 | 0.408 | 0.297 | 0.203 | 0.173 | -0.257 | -0.284 |
| DrTol$_S$ | 0.290 | 0.407 | 0.339 | 0.538 | -0.605 | -0.287 | -0.262 | 0.577 | 0.607 |
| Nreq$_S$ | 0.143 | -0.531 | -0.501 | -0.666 | 0.048 | 0.359 | 0.161 | 0.023 | -0.018 |
| ShTol$_S$ | -0.073 | 0.136 | 0.365 | 0.719 | 0.011 | -0.264 | -0.150 | -0.098 | -0.055 |
| ShTol_seedling$_S$ | -0.094 | 0.189 | 0.306 | 0.869 | 0.024 | -0.362 | -0.217 | -0.145 | -0.083 |
| **Pearson's R (angiosperm)** | | | | | | | | | |
| $g_S$ | -0.431 | -0.174 | 0.000 | 0.509 | 0.646 | 0.134 | 0.077 | -0.745 | -0.707 |
| DrTol$_S$ | 0.621 | 0.582 | 0.451 | 0.191 | -0.838 | -0.645 | -0.581 | 0.728 | 0.811 |
| Nreq$_S$ | -0.171 | -0.508 | -0.430 | -0.478 | 0.125 | 0.432 | 0.258 | -0.019 | -0.079 |
| ShTol$_S$ | -0.110 | 0.263 | 0.346 | 0.712 | -0.002 | -0.167 | -0.059 | -0.036 | -0.019 |
| ShTol_seedling$_S$ | 0.024 | 0.149 | 0.121 | 0.875 | -0.001 | -0.371 | -0.239 | -0.128 | -0.061 |
| **Pearson's R (gymnosperm)** | | | | | | | | | |
| gS | -0.590 | 0.673 | 0.075 | -0.135 | -0.665 | 0.379 | 0.246 | 0.757 | 0.739 |
| DrTolS | 0.250 | 0.536 | 0.465 | 0.784 | -0.458 | 0.253 | 0.171 | 0.522 | 0.509 |
| NreqS | -0.598 | -0.219 | -0.811 | -0.937 | 0.050 | 0.545 | 0.540 | 0.235 | 0.100 |
| ShTolS | 0.218 | -0.047 | 0.667 | 0.863 | 0.006 | -0.589 | -0.543 | -0.290 | -0.158 |
| ShTol_seedlingS | 0.374 | -0.222 | 0.627 | 0.886 | -0.038 | -0.536 | -0.504 | -0.228 | -0.102 |

Pvalue All species

| | Wood density (g/m3) | LMA (g/m2) | Na (g/m2) | Amax (umol/m2/s) | P50 (Mpa) | Ψ_tlp (Mpa) | Ψclose (Mpa) | SM_Ψclose (Mpa) | SM_Ψtlp (Mpa) |
|---|---|---|---|---|---|---|---|---|---|
| gS | 0.005 | 0.011 | 0.392 | 0.188 | 0.180 | 0.391 | 0.455 | 0.261 | 0.212 |
| DrTolS | 0.191 | 0.075 | 0.236 | 0.071 | 0.003 | 0.220 | 0.251 | 0.006 | 0.004 |
| NreqS | 0.525 | 0.016 | 0.068 | 0.018 | 0.832 | 0.120 | 0.487 | 0.922 | 0.937 |
| ShTolS | 0.746 | 0.567 | 0.200 | 0.008 | 0.962 | 0.261 | 0.516 | 0.672 | 0.812 |
| ShTol_seedlingS | 0.679 | 0.424 | 0.288 | 0.000 | 0.917 | 0.117 | 0.345 | 0.530 | 0.720 |

Pvalue angiosperm

| | Wood density (g/m3) | LMA (g/m2) | Na (g/m2) | Amax (umol/m2/s) | P50 (Mpa) | Ψ_tlp (Mpa) | Ψclose (Mpa) | SM_Ψclose (Mpa) | SM_Ψtlp (Mpa) |
|---|---|---|---|---|---|---|---|---|---|
| gS | 0.108 | 0.553 | 0.999 | 0.198 | 0.009 | 0.663 | 0.794 | 0.002 | 0.005 |
| DrTolS | 0.013 | 0.029 | 0.190 | 0.650 | 0.000 | 0.017 | 0.029 | 0.003 | 0.000 |
| NreqS | 0.543 | 0.063 | 0.215 | 0.230 | 0.657 | 0.140 | 0.373 | 0.949 | 0.789 |
| ShTolS | 0.696 | 0.363 | 0.328 | 0.048 | 0.996 | 0.586 | 0.841 | 0.902 | 0.948 |
| ShTol_seedlingS | 0.933 | 0.611 | 0.740 | 0.004 | 0.996 | 0.212 | 0.411 | 0.662 | 0.835 |

Pvalue gymnosperm

| | Wood density (g/m3) | LMA (g/m2) | Na (g/m2) | Amax (umol/m2/s) | P50 (Mpa) | Ψ_tlp (Mpa) | Ψclose (Mpa) | SM_Ψclose (Mpa) | SM_Ψtlp (Mpa) |
|---|---|---|---|---|---|---|---|---|---|
| gS | 0.163 | 0.143 | 0.925 | 0.865 | 0.103 | 0.401 | 0.595 | 0.049 | 0.058 |
| DrTolS | 0.589 | 0.273 | 0.535 | 0.216 | 0.302 | 0.584 | 0.714 | 0.229 | 0.243 |
| NreqS | 0.156 | 0.676 | 0.189 | 0.063 | 0.916 | 0.206 | 0.211 | 0.612 | 0.830 |
| ShTolS | 0.638 | 0.930 | 0.333 | 0.137 | 0.989 | 0.164 | 0.207 | 0.528 | 0.735 |
| ShTol_seedlingS | 0.409 | 0.673 | 0.373 | 0.114 | 0.935 | 0.215 | 0.249 | 0.624 | 0.827 |
