## Appendix C for "Beyond forest succession: a gap model to study ecosystem functioning and tree community composition under climate change"

### Appendix C - Supplementary Figures

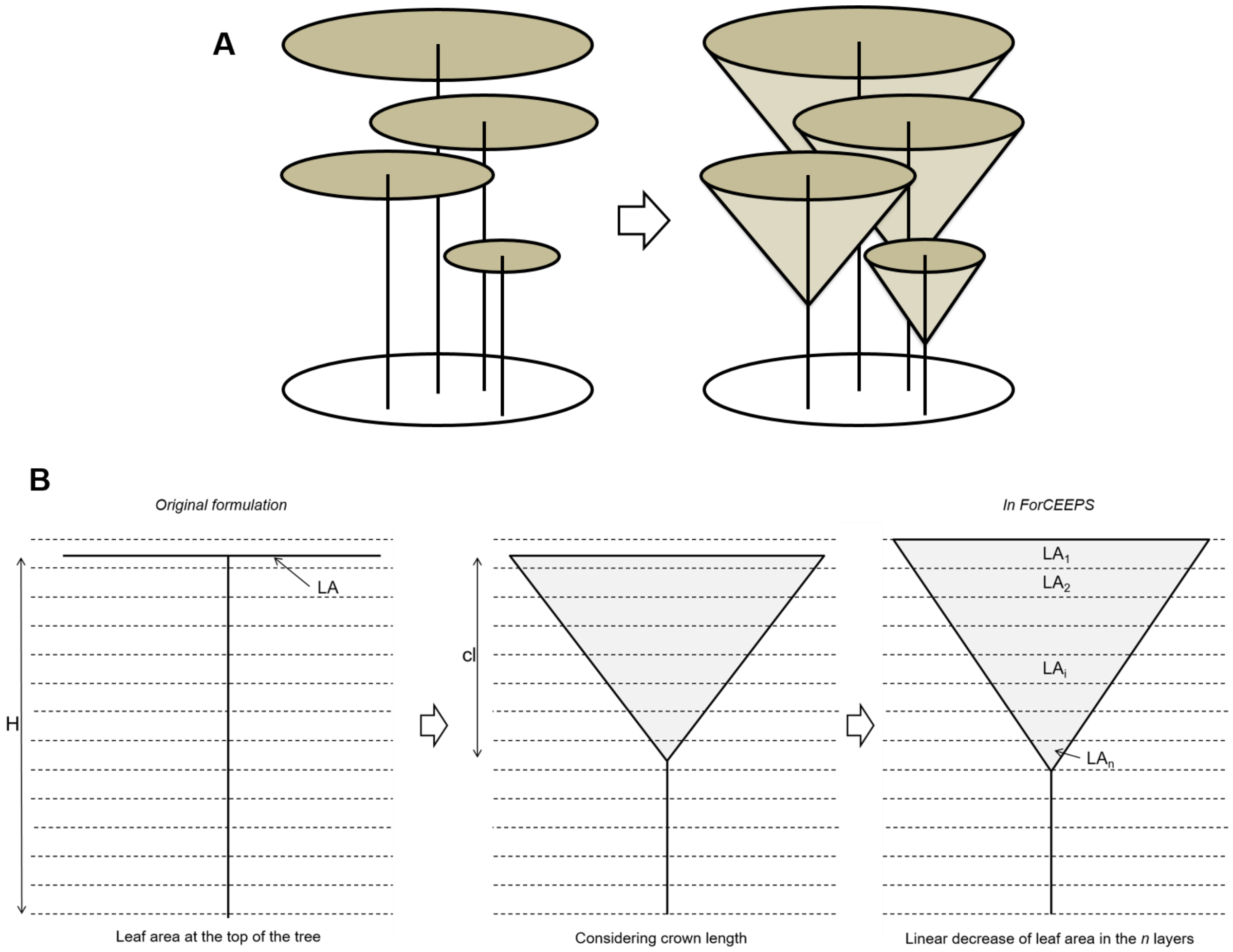


Figure S1. Representation of the foliage distribution at the tree level in ForCEEPS.

In ForClim 2.9.6, the total foliage of a tree is supposed to be concentrated at the top of the tree (on the left in panel A and "*original formulation*" in panel B). In ForCEEPS, the foliage is distributed in the vertical space along the crown length of the tree ("*cl*" on panel B and in main text), and tree leaf area was assumed to decrease linearly from the top to the base of the crown (see main text), through the discretization of the vertical space in layers (on the right in panel A and panel B).

Panel B: *H* is the total height of the tree.

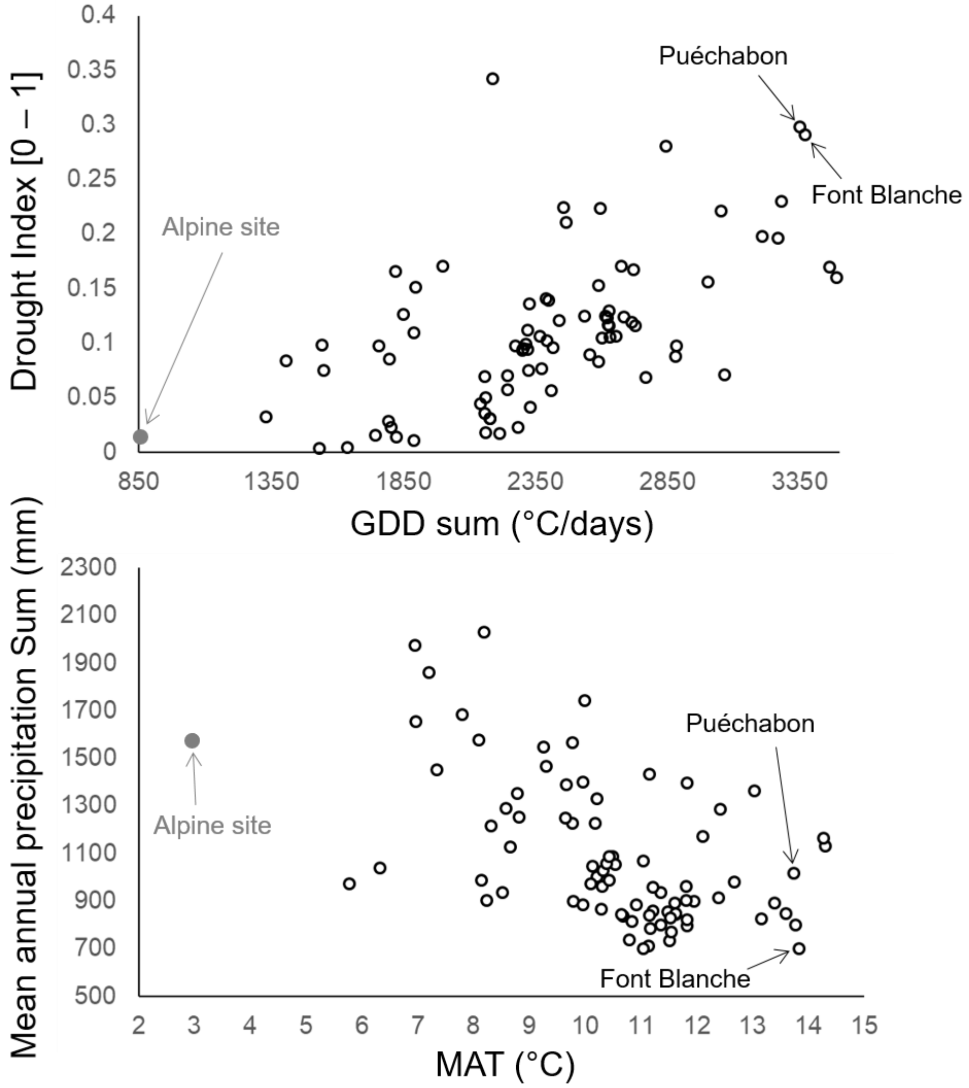


Figure S2. Distribution of the validation sites according to mean climatic variables.

Top panel: Mean drought index calculated in ForCEEPS [0-1] against mean annual growing degree days sums (with base temperature = 5 °C). Drought stress increases with the drought index. Low panel: Mean annual precipitation sum (mm) against mean annual temperature (°C). Mean climatic variables have been calculated across the time period of each dataset (see main text).

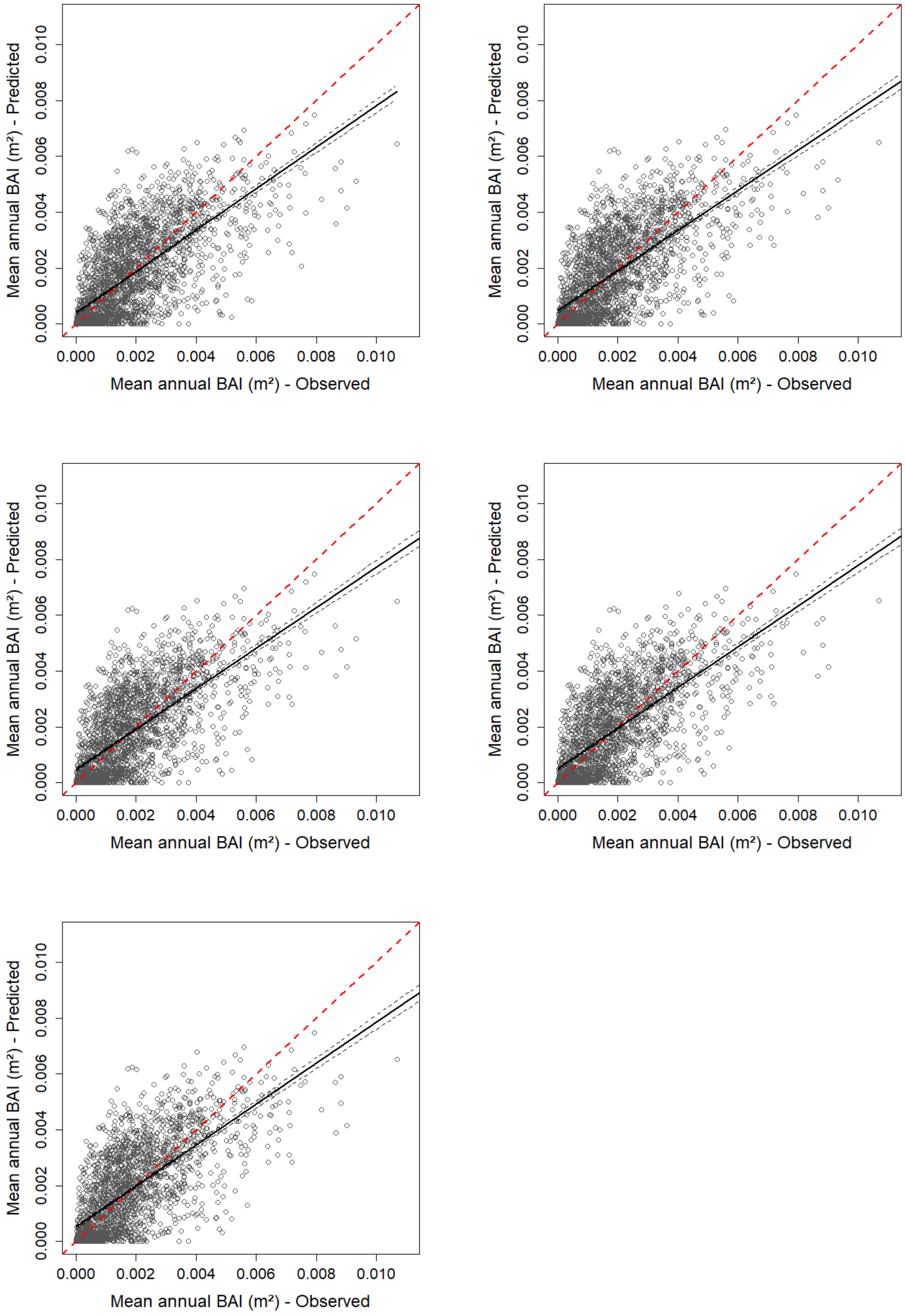


Figure S3. Predicted mean annual tree growth by ForCEEPS for all considered trees against observed mean annual tree growth for the five repetitions conducted. The plain black line is the regression line of the linear model of the relationship between observed and predicted tree growth, with confidence interval represented with the grey dashed lines; the dashed red line is the 1:1 line. Corresponding statistics in table S4.

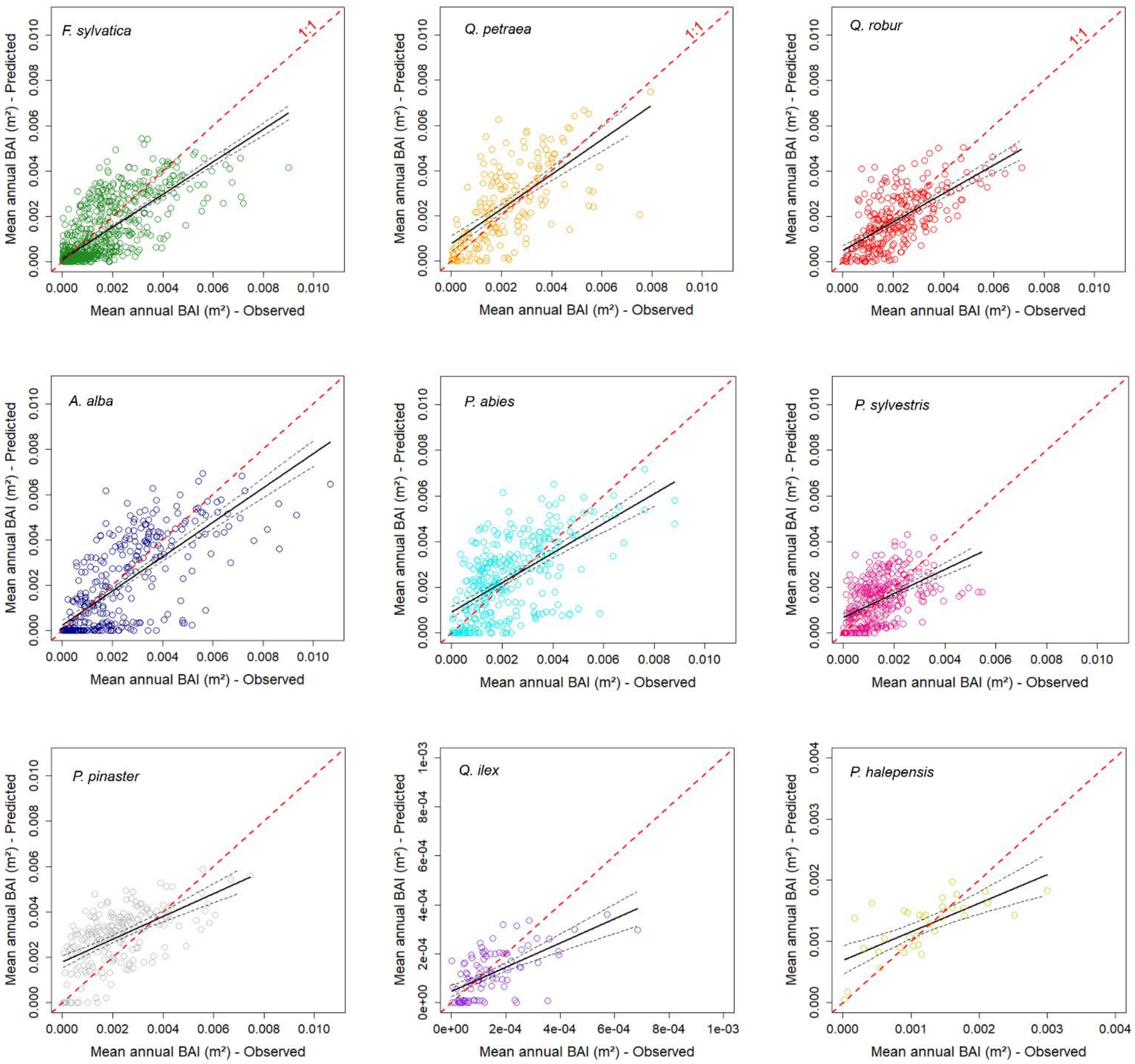


Figure S4. Predicted mean annual tree growth by ForCEEPS for all considered trees against observed mean annual tree growth for each species separately (only one repetition per species is shown). The plain black line is the regression line of the linear model of the relationship between observed and predicted stand productivity, with confidence interval represented with the grey dashed lines; the dashed red line is the 1:1 line. Corresponding statistics in table 2 (main text).

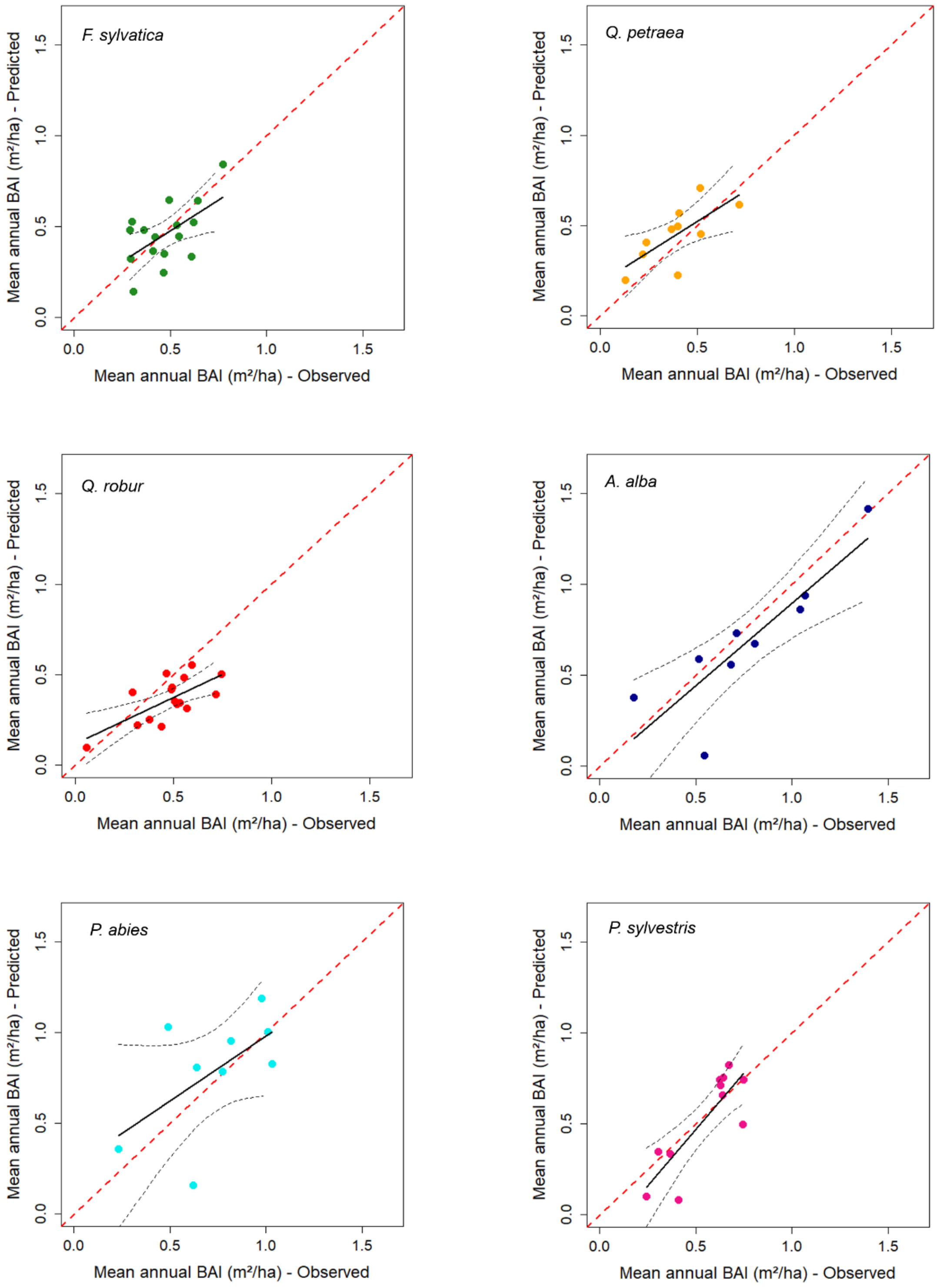
F. sylvatica
Q. petraea
Q. robur
A. alba
P. abies
P. sylvestris
Mean annual BAI (m²/ha) - Predicted
Mean annual BAI (m²/ha) - Observed
0.0
0.5
1.0
1.5

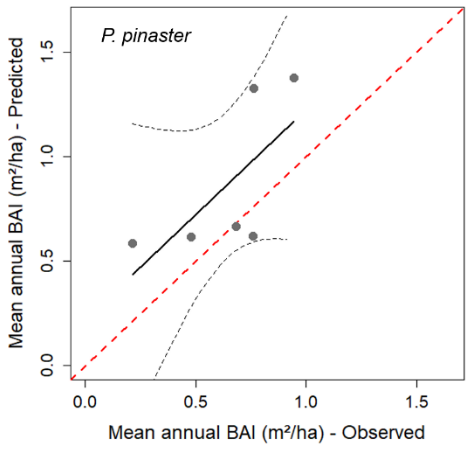


Figure S5. Predicted mean annual stand productivity by ForCEEPS against observed mean annual stand productivity for each species separately (averaged over all repetitions). The plain black line is the regression line of the linear model of the relationship between observed and predicted stand productivity, with confidence interval represented with the grey dashed lines; the dashed red line is the 1:1 line. Corresponding statistics in table 3-b (main text)

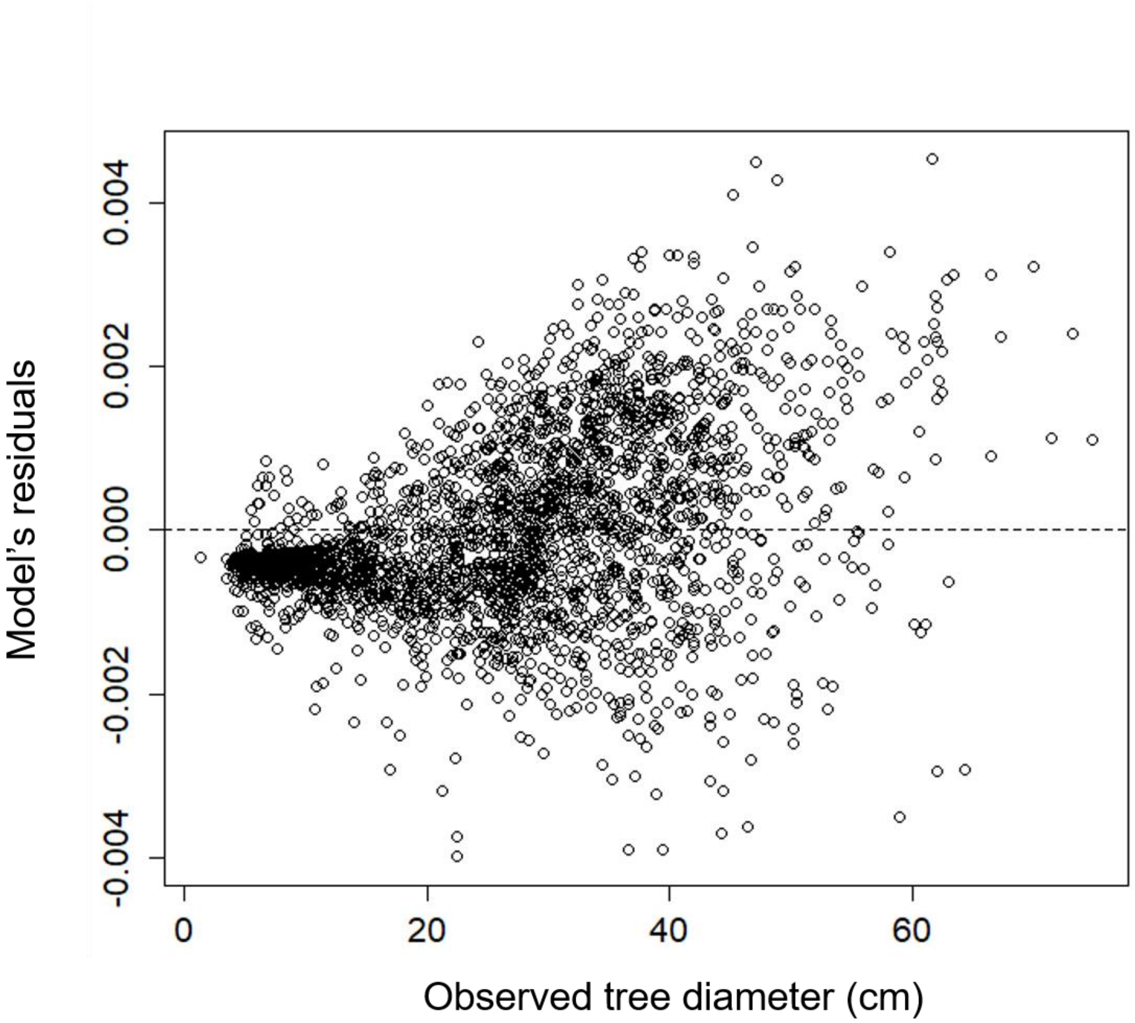


Figure S6. Residuals of the linear model *Predicted tree growth ~ Observed tree growth*, against observed tree diameter (for one repetition). Predicted tree growth simulated by ForCEEPS.

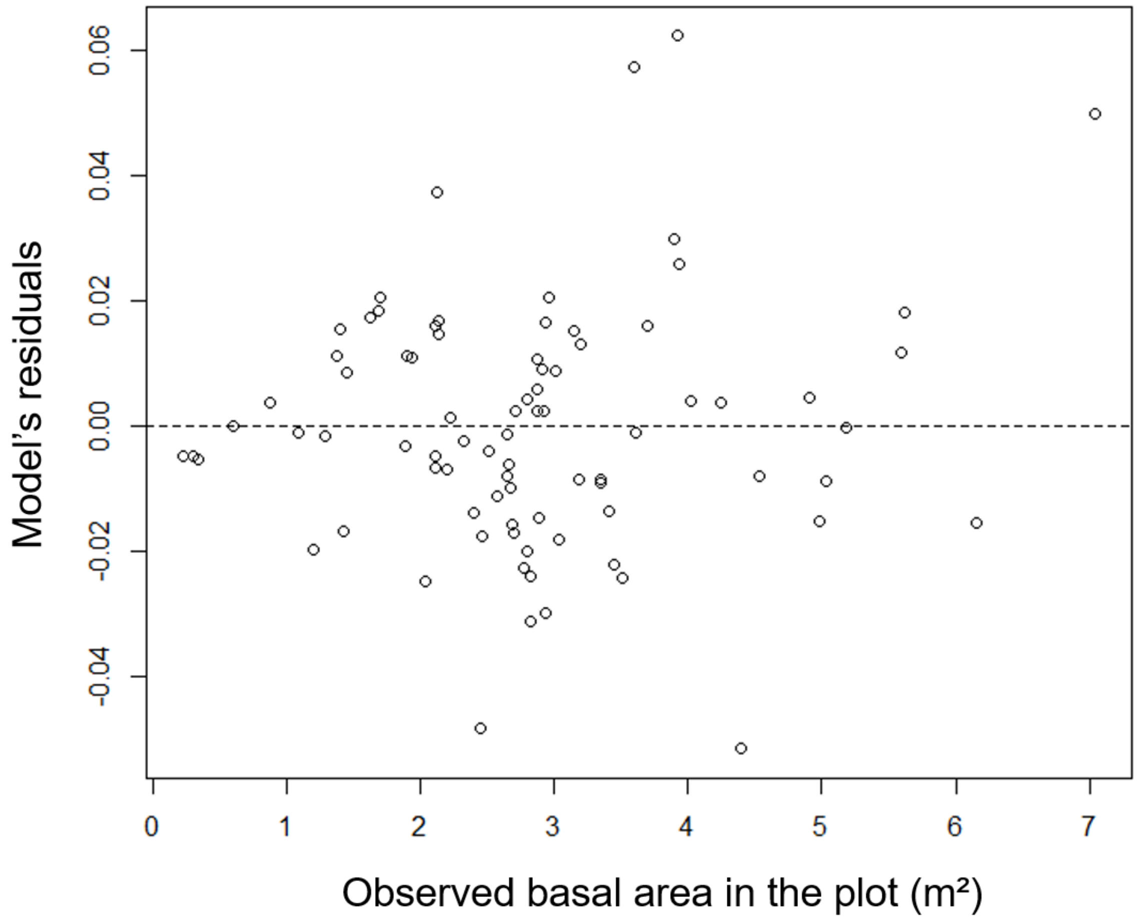


Figure S7. Residuals of the linear model *Predicted stand productivity ~ Observed stand productivity*, against observed basal area in the plot (averaged over all repetitions). Predicted stand productivity (in basal area) simulated by ForCEEPS.
