## Appendix D for "Beyond forest succession: a gap model to study ecosystem functioning and tree community composition under climate change"

### Appendix D – Additional Methods

#### Description of the validation sites and associated datasets

*RENECOFOR network*

The RENECOFOR permanent forest plot network (Ulrich 1997) includes 103 plots in even-aged managed forests covering most of the main tree species and climate conditions in mainland France. Most of the stands included in the RENECOFOR network are monospecific or strongly dominated by one species. The size of the RENECOFOR plots is *ca.* 0.5 ha. Diameter inventories over the 2000-2014 period were used to estimate the tree and stand basal area increment in all validation plots. The time interval between the initial and final inventories in RENECOFOR plots varied between 4 and 14 years. A complete description of the 103 stands can be found in (Ulrich 1997).

*Puéchabon*

The Puéchabon experimental site (43°44’29”N, 3°35’45”E, Southern France) is a Mediterranean evergreen coppice strongly dominated by *Quercus ilex* L. and unmanaged since the last clear cut performed in 1942. The mean canopy height is around 5.5 m and the stem density was 4900 stems ha$^{-1}$ in 2010. Understory species *Buxus sempervirens*, *Phyllirea latifolia*, *Pistacia terebinthus* and *Juniperus oxycedrus* compose a sparse layer with a percent cover under 25%, while rare individuals of *Quercus pubescens* and *Acer monspessulanum* may be present in the overstory. The rocky soil is formed on Jurassic limestone with an average rock volumetric fraction of 0.75 in the first 50 cm and 0.90 below, making a limited soil reserve of extractable water around 140 mm. The stone-free fine fraction of the topsoil (within 0–50 cm) is a homogeneous silty clay loam (USDA texture triangle; 38.8% clay, 35.2% silt and 26% sand). The site has a Mediterranean climate with about 80% of precipitation occurring between

September and April. The mean annual precipitation is 923 mm, and the mean annual temperature is 13.3°C, with a minimum in January (5.5°C) and a maximum in July (22.9°C). The experimental site has its own meteorological station located at 2 m height in a clearing and recording meteorological variables at a half-hourly time-step since 2001 with state-of the-art meteorological instruments compliant with the standards of the FluxNet network (https://fluxnet.fluxdata.org/about/). Biometric measurements in the Puéchabon site consisted of annual diameter surveys over the 2002–2017 period.

*Font Blanche*

The Font-blanche experimental site is a mixed Mediterranean forest located in south-eastern France (43°14’27’’N 5°40’45’’E, altitude 425 m a.s.l). An upper vegetation layer (average 13 m height) is dominated by *Pinus halepensis* whereas a lower tree strata is dominated by *Quercus ilex* (average 5m). A patchy understory is composed different tree and shrub species that do not exceed 4 m height: *Phillyrea latifolia*, *Quercus pubescens*, *Quercus coccifera*, *Arbutus unedo*, *Pistacia terebinthus*. The climate is Mediterranean, with an average annual temperature of 14°C and an average annual precipitation of 700 mm. The rocky substrate is formed by a Cretaceous rudist-bearing limestone with an average rock volumetric fraction of 0.5 in the first 40 cm and about 0.9 below. The soil reserve of extractable water for the ecosystem has been estimated at about 170 mm. The stone-free fine fraction of the topsoil (within 0–50 cm) is a homogeneous silty clay loam. Long term monitoring include meteorology (temperature, radiation, windspeed, rainfall, air humidity), forest – atmosphere CO2 and H2O exchanges (eddy-covariance tower), soil humidity, temperature, and respiration, and a number of biological variables, the most important in the context of this study being the annual tree stem circumference inventory. First measurements started in 2007.

**Field data for calibration.**

The parameterization of ForCEEPS was conducted using the French National Forest inventory dataset (NFI, http://inventaire-forestier.ign.fr/) compiled from 2005 to 2012. The NFI sampling method is based on temporary plots distributed over a systematic random grid of 1 km x 1 km over France (Charru et al. 2010, Vallet and Perot 2011). Biometric data are collected within concentric circles of 6, 9 and 15 m according to the size of the trees: all trees with a circumference greater than 23.5 cm, 70.5 cm, and 117.5 cm are measured in 6 m, 9 m, and 15 m circles, respectively. Biometric measurements include circumference at breast height, the last 5 years of annual radial increment (through radial cores) and total height. A relative weight is given to each measured tree, for upscaling at the stand level.
