## Appendix E for "Beyond forest succession: a gap model to study ecosystem functioning and tree community composition under climate change"

**Appendix E - Information about ecophysiological traits**

**Databases:**

SurEau database (Martin-StPaul et al. 2017)
https://zenodo.org/record/854700#.XnTwWkPjLmE

Cantrip (Keenan & Niinemets 2016)
https://github.com/trevorkeenan/traitPlasticity

Wood density : Zanne et al 2009, see description in Chave et al 2009

**References**

Chave, J., Coomes, D., Jansen, S., Lewis, S. L., Swenson, N. G., & Zanne, A. E. (2009). Towards a worldwide wood economics spectrum. Ecology Letters, 12(4), 351–366. doi: 10.1111/j.1461-0248.2009.01285.x

Keenan, T. F., & Niinemets, Ü. (2016). Global leaf trait estimates biased due to plasticity in the shade. *Nature Plants*, *3*(December), 1–6. doi: 10.1038/nplants.2016.201

Martin-StPaul, N., Delzon, S., & Cochard, H. (2017). Plant resistance to drought depends on timely stomatal closure. *Ecology Letters*, 1–23. doi: 10.1111/ele.12851

Zanne, A.E., Lopez-Gonzalez, G.*, Coomes, D.A., Ilic, J., Jansen, S., Lewis, S.L., Miller, R.B., Swenson, N.G., Wiemann, M.C., and Chave, J. 2009. Global wood density database. Dryad. Identifier: http://hdl.handle.net/10255/dryad.235

### **Traits values used**

(see main text for trait description)

| Species | LMA | Na | Aa | P50 | Psi_tlp | Psi_close | SM_Pclose | SM_Ptlp |
|---|---|---|---|---|---|---|---|---|
| Abies alba | 236.00 | 0.9 | 7.5 | -3.79 | -1.6 | -1.6 | 2.19 | 5.98 |
| Picea abies | 298.81 | 4.03 | | -3.61 | -2.5 | -2.5 | 1.11 | 4.72 |
| Pinus halepensis | 390 | | 15 | -5.14 | -2.2 | -2.35 | 2.79 | 7.93 |
| Pinus pinaster | 461.22 | 6.92 | 18.69 | -3.73 | -2 | -2 | 1.73 | 5.46 |
| Pinus sylvestris | 258.85 | 3.58 | 15.42 | -3.09 | -2.1 | -2 | 1.09 | 4.18 |
| Larix decidua | 117.99 | | | -3.81 | -2.67 | -2.67 | 1.14 | 4.95 |
| Fagus sylvatica | 90 | 1.90 | 7.22 | -3.15 | -2.5 | -2.5 | 0.65 | 3.8 |
| Quercus ilex | 213.25 | 3.45 | 8.22 | -6.9 | -3.15 | -3.18 | 3.73 | 10.63 |
| Quercus petraea | 126.31 | 3.13 | 12.50 | -3.3 | -2.7 | -2.8 | 0.5 | 3.8 |
| Quercus robur | 127.96 | 3.29 | 12.41 | -4.81 | -2.8 | -2.75 | 2.06 | 6.87 |
| Pinus cembra | | | | -3.3 | -2.34 | -2.34 | 0.96 | 4.26 |
| Acer campestre | 90.37 | | | -5.74 | -1.9 | -2.5 | 3.24 | 8.98 |
| Acer platanoides | | | | -4.18 | -1.6 | -1.6 | 2.58 | 6.76 |
| Acer pseudoplatanus | 50 | | 6.35 | -3.13 | -1.24 | -1.22 | 1.91 | 5.04 |
| Betula pendula | 83.40 | 2.64 | | -2.01 | -1.825 | -1.8625 | 0.1475 | 2.1575 |
| Carpinus betulus | 83.95 | | | -3.75 | -2.33 | -3.015 | 0.735 | 4.485 |
| Fraxinus excelsior | 126.68 | 2.27 | 7.22 | -2.8 | -2.1 | -2.33 | 0.47 | 3.27 |
| Populus tremula | 105.36 | 2.43 | 15.39 | -2.6 | -1.8 | -1.8 | 0.8 | 3.4 |
| Quercus pubescens | 97.5 | 1.87 | 14 | -4.8 | -2.63 | -2.63 | 2.17 | 6.97 |
| Sorbus aria | 98.08 | 2.24 | | -5.38 | -2.80 | -3.00 | 2.38 | 7.76 |
| Sorbus aucuparia | 92.85 | 2.22 | | -4.19 | | | | |
| Ulmus glabra | 44.6 | | | -3.5 | | -2.3 | 1.2 | 4.7 |
